# The complete genome of a songbird

**DOI:** 10.1101/2025.10.14.682431

**Authors:** Giulio Formenti, Nivesh Jain, Jack A. Medico, Marco Sollitto, Dmitry Antipov, Suziane Barcellos, Matthew Biegler, Inês Borges, J King Chang, Ying Chen, Haoyu Cheng, Helena Conceição, Matthew Davenport, Lorraine De Oliveira, Erick Duarte, Gillian Durham, Jonathan Fenn, Niamh Forde, Pedro A. Galante, Kenji Gerhardt, Alice M. Giani, Simona Giunta, Juhyun Kim, Aleksey Komissarov, Bonhwang Koo, Sergey Koren, Denis Larkin, Chul Lee, Heng Li, Kateryna Makova, Patrick Masterson, Terence Murphy, Kirsty McCaffrey, Rafael L.V. Mercuri, Yeojung Na, Mary J. O’Connell, Shujun Ou, Adam Phillippy, Marina Popova, Arang Rhie, Francisco J. Ruiz-Ruano, Simona Secomandi, Linnéa Smeds, Alexander Suh, Tatiana Tilley, Niki Vontzou, Paul D. Waters, Jennifer Balacco, Erich D. Jarvis

## Abstract

Bird genomes are the smallest among amniotes, but remain challenging to assemble due to their structural complexity. This study presents the first fully phased, diploid, telomere-to-telomere (T2T) reference genome for the zebra finch (*Taeniopygia guttata*), a model organism for neuroscience and evolutionary genomics. Combining multiple sequencing strategies resulted in closing nearly all gaps, adding ∼90 Mbp of previously missing sequence (7.8%). This includes T2T assemblies for all microchromosomes, including dot chromosomes, and the previously almost entirely missing chr16. The T2T genome is comprehensively annotated for genes, repeats, structural variants, and long-read methylation calls. Complete centromeric structures were assembled and annotated along with kinetochore binding sites. Relative to the previous high-quality reference of the Vertebrate Genomes Project, 2,778 (8.51%) previously unassembled or unannotated genes were identified, of which 9% overlap with segmental duplications. This first complete genome of a songbird, now the new public reference, illuminates avian genome architecture and function.

## Introduction

Birds possess the smallest genomes among amniotes (birds, reptiles, and mammals), typically ranging between 0.9 and 1.3 Gbp^1^, yet they show just as complex phenotypes^2^. Their sequencing and assembly presents unique challenges, particularly due to the presence of numerous small chromosomes, known as microchromosomes^3^. Microchromosomes are known to be gene-rich, GC-rich, with high recombination rates and short introns, and despite being generally considered highly repetitive, they show low densities of TEs and other repeats^1,4,5^. Bird microchromosome sequences are often underrepresented in genome assemblies due to substantial assembly challenges^6^, but contain many coding sequences and are generally highly conserved in avian evolution^7–9^.

The zebra finch (*Taeniopygia guttata*), a songbird native to Australia and the Lesser Sunda Islands, is a central model organism in neurobiology^10^, particularly for studying vocal learning, as well as in evolutionary genomics. Karyotype analyses show 40 chromosome pairs^11^, traditionally recognized as 7 macrochromosome pairs and 32 microchromosome pairs, plus the Z and W sex chromosomes^12^. In passerine birds, the Z chromosome is usually considered a macrochromosome, while the W chromosome is considered a microchromosome^13^.

The history of zebra finch genome assemblies reflects demands of the songbird field and major advances in sequencing technologies^14,15^. The first reference genome, TaeGut1.0 (Taeniopygia_guttata-3.2.4, GCA_000151805.2), published in 2010, was the second bird genome sequenced after chicken, and generated using Sanger sequencing of BAC clones (∼5.5× coverage) and a genetic linkage map^16,17^. This assembly was highly fragmented (contig N50 = 39 Kbp; scaffold N50 = 10 Mbp), capturing only a subset of macrochromosomes and lacking much of the microchromosomal content. In 2021, the Vertebrate Genomes Project (VGP) released bTaeGut1, a major improvement produced from the same male individual (Black17) using PacBio Continuous Long Reads (CLR), 10x Genomics linked reads, Bionano optical maps, Hi-C chromosomal linked reads, and parental Illumina reads to enable haplotype phasing^3^. This improvement also involved the generation of new algorithms, particularly ones that phased haplotypes and scaffolded contigs (continuous gapless sequences) into chromosomes, and was one of the genomes in the VGP flagship study^3^. bTaeGut1 corrected thousands of structural mis-joins (14,381), resolved nine inversions, closed tens of thousands of gaps (estimated 58,031), added 49.4–68.5 Mb of missing sequence, identified seven additional chromosomes (30–36), and detected haplotype-specific structural variants on chromosomes 5, 11, and 13. Telomeric motifs recovered on some scaffolds suggested increased completeness. Version bTaeGut1.4.pri became the official reference genome in 2021 (GCA_003957565.4). Alongside this, a female genome from a different laboratory population (Blue55, ToLID bTaeGut2/E055JL) was also sequenced using the same approach, with additional Illumina data from the parents to generate phased haplotypes^18 18^GCA_008822105.2). This assembly partially recovered the W chromosome. In 2024, the VGP generated an updated phased diploid assembly of Blue55/bTaeGut2 (GCA_051427915.1, GCA_051428105.1) using more accurate PacBio HiFi reads, Hi-C, and the latest VGP pipeline^15^, further improving assembly continuity and accuracy, but this was not initially released in NCBI and did not become a reference.

The first complete telomere-to-telomere (T2T) genome of a vertebrate was a human in 2022^19^, which was derived from a haploid cell line, making assembly easier, since there were no haplotypes to phase. Thereafter, diploid assembly protocols were developed that have achieved near T2T completeness in humans^20–23^. In addition, gapless and near-error free assemblies now exist for some non-model species, though usually only one haplotype is presented at T2T^24–27^. The only other avian T2T genome available to date is that of the chicken^28^, though the assembly was not fully phased and has 15 gaps in chrW. In the study, the authors defined dot chromosomes as among the smallest of the microchromosomes.

In the current study, multiple sequencing data sets were generated for a female individual (bTaeGut7/Red 201; the heterogametic sex with both sex chromosomes) and its parents (bTaeGut8/Red 52 mother; bTaeGut9/Green 47 father) to assemble the first fully phased, manually curated, extensively annotated, T2T diploid reference genome for a bird, the zebra finch. This assembly resolves all autosomes and sex chromosomes, including dot chromosomes. The centromeres and telomeres are all present and complete, and a model sequence for rDNA arrays was added, ultimately representing a comprehensive and gapless representation of the somatic genome. NCBI has now made it the public reference genome for the zebra finch. Given the ∼100 million years of divergence between the chicken and zebra finch lineages^17,29^, this resource provides a critical reference for comparative and functional genomics across birds. The analysis of this new reference precisely reveals the structural organization of chromosomes in the zebra finch and suggests a model of chromosomal evolution common to birds.

## Results

### A near-complete diploid reference genome of the zebra finch

The new T2T zebra finch reference genome was generated using a combination of PacBio HiFi (coverage ∼122x), Oxford Nanopore Technology (ONT) R10 ligation (∼117x) and ultra-long reads (∼255x, of which ∼29x >100 Kbp), Hi-C long range information (∼57x) and parental Illumina short reads for phasing (**Figure 1A**; **Supplementary Figure 1, Supplementary Table 1**). Sequencing ONT reads required a specialized protocol developed at the Vertebrate Genome Laboratory (VGL), with multiple rounds of pore re-opening and cleaning steps, due presumably to the complex structure of avian genomes. A hybrid assembly strategy was employed, in which multiple assemblies were generated using different combinations of sequencing datasets, assemblers (mainly Verkko^20^ and some contribution from hifiasm for residual gaps^30^), and tool parameters. For example, ONT-corrected reads, when used to generate the initial contig layout of the assembly graph, filled in gaps present in contigs generated only from HiFi reads, particularly in microchromosomes. No single assembly run produced a complete and gapless reconstruction of all chromosomes; thus, all chromosomes were manually reviewed and curated via manual graph resolution, also assigned by gfalign, a newly developed tool developed to aid and streamline T2T genome assembly (**Supplementary information**). The best representation for each of the 40 chromosomes for each haplotype was selected for inclusion in the final reference genome (**Figure 1B**). These can be classified into 7 macrochromosomes and 32 microchromosomes (of which 11 of the latter are dot chromosomes), plus the Z and W chromosomes.

**Figure 1.**
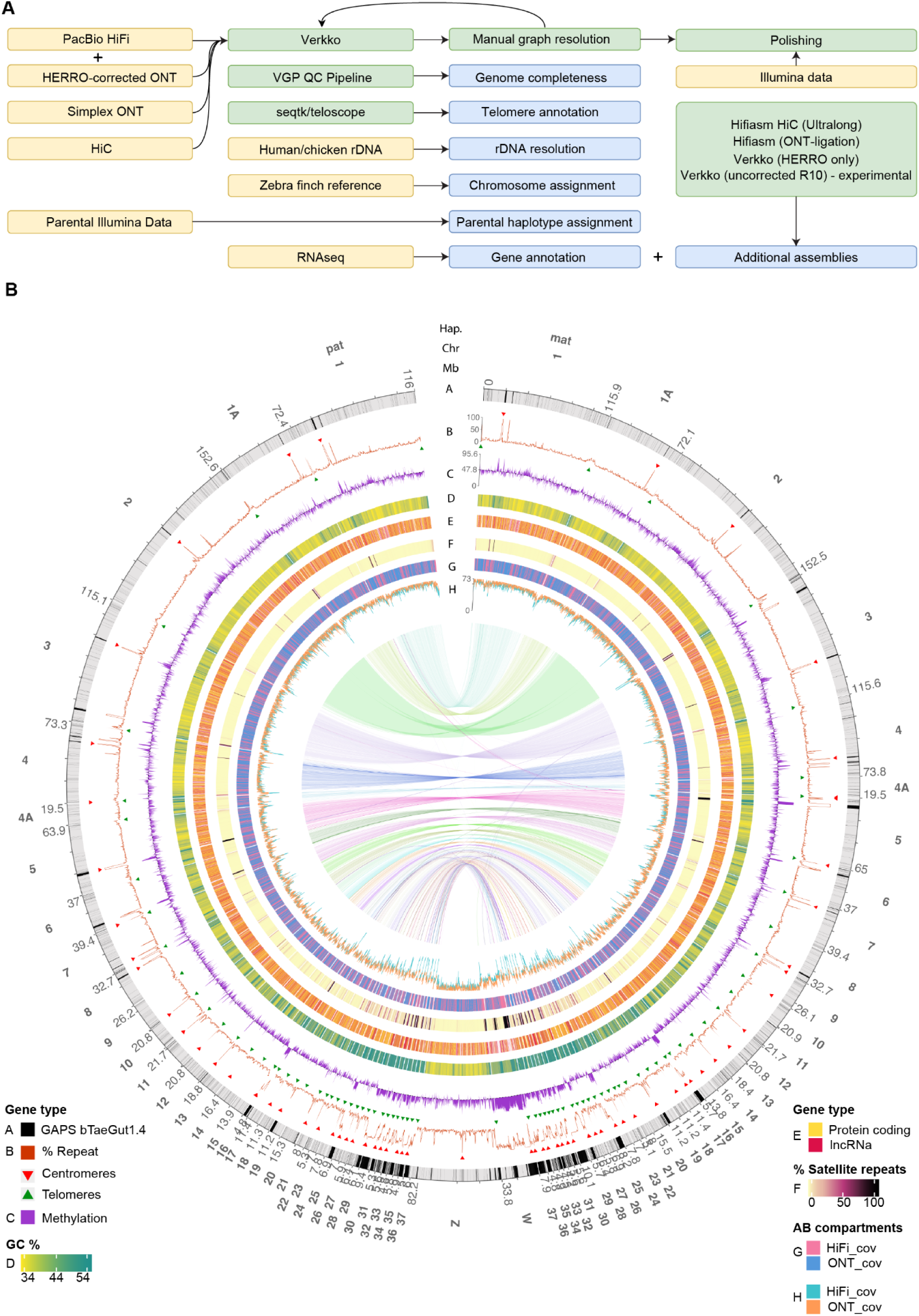
Assembly of a complete zebra finch genome. (A) Final assembly strategy used for the bTaeGut7 reference diploid genome. (B) Diploid Circos plot, illustrating the gaps in the previous bTaeGut4 reference (track A), repeat distribution with telomere and centromere positions (track B), minisatellite repeats distribution (track C), GC content (track D), methylation levels (track E), genes distribution (track F, coding genes and long non-coding RNA coloured differently), and HiFi and ONT coverage distribution (track G). Synteny between the paternal (pat) and maternal (mat) haplotypes is shown as links in the center of the plot.

The resulting assembly spans 1.14 Gbp for the maternal haplotype plus the Z chromosome from the paternal haplotype, which was designated as the main primary reference (**Supplementary Table 2**). The paternal autosomes span 1.02 Gbp. Telomere sequences are present on both ends of all chromosomes (**Figure 2A**), parental haplotypes are fully phased (**Supplementary Figure 2**), all centromeres resolved, and both sex chromosomes and the mitogenome are represented (**Supplementary Figure 3**). Estimated per-base accuracy is Q55.1 (∼3 nucleotide errors per Mbp), measured from hybrid 31-mers of Illumina and HiFi reads (**Supplementary Figure 2, Supplementary Tables 3-5**). The real relative chromosome sizes largely do not match the relative chromosome numbers that were previously assigned^4,17^ (**Supplementary Figure 4**). Zebra finch macrochromosomes were originally assigned based on homology with chicken chromosomes (which were named according to size), and zebra finch microchromosomes were originally named by relative size. There were only 5 sequence gaps remaining in the assembly, 4 of which are on the Z chromosome (the 4th largest chromosome, 82.2 Mbp, **Supplementary Figure 5**) and 1 in the rDNA tangles on chr37 (**Supplementary Figure 6**), similarly to the human T2T assembly^19^. Sequence models were used to provide a representation of several of these residual tangles, filling in 2 of the gaps on the Z chromosome and the haplotype-aware rDNA models for chr37. Despite the high repeat content (83%), the W chromosome is completely resolved with no gaps. The reference has an estimate of only 0.013% of hamming haplotype switch error rate (**Supplementary Figure 2**). Extensive annotation tracks have been generated and are available in the UCSC Genome Browser (**Supplementary Figure 7**).

**Figure 2.**
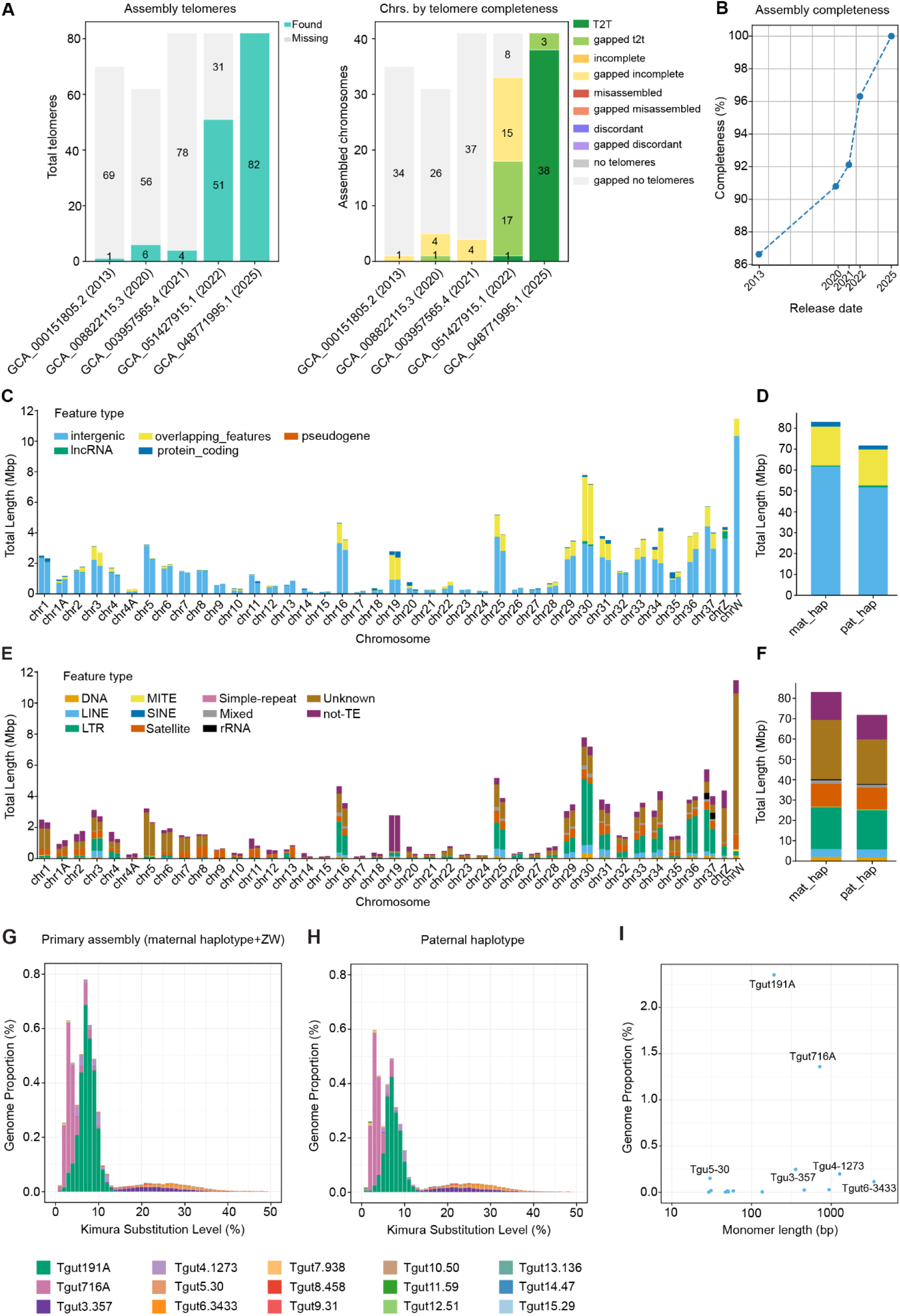
Improvements over bTaeGut1.4 and other previous references. (A) Telomere and gap completeness across zebra finch references (**Supplementary Table 6**). Displayed are improvements in total telomeres found and chromosomes by telomere and gap completeness. (B) Progression of zebra finch reference genome completeness over time relative to the T2T assembly. (C) Distribution of genomic features within previously unassembled regions across chromosomes, including of each haplotype, annotated as protein-coding genes, lncRNAs, intergenic regions, overlapping features (presenting 2 or more overlapping features), and pseudogenes. (D) Total number of bases inside previously unassembled regions and the same annotated features. (E,F) Stacked bar plot showing the composition of repeat classes annotated within previously unassembled regions, and enrichment plot indicating the relative abundance of repeat on each chromosome. (G,H) Distribution of abundance and divergence of satDNA families in the maternal and paternal haplotypes, respectively. Divergence is calculated as the Kimura two-parameter distance to a consensus sequence. (I) Satellite monomer length in relation to its proportion in the genome.

*Improvements over bTaeGut1.4*

Relative to the previous long-read based VGP bTaeGut1.4 reference, the new bTaeGut7 reference incorporates 89.9 Mbp of additional sequence (7.8%), including pericentromeric and subtelomeric sequences (**Figure 2B, Supplementary Table 6**). Furthermore, 49.4 Mb of the newly assembled regions overlap with segmental duplication blocks. The W chromosome the (10th largest chromosome, 33.8 Mbp) contains 11.5 Mbp (34.1%) of previously unassembled sequence. The genic composition of the newly assembled regions varies substantially between chromosomes and even between maternal and paternal haplotypes (**Figure 2C,D**). The newly assembled regions are predominantly intergenic, but do also contain protein-coding genes, particularly enriched on chr16, 18, and 20, and a notable presence of lncRNAs, particularly on chrZ. Pseudogenes are generally underrepresented and not significantly enriched in the newly assembled regions of most chromosomes, with the exception of maternal chr28. A total of 2,788 newly assembled genes were identified across the maternal haplotype and the Z chromosome. Focusing on the maternal haplotype autosomes (2,682 genes), 2,526 were entirely contained within the newly assembled regions, while 156 partially overlapped with previously assembled regions, covering at least 75% of their length. These comprised 1,009 protein-coding genes, 1,564 long non-coding RNAs, and 109 pseudogenes. On the paternal haplotype autosomes, 2,645 genes were annotated within previously unassembled regions, of which 2,491 were fully contained and 154 partially overlapped (≥75% of gene length). This set included 929 protein-coding genes, 1,610 lncRNAs, and 106 pseudogenes. Hence, many previous missing protein-coding genes identified were absent or partially annotated in the previous assembly. This includes the synapsin 1 gene (SYN1) important for brain function, previously thought to be absent in zebra finch^17^, which was identified on chr31 in the bTaeGut7 assembly, following the recovery of a genomic region missing from earlier assemblies. Other examples of previously missing genes are PPP5C, CLIP3, CAPNS1, DMWD, PPP1R18, CSNK2B, and IFT7.

### Repetitive regions are now fully assembled

Previous unassembled regions also contain many repetitive elements (**Figure 2E,F**). SINEs were among the most underrepresented elements in previously unassembled regions together with MITE and “Simple repeats” elements. LINEs were enriched in e.g. chr25, chr30 and chr31, and LTR elements were also enriched in previous unassembled regions located in chr16, chr25, chr30 (around 4 Mbp), chr36, and chr37. TEs overrepresentation in previous unassembled regions suggests that these regions may harbor functionally relevant regulatory elements. Satellite portions in previously unassembled regions are consistently large, including very high loads on sex chromosomes (900 kbp on chrW and 202 kbp on chrZ). Particularly, centromeric satellite sequences were almost entirely missing for all chromosomes in bTaeGut1.4 and are now fully assembled in bTaeGut7. Indeed, satellites of various kinds are abundant in bTaeGut7 (**Supplementary Figure 8**), with varying degrees of conservation (**Figure 2G,H**) and monomer length (**Figure 2I**).

### Resolution of the rDNA

Ribosomal 5S rDNA arrays were 84 kbp and 245 kbp, comprising two high-GC (>65%) clusters located on the medial region of chromosome 2q, separated by ∼680 kbp with two coding genes (TMX, DSEL) and multiple non-coding RNAs. The 5S monomer is ∼378 bp, and it was previously characterized as a highly frequent satellite sequence labelled Tgut378A^31^. The two 5S arrays were almost completely missing in the previous reference and are fully resolved in both bTaeGut7 haplotypes. 18/28S rDNA probes were previously shown to hybridize onto a single pair of microchromosomes^13^. In the T2T assembly, the chromosome pair with the 18/28S rDNA is chr37, named based on homology with bTaeGut1.4. With all dot chromosomes resolved, chr37 is the 27^th^ largest chromosome, but a decision was made to keep numbering consistent with the past literature. Like in humans^19^, the full rDNA cluster was too similar and repetitive to be resolved with any T2T assembly pipeline. Thus, a model sequence derived from a random mixture of rDNA morphs found in each haplotype^32^ was used to fill in the 18S/28S rDNA array on chr37 (**Supplementary information**). The two rDNA clusters varied 2-fold in size between haplotypes (1.4 Mbp vs 0.72 Mbp for maternal and paternal, respectively), reminiscent of the striking variability seen in humans.^33^

### Centromere characterization

The availability of fully resolved centromeres in the new diploid reference genome enables, for the first time, a detailed investigation into the nature and organization of zebra finch centromeres. Avian centromeres have often been considered primarily epigenetically determined^34^, characterized by CENP-A histone accumulation, yet a number of centromere-specific sequences have also been identified^35^. Most avian centromere-specific monomers range from 101 to 593 bp in length^31^. In chicken, the 41 bp CNM (chicken erythrocyte nuclear membrane^36^) is a well-characterized centromeric repeat, and additional elements such as PO41 and TM also function as centromeres in Galliformes^28,37^. However, little is known about centromeric composition in Passeriformes, with centromeric repeats that appear to have been previously studied only in the chaffinch (*Fringilla coelebs*)^38^. In the zebra finch, two satellites, Tgut191A (191 bp) and Tgut716A (716 bp), have been proposed as pericentromeric^31^. Based on quantification from short-read data, these repeats comprise approximately 5% of the genome, with Tgut191A predicted to contribute ∼57 Mbp (∼300,000 copies) and Tgut716A ∼52.6 Mbp (∼70,000 copies)^31^. However, due to the difficulties of assembling satellite DNA even with PacBio CLR data used for bTaeGut1.4, that assembly contains only ∼1,500 and ∼300 copies of Tgut191A and Tgut716A, respectively^31^. In the T2T assembly, ∼60K copies of Tgut191A were annotated on the autosomes (59,913 and 61,485 in the maternal and paternal haplotypes, respectively), plus another 47,133 copies on the W chromosome, and 9,081 on the Z chromosome (**Figure 3A**). Tgut716A is present in 13,407 and 13,296 copies in the maternal and paternal autosomes, as well as in 1,598 and 708 on the W and Z chromosomes, respectively. No significant coverage issues were observed across the centromere satellites, suggesting that the difference with the estimates from raw sequencing data are likely explained by the combination of the well-known short-read sequencing bias^39^ and individual haplotype variation in satellite repeats^40^.

**Figure 3.**
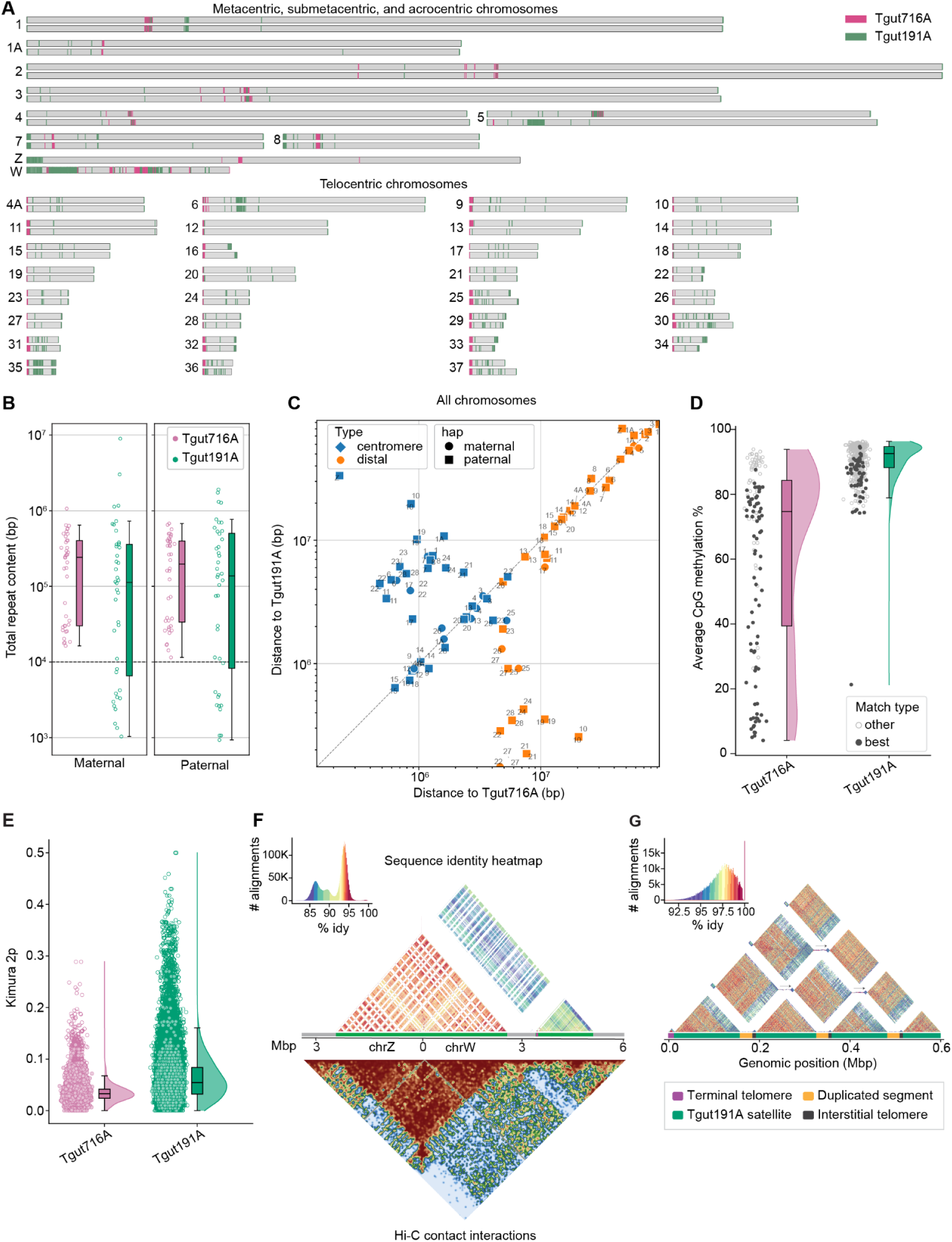
Characterization of centromeric repeats. (A) Overview of centromeric satellite localizations. (B) Tgut716A and Tgut191A presence and length in all chromosomes. (C) Colocalization of putative centromeres with cytogenetic maps. The axes represent the distance from Tgut716A and Tgut191A of centromeric and distal PCR primers from a previous study^41^. (D) Methylation colocalization. Points show the average methylation for individual repeat arrays. One “best”, i.e. with the lowest methylation average is also highlighted for each chromosome. (E) Kimura 2-parameter (K2P) distances of Tgut716A and Tgut191A with respect to their chromosome consensus. The filled data points represent repeats within the centromere (Tgut716A) and PAR (Tgut191A). (F) Side by side comparison of PAR regions in chrZ and chrW. Image shows an annotated sequence identity heatmap (top) and the corresponding location in the Hi-C map (bottom), using StainedGlass^58^ and PretextView, respectively. Tgut716A, Tgut191A, and unique sequences are indicated using different colors. (G) Sequence identity heatmaps of 600 kbp in the p-arm of the maternal chr35 containing multiple ITS tandem arrays interleaved with duplicated segments and Tgut191A repeats.

No other satellite repeats were ubiquitously present as Tgut191A and Tgut716A to be candidate centromeric repeats. Tgut716A emerged as the strongest candidate for a centromeric function, directly involved in kinetochore binding. First, on all chromosomes there was at least one candidate large Tgut716A array (largest array 677 kbp on chr13 paternal, smallest 11.5 kbp on chr19 paternal), whereas Tgut191A was often more limited, less than 10 kbp/chromosome in total (**Figure 3B**). Second, Tgut716A was always (100%) found close to centromeric locations in cytogenetic maps derived from multiple experiments^17,41^, whereas Tgut191A usually was present on the opposite side in acrocentric chromosomes (**Figure 3C**). Third, Tgut716A exhibited at least one clear methylation dip per chromosome (a characteristic of centromeres in the human T2T assembly^42^), while Tgut191A remained constitutively hypermethylated (**Figure 3D**). Fourth, in microchromosomes, Tgut716A was always (100%) flanking heterochromatin, especially in dot chromosomes, whereas Tgut191A was usually found next to the euchromatic regions. Heterochromatin flanking centromeres is believed to play an important role in stabilizing centromeres, promoting proper attachment of chromosomes to the mitotic and meiotic spindles by helping recruiting and retaining cohesin^43^. The Tgut716A monomer is also significantly less divergent than that of Tgut191A (**Figure 3E**). Finally, the candidate centromeric positions for the macrochromosomes identified using Tgut716A array position, conservation, and methylation substantially coincide with the primary constriction sites inferred from the karyotype (**Supplementary Figure 9**).

Earlier investigations did not detect a CENP-B box in these repeats^31^. This was consistent with the belief that the CENP-B locus is absent in non-mammalian species^44,45^. In the T2T zebra finch assembly, while Tgut191A showed no homology with known CENP-B box motifs, Tgut716A contained motifs with high sequence similarity with the human CENP-B box (**Supplementary Figure 10**). In addition, BLAST searches for CENP-B homologs in the zebra finch T2T genome identified TIGD4 (Tigger Transposable Element Derived 4) as the top candidate, although divergent (57% query coverage and 28.78% sequence identity). TIGD4, like CENP-B, is a Group I pogoR-derived transposase. Also similar to CENP-B, AlphaFold3 modeling predicts TIGD4 potential to form dimers and directly bind to CENP-C and to the Tgut716A satellite monomer (pTM = 0.33), with the DNA-binding domain localized to residues 215-430 (**Supplementary Figure 11, Supplementary Information**). TIGD4 is located in a syntenic block conserved across birds, amphibians, and mammals (chr4, ∼42-44 Mbp), surrounded by syntenic genes like ARFIP1 and TMEM154 (**Supplementary Figure 12**). TIGD4 is significantly more highly expressed in testis than in brain, with a log₂ fold change (MLE; brain vs testis) of –2.49526 (Wald test-value=9.72×10⁻⁶⁹; padj= 3.02×10⁻⁶⁸; mean= 130.444; lfcSE= 0.1424; Wald statistic = –17.5221), a classical hallmark of centromeric proteins^46^ (**Supplementary Figure 13**). The identification of both a candidate CENP-B box in Tgut716A and TIGD4 open to the possibility of a complete CENP-like system in birds, previously thought to be a prerogative of mammals.

The sequence of Tgut716A shows no similarity to centromeric satellites identified in chicken^28^. In RepBase^47^, it exhibits only limited homology (443 bp, 69% identity) with crowSat1, a satellite suspected to be a major heterochromatin component in crows^48^. In contrast, another study^31^ noted some satellite repeats in multiple other Passeriformes closely related to the zebra finch that show high similarity to Tgut716A (76–85%) and to Tgut191A (83–91%), supporting the hypothesis that these repeats are common within this lineage and may play similar functional roles. These prior findings are consistent with the view that centromeric satellite sequences show little evidence of conservation beyond ∼50MY of divergence across eukaryotes^49^ (Passeriformes origin ∼35MY and Passeriformes/Chicken divergence ∼90MY^29^). Allowing to investigate these ideas further, the T2T assembly clearly shows that when Tgut716A is found at chromosome termini, it lies directly adjacent to telomeric repeats, supporting the notion that all microchromosomes in the zebra finch are telocentric, rather than acrocentric^13^. While the 191 bp unit length of Tgut191A is consistent with mononucleosome organization, the association of Tgut716A with centromere binding implies that zebra finch centromeres may adopt a tetranucleosome structure. Interestingly, the presence of so many copies of Tgut191A on the W chromosome, making up 26% of its sequence, suggests a distinct role or evolutionary trajectory for this satellite repeat in shaping the architecture of sex chromosomes. In support of this idea, the pseudoautosomal regions (PAR) in the W chromosome appears to be located in the first ∼3 Mbp of the p arm, flanked by a relatively short centromere (51 kbp), with cytogenetic studies supporting recombination in this region^12^. Similarly, in the Z chromosome the PAR is represented by the first ∼3 Mbp of sequence. Tgut191A makes up the majority of the sequence in both regions, suggesting its primary role in the synapsis of the sex chromosomes (**Figure 3F**, **Supplementary Figure 14**). Tgut191A monomers within the PAR significantly show ∼18% lower divergence (K2P 0.052 vs 0.064) and higher percent identity (94.54% vs 93.24%) than monomers outside the PAR region. This may be the result of selective pressures to improve ZW pairing efficiency or the result of repeat homogenization due to increased recombination in the PAR^50^. Copies of Tgut191A outside the PAR, particularly those in the W chromosome, may represent ancestral synaptic regions. In other chromosomes, Tgut191A is predominantly found either as subcentromeric in submetacentric macrochromosomes or as subtelomeric at the opposite end of centromeres in telocentric microchromosomes (**Figure 3A**), suggesting that it may still play a role in synapsis, since microchromosomes are spatially separated into a central compartment at interphase and during mitosis and meiosis^9^. Although not as well assembled, homologs of Tgut191A were present (>1K copies) in 15/137 VGP high-quality genomes, primarily in passerine birds, whereas in other bird lineages these may have either diverged beyond detection with current methods or a different mechanism may be at play (**Supplementary Figure 15**). The repetitive structure of sex chromosomes and the fact that the PAR region is often collapsed, even in long-read assemblies, make it difficult to obtain well-assembled sequences. Nonetheless, the sex chromosomes (either Z or W) of 17/134 VGP bird species with identified sex chromosomes were enriched of Tgut191A-like sequences in the first and last ∼3 Mbp, as well as in the intervening region. Enrichment restricted to the first ∼3 Mbp of chrZ was observed in the white-throated sparrow (*Zonotrichia albicollis*) and in the Society finch (*Lonchura striata*), whereas a pattern similar to one seen in zebra finch chrW, including both the first ∼3 Mbp and the internal region of the W chromosome, was detected in the Savannah sparrow (*Passerculus sandwichensis*).

### Terminal and interstitial telomeres

Using a newly developed tool called Telescope (Medico et al., in preparation), 1.82 Mbp of telomeric DNA were identified in the diploid genome assembly, comprising predominantly canonical CCCTAA/TTAGGG repeats (1.73 Mbp, 94.8%). Telomere variant regions (TVRs), characterized by non-canonical repeats, varied across chromosomes, with chr12 (22.1%) and paternal chr6 (0.6%) representing the extremes (**Supplementary Figure 16**). Terminal telomere length averaged 11.39±5.06 Kbp (mean±SD), with a median of 9.96 Kbp, and range 0.95-30.34 Kbp, with non-normal distribution (W=0.9236, p=1.746e-7, N=160) and no differences by haplotype or chromosome arms (**Supplementary Information**). Our mean telomere length falls within the ∼8–30 Kbp range reported by published Telomere Restriction Fragment (TRF) based studies in zebra finches^51–54^. However, TRF assays produce only per-individual means and are influenced by enzyme selection and gel resolution that limits reproducibility^55^. Unlike TRF, the T2T assembly yields nucleotide-level, per-chromosome and arm-specific resolution.

Interstitial telomeric sequences (ITSs) have been previously described in birds^56^, including recently in Passeriformes^57^. Using DNA denaturation-based methods, it was estimated that ITSs comprised 15-40% of the total telomeric DNA in zebra finch^53^. However, using cytogenetic methods, studies did not observe ITSs in the zebra finch^13^. In bTaeGut7, ITS blocks were present, and they were of variable length, with the largest arrays (>1 Kbp) within 700 Kbp of chromosomes ends of both haplotypes for chr16, 32, 33, 34 and 35 (range 1.6-8.4 Kbp) (**Supplementary Figure 16**). Clusters of multiple ITS block units were found with the same number per haplotype, three in chr35 and two in chr16 (**Figure 3G**). In contrast, chr34 showed three ITS units in the paternal haplotype and only one in the maternal haplotype. The presence of ITS in the zebra finch T2T assembly suggests that the cytogenetic studies might have missed them due to the limited resolution of fluorescent in-situ hybridization (FISH) assays. The presence of large ITS arrays in dot microchromosomes supports the hypothesis that these chromosomes have undergone structural fusion and other rearrangements.

### General chromosome organization

A typical avian genome contains ∼10 macrochromosomes and ∼30 microchromosomes^28,59,60^. Cytogenetically, 7 macrochromosome pairs and 32 microchromosome pairs, plus the Z and W chromosomes, are visible in the zebra finch karyotype (2n=80N=40, **Supplementary Figure 17, Supplementary Table 7**). Chromosome morphologies corroborate previously described karyotypes^13^ (**Supplementary Table 8**), except for chromosomes 6 and 8 that are telocentric (**Supplementary Table 9**). Consistent with previous findings^4,28,61^, zebra finch microchromosomes were significantly GC-biased (p=1.39e-10), repeat-rich (p=1.67e-2), gene-dense (p=3.96e-11), but show comparable methylation levels to macrochromosomes (p=0.19). Analyses of these patterns revealed a very specific chromosome sequence organization, particularly evident in the dot microchromosomes (5.99 Mbp ± 1.57 Mbp; **Supplementary Figure 14**). Zebra finch microchromosomes include 11 dot chromosomes (**Supplementary Table 9**), with shared sequence and epigenetic signatures. All dot chromosomes were composed of a proximal telomere, the centromere sequence (Tgut716A), one heterochromatin portion and one euchromatin portion, an (optional) satellite repeat (Tgut191A), and a distal telomere (**Figure 4a**, **Supplementary Figure 18**). This model is herein dubbed TCHE[S]T (telomere-centromere-heterochromatin-euchromatin-satellite-telomere), and is found in all dot chromosome pairs. In all dot chromosomes, the euchromatic (coding) portion also perfectly aligned with A compartments (gene rich regions), and the heterochromatic (non-coding) portion aligned with B compartments (gene poor regions), suggesting that this organization positions microchromosomal euchromatin toward the nuclear interior and heterochromatin toward the periphery^62^ (**Figure 4A**). Non-dot microchromosomes showed a more relaxed TCHEST organization, characterized by interleaved regions of heterochromatin and euchromatin, i.e. a TC(HE)_n_ST pattern. Dot chromosomes were highly methylated compared to other chromosome types (64.9% vs 45%; p=4.43e-11), and, despite their smaller size, contained a higher density of genes (63.9/Mbp vs 37.4/Mbp; p=9.91e-10) and repeats (65.9% vs. 13.2%; p=3.31e-10), and a higher GC content (53.4% vs 45.6%; p=5.24e-10). Dot chromosomes’ A compartments (euchromatin) were more GC rich than B compartments (55.1% vs. 52.4%; p<2.2e-16), repeat depleted (39.7% vs. 75.4%; p<2.2e-16), less methylated (57% vs. 63.5%; p<2.2e-16) and gene rich (77.2% vs. 38.1%; 173 genes/Mbp vs 98.5 genes/Mbp; p<2.2e-16) (**Figure 4B**). The same features in the other chromosomes classes reflect a weaker separation between the heterochromatin and euchromatin compartments (GC 43.7% vs. 40.3%; repeats 8.6% vs. 9.1%; genes 76.2% vs. 53.8%, 127 genes/Mbp vs. 72.8 genes/Mbp), showing that the enrichment patterns observed in dot chromosomes are more stereotyped.

**Figure 4.**
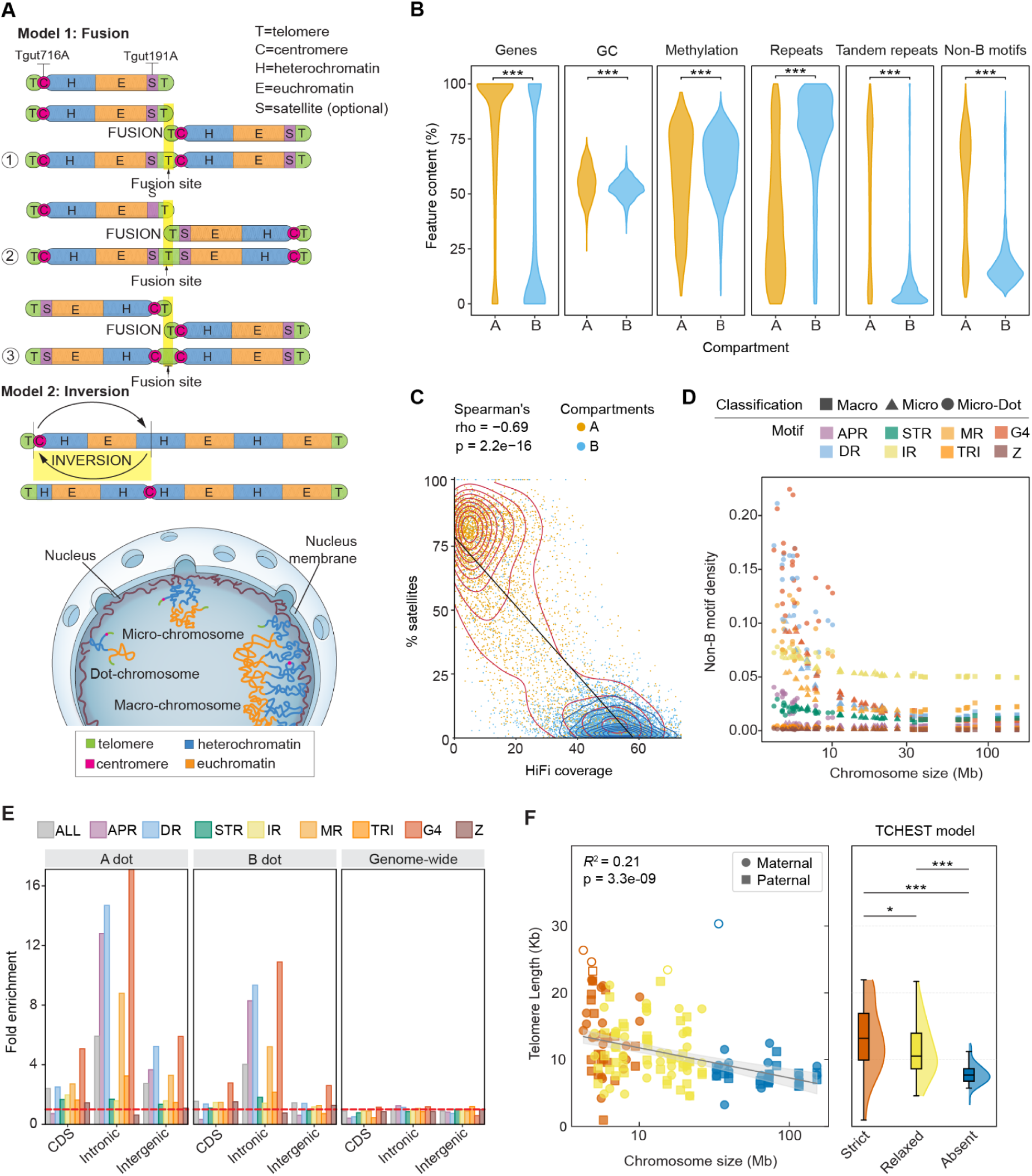
Chromosome architecture in the zebra finch. (A) In the TCHEST model, centromere satellite repeats (Tgut716A) are directly adjacent to the telomere, and are followed by heterochromatin. Heterochromatin and euchromatin can either alternate in the chromosome body (relaxed TCHEST) or exist in only two organized sections (dot chromosomes). A subtelomeric secondary satellite array (Tgut191A) can optionally be present flanking euchromatin. Macrochromosomes often show the two satellites arrays juxtaposed or interleaved, which may be suggestive of different patterns of chromosome fusion followed by rearrangements. (B) Enrichment patterns in 10kb non-overlapping windows of A and B compartments on dot chromosomes for genes, GC content, methylation, repeats, satellites and non-B DNA motifs. ‘***’ denotes p<0.001 adjusted with FDR. (C) Correlation between minisatellite repeats and HiFi coverage in dot chromosomes’ A and B compartments. Each point represents a 10 kb window coloured by its A/B assignment. Density contours are shown for each compartment to highlight data distribution. The black line represents a regression fit across all data points. (D) Non-B DNA motif density in relation to chromosome size. (E) Non-B DNA motif enrichment in and between genes in A and B compartments of dot chromosomes compared to the genome-wide average. The dashed red line on y=1 symbolises no enrichment in relation to the average motif densities in the genome. “ALL” is all non-B motifs considered together. (F) Scatter plot of chromosome size (Mbp) and telomere length (kbp). The box plots compare telomere lengths between chromosome types based on their TCHEST model categories. Hollow datapoints represent outliers.

Dot chromosomes also showed dramatic Pacbio HiFi coverage dropouts in their highly-euchromatic region (average coverage of 22.8x vs. 48.5x of heterochromatin), with a strong negative correlation with the presence of degenerate minisatellite repeats (repeat unit 10-30 bp; Spearman’s ρ=-0.69; p<2.2e-16; **Figure 4C**); whereas ONT simplex and ultra-long reads showed a much more modest dropout in this region, **Supplementary Figure 18**). The density of minisatellite repeats is indeed far higher than in heterochromatin (51.3% vs. 11.1%, **Figure 4B,C**, **Supplementary Figure 18**). Both euchromatic and heterochromatic regions across the rest of the genome showed a lower minisatellite density overall, with a far less pronounced difference in tandem repeats content (2.74% vs. 1.84%) and marginal to no HiFi coverage dropout (62.8x vs 64.7x) between compartments, also reflected in a weaker though still highly significant negative correlation between HiFi coverage and tandem repeats density (Spearman’s ρ=-0.32; p<2.2e-16, **Supplementary Figure 19**). One explanation for this dropout is that microchromosomes are enriched in non-B DNA motifs, i.e. specific sequences that have the potential to form non-canonical 3D DNA structures (**Figure 4D**), which PacBio HiFi sequencing has difficulty sequencing through^63^. Indeed, non-B DNA motifs are 56% more abundant inside previously unassembled regions than outside of them (**Supplementary Table 10**). This effect is particularly evident in the euchromatic regions of dot chromosomes (**Figure 4B**) where A-phased repeats (APR), direct repeats (DR) and G-quadruplexes (G4) colocalize with the minisatellites, causing a strong enrichment for these motifs especially in euchromatin intronic regions, not seen in genes on a genome-wide scale (**Figure 4E**). Additionally, dot chromosomes show sequence similarity in their heterochromatic sequence, which also complicated the assembly graph (**Supplementary Figure 20**). These observations are further investigated in a companion manuscript (Smeds et al., under review).

Dot microchromosomes (TCHEST strict, **Supplementary Table 9**) showed longer telomeres (mean 13.9 kbp) than non-dot microchromosomes (TCHEST relaxed, mean 11.5 kbp, p-adj=1.74e-06) and macrochromosomes (TCHEST absent, mean 8.3. kbp, p-adj=1.67e-07) (**Figure 4F**). Longer telomeres could help provide very small chromosomes a greater protection against chromosome erosion during senescence, particularly protecting the adjacent centromere. Since the TC(HE)_n_ST organization is found in all 30 zebra finch microchromosomes, it could represent an ancestral chromosome organization in birds. Alternatively, dot chromosomes may also represent a derived state in which evolutionary pressures maintain a strict TCHEST structure compatible with their small size and their spatial organization in the nucleus. Relaxation of the TCHEST model in the macrochromosomes is consistent with the hypothesis that they may have originated by fusion of microchromosomes^9^. The presence of satellite repeats Tgut716A and Tgut191A occur adjacent to or interleaved in the centromeres in all macrochromosomes including chrW and except chr6 and chrZ (**Figure 3A**), consistent with head-to-tail fusions followed by multiple local rearrangements (**Figure 4A**). Interestingly, chr6 is the only telocentric macrochromosome (37 Mbp, **Supplementary Table 9**). As in the TCHEST model, in the first 7 Mbp, Tgut716 is followed by heterochromatin and euchromatin, and then by a large Tgut191A array, compatible with a tail-to-tail fusion of a micro and a macrochromosome accompanied by the loss of the ancestral centromere. This supports previous observations that macrochromosome arms in birds often arise via Robertsonian fusions of ancestral microchromosomes, which retain some sequence features such as higher GC content and interactivity^9,64,65^. These properties erode over time post-fusion. Alternatively, the submetacentric position of macrochromosome centromeres could be the result of more recent inversion events (**Figure 4A**). This is supported by the colocalization of centromeres with evolutionary breakpoints^65^ and is consistent with the known pericentric inversion on chr5 that makes the chromosome acrocentric. This overall hypothesis is impacted by reconstructions of the bird ancestral karyotype, which generally do not support fusions of micro and macrochromosomes in the bird lineage^65^. One alternative is that the fusions may have occurred in the ancestral reptilian ancestors.

### Genome-wide sequence and structural variation

Comparison of the two haplotypes reveals both extensive sequence and structural variation as well as extensive runs of homozygosity, which is expected given the inbreeding of captive populations (**Figure 5A**, **Supplementary Tables 11,12**). Chr2 is largely homozygous in this individual. In dot chromosomes, the structural variation translates into significant differences in terms of the total size of the two haplotypes (**Figure 5B**). Variation is predominantly explained by large duplications coupled with inversions and other rearrangements that involve the heterochromatin, while euchromatic sequence is largely conserved (**Figure 5C, Supplementary Figure 21**). In macrochromosomes, only chr5 is dramatically different between the two haplotypes, and this is associated with a previously described^17,66^ large pericentric inversion, which is inherited in a Mendelian pattern^41,66^. Intrachromosomal inversions and translocations are common chromosome rearrangements in birds^67^. The inversion actually appears to involve multiple rearrangements (**Figure 5D**), which based on the alignment of the two now complete haplotypes, it can hypothesized that it likely involves either 1 inversion and 3 translocations or 2 inversions and 2 translocations (**Figure 5E**). While no evidence was found that implicates the chr5 inversion in adaptation^68^, multiple polymorphic inversions on the Z chromosome have been linked to variations in sperm morphology and reproductive traits^69^. In bTaeGut7, chrZ shows two inversions compared to bTaeGut1.4 (**Figure 5F**). Inversions on chrZ have been previously documented and classified into haplotypes A, B, C and D (where D is the intermediate between haplotypes A and C or B and C)^70^. The breakpoints of the smaller of these two inversions on chrZ have been previously shown to disrupt serine/theronine-protein kinase PAK 3-like gene, with the gene being duplicated around the breakpoint^71^. The second inversion on chrZ involves multiple genes, including RAD23B and ZNF462 that were previously associated with a sperm phenotype, namely with midpiece length^72^. Interestingly, both inversions are adjacent to two extremely large repetitive arrays, which are among the 4 gaps that still need to be fully resolved in bTaeGut7 (**Supplementary information**). A few other observed large structural variants are present between the maternal and paternal haplotypes of bTaeGut7, including duplications and inverted translocations (**Supplementary Table 13**).

**Figure 5.**
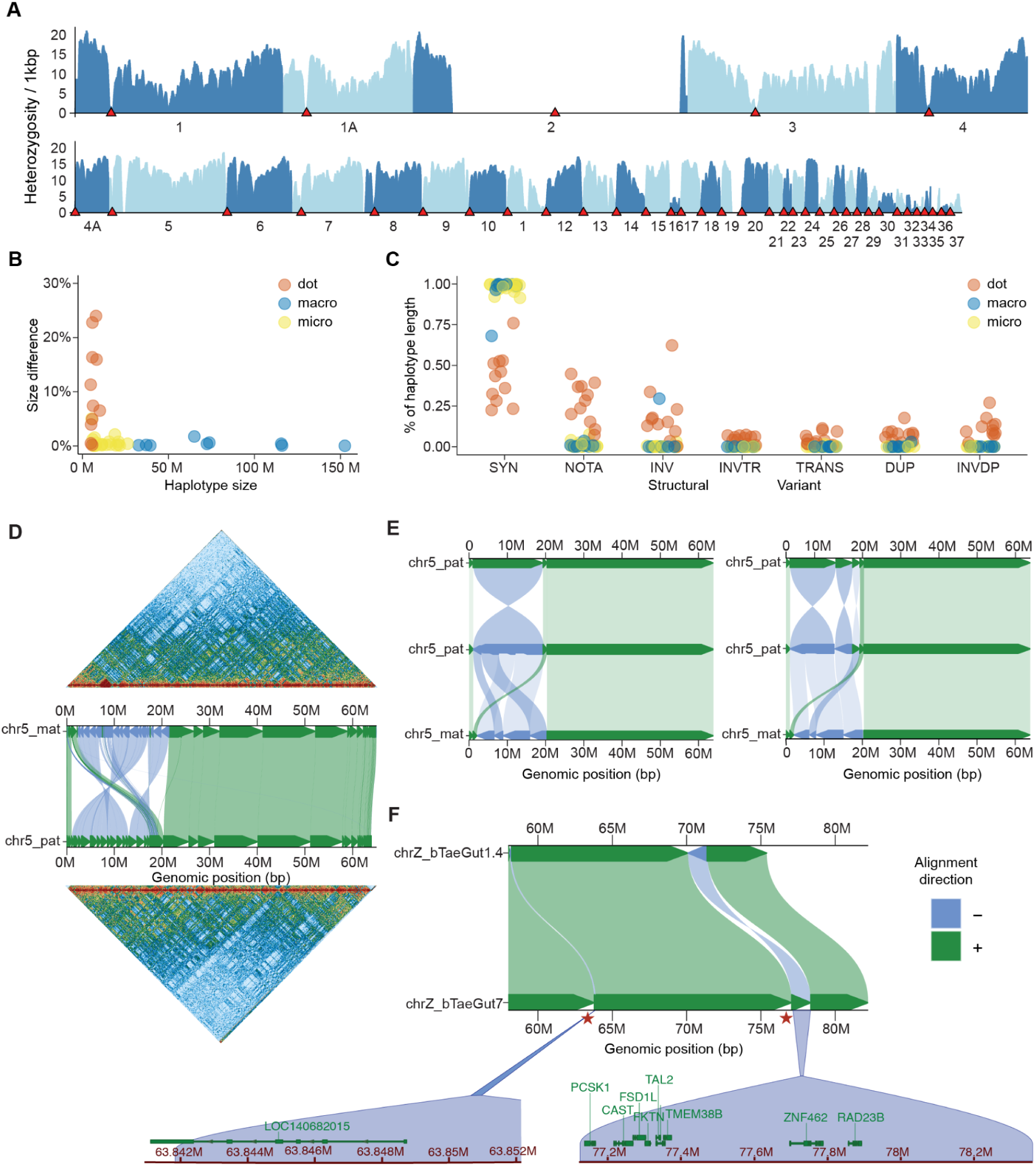
Characterization of structural variants. (A) Heterozygosity, measured as the total number of SNPs per 1kbp, across all chromosomes with centromeres labelled as red triangles. (B) Maternal and paternal haplotype size difference for each chromosome colored by the chromosome type. (C) Sequence length measured as percentage of the size of the maternal chromosome being identified as syntenic regions (SYN), not aligned between two haplotypes (NOTAL), inversions (INV), inverted translocations (INVTR), translocations (TRANS) and duplications (DUP). Each chromosome is colored by the chromosome type. (D) Chr5 for the maternal and paternal haplotypes, as displayed in PretextView and SVbyEye. (E) Schematic of translocations and inversions between the maternal and paternal haplotypes. Left side shows 1 inversion and 3 translocations and the right side, 2 inversions and 2 translocations. (F) Comparison of chrZ between bTaeGut1.4 and bTaeGut7, displayed using SVbyEye and NCBI’s Comparative Genome Browser to highlight the genes present in the inversions. Red stars indicate the location of the tangles present on chrZ.

### Characterization of genes

Gene annotation based on RNA-seq and comparative genomics identified 16,575 protein-coding genes, 184 pseudogenes, and 14,658 lncRNAs in the maternal haplotype. Of these, 308 lncRNAs, 78 protein-coding genes, and 6 pseudogenes are all located on chromosome W. In the paternal haplotype, 17,314 protein-coding genes, 177 pseudogenes, and 15,586 non-coding RNAs were identified, including annotations on chromosome Z, which harbors 987 lncRNAs, 923 protein-coding genes, and 13 pseudogenes. This includes 77 and 76 processed pseudogenes in the maternal and paternal assemblies, respectively, while 29 were detected on the sex chromosomes (**Supplementary Figure 22**; **Supplementary Information**). One-to-one orthologs of 12,785 and 13,366 human genes were identified in the maternal and paternal haplotypes, respectively, which represents 77% of zebra finch protein-coding genes.

Chromosomes displayed distinct characteristics at the gene level (**Figure 6A**). Relative to all other chromosomes, dot-chromosomes show a reduced gene length, with both smaller intron and exon sizes (**Figure 6B**). Furthermore, they showed different codon usage patterns (**Figure 6C**) with the 3rd position of most codons often changed to G or C, without changing the amino acid sequence, which is likely explained by GC-biased gene conversion in small chromosomes. Interestingly, the opposite phenomenon is observed in fruit flies that also harbor dot chromosomes^73^. As one might expect, protein coding genes and lncRNA genes scaled linearly with chromosome size (**Figure 6A**). But surprisingly, processed pseudogenes, or retrocopies (mRNA molecules retrotranscribed and inserted in the genome)^74^, did not. Dot chromosomes were slightly enriched in retrogenes, the remaining microchromosomes slightly depleted, whereas sex chromosomes show a dramatic enrichment relative to the general trend (χ²(1, N = 107) = 8.72, p = 0.0031; **Supplementary Figures 23,24**), with 16 retrocopies on Z and 13 on W (**Supplementary Tables 14-19**). On chrZ, most retrocopies (13 out 16, 81.25%) derived from autosomal parental genes (**Supplementary Figure 24, Supplementary Table 14**), consistent with a role for Z as a receptive site under dosage-compensation pressures. In contrast, all W-linked retrocopies originated from autosomal donors, reflecting the advanced degeneration and transcriptional inactivity of W. Together, these findings reveal contrasting retroposition dynamics of the two sex chromosomes, with Z functioning as both source and sink, and W acting largely as a repository with limited evidence of functional retention (**Supplementary Figure 25; Supplementary Information**).

**Figure 6.**
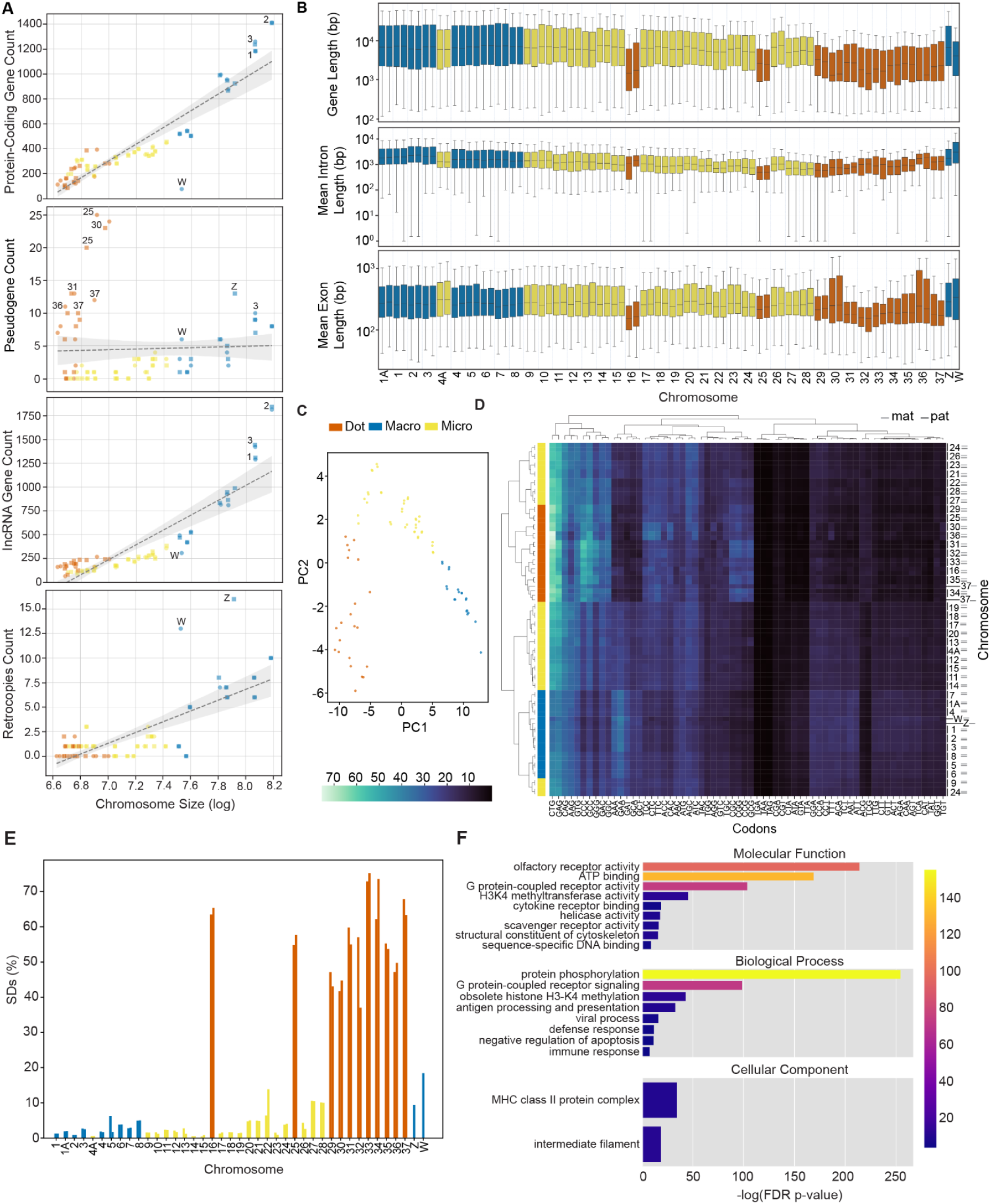
Zebra finch genome annotation. (A) Gene count versus chromosome size for protein-coding genes, pseudogenes, lncRNAs, and retrocopies. Scatter plots show the relationship between chromosome size (log-transformed) and the number of annotated genes for each category. Chromosomes are colored, considering their classification as micro/macro/dot chromosomes. Each point represents a chromosome, with labels indicating chromosome IDs. Dashed lines represent linear regression trends with 95% confidence intervals. (B) Distribution of gene lengths, mean intron lengths, and mean exon lengths across all chromosomes in the diploid assembly. Y-axis values are plotted on a log10 scale and measured in base pairs (bp). (C) The PCA plot visualizes the clustering of chromosomes based on their codon usage patterns. (D) The heatmap displays the codon usage distribution across the chromosomes, with color intensity indicating the frequency of the specific codon. Lighter shades (green/yellow) correspond to higher usage, while darker shades reflect lower usage. (E) The percentage of segmental duplications (SDs) for each chromosome, with chromosomes categorized based on their classification. (F) Barplots show the significantly enriched Gene Ontology (GO) terms in the duplicated regions of the genome for each category: Molecular Function (MF), Biological Process (BP), and Cellular Component (CC). The x-axis represents the statistical significance of enrichment (–log(FDR p-value)), while the bar color reflects the magnitude of observed-minus-expected gene counts (Obs–Exp), with warmer colors indicating greater enrichment.

Annotation identified 116 distinct miRNA families across all birds, of which 110 were present in the zebra finch assembly (**Supplementary Table 20; Supplementary Information**). Closer inspection of miRNA gene family presence/absence profiles revealed that zebra finch follows the expected passerine miRNA gene family profile, which appears to be clearly distinct from that of non-passerine birds (**Supplementary Figure 26; Supplementary Information**; **Supplementary Table 21**), with some miRNA gene families selectively associated with passerine birds (**Supplementary Figure 27; Supplementary Information**).

### Identification of segmental duplications

Segmental duplications are long regions of DNA (>1,000bp) that are repeated on the same or different chromosomes, and are the source of new gene evolution^75^. The maternal and paternal haplotypes were composed of 6.33% and 5.72% segmental duplications, respectively, with a strong variability in SDs frequencies across various chromosomes. In the maternal haplotype, 3,006 genes are located entirely within segmentally duplicated regions, including 1,806 lncRNAs, 1,113 protein-coding genes, and 87 pseudogenes. In the paternal haplotype, a comparable 3,160 genes are fully contained in segmental duplications, consisting of 1,930 lncRNAs, 1,135 protein-coding genes, and 95 pseudogenes. Dot chromosomes and sex chromosomes exhibited a prominent accumulation of segmental duplications (**Figure 6E**), particularly in the heterochromatic regions. Enrichment analyses of duplicated regions also reveal extensive lineage-specific gene expansions, notably olfactory receptor (OR) families, chromatin regulators, immune genes, kinases, and transcription factors. OR genes show the strongest diversification, with large copy numbers across several families, while other significantly enriched functions include histone methyltransferases (SETD1A), protein kinases (e.g., PAK 3-like and PIM-1 like genes). In the reverse direction, several gene duplications reported in the first zebra finch reference^17^ are not supported by bTaeGut7, indicating they are likely variable in the population or false haplotype duplication artifacts^76^. These apparent false duplications in the prior assembly include the tandem duplication of caspase-3 (a duplication occurring only in the chicken, labelled as caspase-1 but part of the caspase-3 orthogroup), the duplication of the serine protease neurotrypsin PRSS12 (single copy in both zebra finch and chicken), and the duplication of the aspartyl protease β-secretase 1 (BACE1). Together, the patterns in the T2T assembly point to a more accurately defined recent, zebra finch-specific amplification of sensory, regulatory, immune gene repertoires, potentially contributing to unique ecological adaptations and evolutionary trajectories in this lineage (**Figure 6F, Supplementary Information**).

## Discussion and future perspectives

While past zebra finch reference genomes gradually improved contiguity and accuracy, they remained constrained by persistent gaps, collapsed repeats, and unresolved microchromosomes^3^. The first T2T diploid reference genome of a songbird now overcomes these limitations, offering a complete representation of a zebra finch genome that captures its full chromosomal complexity, repeat architecture, and gene space.

An analysis of this T2T genome and other high-quality bird genomes reveals the level of conservation of satellite repeats, particularly the most abundant centromeric satellite repeat, Tgut716A, and of the second most abundant satellite, Tgut191A. The identification of Tgut716A as a conserved, methylation-depleated, kinetochore-associated satellite repeat, combined with the discovery of a candidate CENP-B box and its potential binding partner TIGD4, provides the first evidence of a complete CENP-B/CENP-B box-like system in birds. This not only overturns previous assumptions that birds may lack such a system^31,49,77^, but also introduces a new framework for investigating centromere evolution in birds and in vertebrates in general. Centromeric satellites are expected to diverge rapidly due to several factors, including^77^: molecular drive, which promotes homogenization within chromosomes but divergence between species or chromosome pairs; centromere meiotic drive, which imposes strong selection during female meiosis; conflict with kinetochore proteins, leading to positive selection; and suppressed recombination at centromeres. The presence of highly similar repeats across other passerine species raises the possibility that this sequence is ancestral to Passeriformes. Similar to Tgut716A, the Tgut191A repeat seems to have been exapted to play multiple functional roles, from the primary synaptic sequence in the sex chromosomes to potentially acting as a spacer in microchromosomes, separating the euchromatic DNA from the telomere, for instance mitigating the TPE. Tgut191A challenges the notion of satellite DNA as genomic “dark matter” and highlights how repetitive elements can be exapted to acquire precise and lineage-specific biological functions.

The availability of 40 gapless pairs of T2T zebra finch chromosomes reveals a general pattern of chromosome structural organization, which presumably is common to other species, at least in passerine birds. The resolved chromosome structure, clearly visible in microchromosomes, and especially in dot chromosomes, reveal a stereotyped recurring organizational motif (the TCHEST model). The recurrence and conservation of this organization aligns with prior observations that microchromosomes are highly conserved in sequence and structure across bird lineages. Indeed, at least 31 pairs of microchromosomes are thought to have been present in the ancestral bird–reptile karyotype ∼300 MYA, with some synteny extending to basal chordates like amphioxus^9^. Recent reconstructions even suggest that bird microchromosomes correspond closely to ancestral chordate chromosomes, possibly dating back ∼684 million years, as shown by conserved synteny with amphioxus chromosomes and proto-vertebrate karyotypes^9^. Their retention in avian genomes likely reflects functional constraints and their role as ancestral building blocks of vertebrate genomes.

The existence of the recurring TCHEST structure has broad implications for our understanding of chromosome architecture. It is a structure that aligns with nuclear architecture, chromatin accessibility, and transcriptional regulation. These patterns challenge the simplistic view of microchromosomes as merely small in functional role and instead suggest that their internal structure is gene rich, under strong constraints, and potentially co-evolved with epigenetic and 3D chromatin organizations. Microchromosomes in birds and reptiles tend to cluster centrally within the interphase nucleus, with macrochromosomes localizing peripherally^62^. This pattern is thought to be driven by genomic properties, such as high gene density and GC content, rather than chromosome size^62^. Dot microchromosomes, by maintaining strict compartmentalization of heterochromatin and euchromatin, may facilitate such nuclear architecture and interaction networks. Even when fused to macrochromosomes, ancestral microchromosomal regions tend to retain high interaction probabilities and central nuclear positions, though these features gradually degrade over time^9^.

Future T2T genome assemblies will be useful for confirming the general chromosome organization identified in the zebra finch in other birds and vertebrates. It will be interesting also to assess its conservation in zebra finch populations and in the other recognized subspecies. The zebra finch has long been considered a single species with subspecies, but recent evidence has recognized two species based on assortative mating in captivity: the Australian zebra finch (*T. castanotis*) and the Timor zebra finch (*T. guttata*)^78^. All existing reference genomes to date, including the one presented here, are based on individuals from *T. castanotis*.

Most, if not all, passerine birds possess a unique germline-restricted chromosome (GRC), which appear particularly challenging to sequence and assemble, due to the presence of paralogs recently duplicated from the rest of the genome and highly homogenized repeats^79,80^. This latter feature is especially prominent in macro-GRCs, such as the one present in the zebra finch^12^, and it will be interesting to determine what insights this GRC may reveal into zebra finch biology.

The annotation of the new assembly increased the amount of protein-coding genes in birds by 10% (over 2000 new genes). The sex chromosome relative composition was as expected, where, chrW is gene-poor and chrZ is gene-rich, consistent with W degeneration and the maintenance of extensive functional content on Z. The roughly 6% of segmental duplication in the genome are pronounced on dot and sex chromosomes. These duplicated regions harbor thousands of genes, including both coding and noncoding loci, and exhibit significant enrichment for sensory and regulatory functions. The expansion of OR families, alongside the duplication of kinases, chromatin regulators, and immune-related genes, points to a strong lineage-specific adaptive signal. Such expansions may underlie ecological specializations in communication, perception, and immunity that distinguish zebra finches and their relatives from other avian lineages. These insights refine our understanding of passerine genome evolution and underscore the utility of haplotype-resolved, gap-free assemblies in disentangling the contributions of chromosomal context, duplication, and lineage-specific innovation.

This genome also challenges current thinking about telomere dynamics. The presence of previously unreported large interstitial telomeric arrays and the unusually long telomeres on micro- and dot chromosomes suggest differences in telomere biology between chromosomes of different sizes and compositions. These differences may reflect selective pressures to protect chromosome ends in small chromosomes or could represent by-products of past chromosomal fusion and rearrangement events.

Importantly, this resource enables a new generation of functional genomics in a key model species. With gapless chromosomes, full annotation of repeat landscapes, and resolved centromeres, questions that were previously inaccessible, ranging from the regulatory logic of vocal learning circuits to the genetic basis of sex chromosome evolution, can now start to be addressed. The haplotype resolution also opens the door to diploid-aware analyses of selection, gene regulation, and chromosomal rearrangements within and between populations. The fine-grained resolution of structural variants on chromosomes such as chr5 and chrZ, including known inversions linked to sperm morphology, provides a foundation for studying the phenotypic consequences of chromosomal polymorphisms at nucleotide-level precision. This work is both the culmination of a decade of assembly improvements and a lens that redefines what is biologically visible.

## Methods

### Animal care and sample collection

Zebra finches were cared for in accordance with standards set by the American Association for Accreditation of Laboratory Animal Care (AAALAC) and Rockefeller University’s Animal Use and Care Committee (IACUC). For DNA samples from bTaeGut7 and parents, one or more 50-100µL blood samples were obtained by inserting a 30-gauge syringe (BD, Cat. #328440) in the neck jugular vein. Each sample was drawn no less than two weeks apart to obtain sufficient material without causing long-term harm to the animal.

### Genome sequencing

Data generated for bTaeGut7 included PacBio HiFi long reads, Simplex ONT long reads, standard Illumina short reads, and Arima Hi-C short reads. A total of 134.3 Gbp of PacBio HiFi coverage was generated from 4 sequencing runs of one library (7.5M reads, N50 18,348 bp). ONT data was obtained from nineteen libraries, totaling 409.1 Gbp (22.5 million reads, N50 32,843). This included data generated using standard ONT ligation sequencing, in which DNA was extracted from nucleated blood using the Monarch® HMW DNA Extraction Kit for Cells & Blood (#T3050L) with an agitation speed of 2000 rpm during cell lysis. 10uL of blood was used for HMW DNA extraction and about 2 µg of extracted DNA was used for each sequencing run. 5 Standard ONT ligation libraries were prepared using the Ligation Sequencing Kit V14 (SQK-LSK114) and sequenced on R10.4.1M flowcells, generating 128.6 Gbp of data (6.9 million reads, N50 27,257).

Ultra-high molecular weight (UHMW) DNA for ONT ultra-long sequencing was extracted from nucleated blood using the Monarch® HMW DNA Extraction Kit for Cells & Blood with an agitation speed of 300 rpm during cell lysis. Approximately 40 µg of UHMW DNA, extracted from 15-20uL nucleated blood was used for each library preparation and sequencing run. A total of 11 flowcells were used, 3 R9.4.1 and 8 R10.4.1M, with library preparation performed using the Ultra-Long DNA Sequencing Kit V1 (SQK-ULK001) and the Ultra-Long DNA Sequencing Kit V14 (SQK-ULK114), respectively. This yielded 280.5 Gbp of data (15.6 million reads, N50 38,417). Ultra-long reads were obtained by combining all ONT simplex runs and filtering for reads longer than 100 Kbp, yielding 32.1 Gbp of UL data (246,000 reads, N50 126,379). It should be noted that ONT sequencing efficiency was overall low. The sequencing pores on the flow cells became clogged rapidly, and the only way to increase yield was to perform nuclease flushes (two to three subsequent flow cell washes) and rerun the same library; a total of 15 sequencing reactions were run, as opposed to the 2-3 usually required for human samples. A significant number of pores that should have recovered did not, indicating a persistent issue. Circumstantial evidence suggests this is often the case for avian DNA using standard ONT protocols.

For PacBio HiFi sequencing, 6 µg DNA was sheared to a peak of 15 kbp to 20 kbp using a Megaruptor 3 instrument (Diagenode, Denville, NJ, USA) and then purified with Ampure PB beads (PacBio PN 100-265-900). A library was prepared with 5 µg of the DNA and SMRTbell prep kit 3.0 (Pacific Biosciences PN 102-182-700). The library was size-selected to remove fragments below 10 kbp with a Pippin HT instrument (Sage Science, Beverly, MA, USA), and then sequenced with a PacBio Sequel IIe instrument, 40-hour movie time, and PacBio Binding kit 3.2 (102-333-300). 62.6 Gbp of Arima Hi-C data was generated using the v2 kit (Arima Genomics PN A410110). Illumina parental data was generated from bTaeGut8 (maternal) and bTaeGut9 (paternal). Reads statistics were computed with rdeval^81^.

### HERRO-correction of ONT reads

Error correction of nanopore reads was performed using HERRO (Haplotype-aware ERRor cOrrection) v0.7.3+6e6c45c, which combines all-vs-all read alignment with a deep learning model that incorporates haplotype-aware information to improve single-read accuracy^82^. Due to the computational demands of the algorithm, correction was executed on a high-performance computing node (hpc_a10) on the Rockefeller University HPC cluster, equipped with an NVIDIA A10 GPU (24 GB GPU memory) and 32 CPU cores. A total of 15 unfiltered Oxford Nanopore R10 sequencing runs, comprising 323.57 Gb of data, were processed over 83 hours, 46 minutes, and 23 seconds, corresponding to an average processing time of 15.53 minutes per gigabase.

### Genome assembly

The assembly was generated using a hybrid strategy, in which multiple assemblies were generated using different sequencing data, assemblers, and tool parameters. This strategy was needed since no single assembly pipeline was able to automatically generate a gapless representation of each chromosome. All chromosomes were manually reviewed and a decision was made as to the best representative chromosome reconstruction (**Supplementary Table 22**). Nine assembly strategies were tested with Verkko v2.2.1, and two with hifiasm, including different combinations of inputs and parameters. HiFi-only chromosomes, when gapless, were favored over those from assemblies that included ONT data at the contigging stage owing to generally higher sequence accuracy of HiFi. Representative chromosomes were then combined in a single reference genome, with fully phased maternal and paternal haplotypes. The final automated assembly backbone was based primarily on two Verkko assemblies. The first Verkko assembly (asm1) used all the HiFi data to generate the assembly graph, all the ONT raw data for graph resolution, and Hi-C for haplotype phasing. The second Verkko assembly (asm5) used all HiFi data and HERRO-corrected ONT reads except R9 reads to generate the assembly, all the ONT raw data for graph resolution, and Hi-C for phasing. HERRO-corrected R9 reads were excluded to avoid potential biases introduced in the graph by undercorrected reads. This assembly required fine-tuning of the MBG parameters in Verkko at the graph building stage. Specifically, the parameter -u (Minimum average unitig abundance) was evaluated in the range 3-50 and the optimal value of 12 was chosen using the elbow method by majority rule evaluating assembly size (u=12), n50 (u=22), number of segments (u=12), average segment length (u=12), number of edges (u=12) and dead ends (u=14) in the resulting graph. With u=12, graph assembly size was 1,664,322,498, N50 was 3,036,481, number of segments was 16,269, average segment length was 102,300.23, number of edges 41,210 number of dead ends 2,193. Graph assembly size was considered reasonable given the heterozygosity of the sample.

N50 was evaluated not to be a key parameter at this stage because it does not take small segments into account. This was necessary to account for the very high coverage of HERRO-corrected ONT data and the lower quality of HERRO-corrected reads compared to HiFi reads. Ultimately, the following MBG parameters were used: k=1001,w=100,a=1,u=12,t=4,r=15000,R=4000,hpcvariantcov=25,errormasking=collapse-ms at,endkmers=no,blunt=no,keepgaps=no,guesswork=yes,copycountfilter=no,onlylocal=no,filte rwithinunitig=yes,cleaning=yes,cache=no. Asm1 contributed 24 maternal and 26 paternal T2T chromosomes. Asm5 contributed 16 maternal and 14 paternal T2T chromosomes, including all dot chromosomes and the sex chromosomes. The other assemblies used for the manual curation steps included a hifiasm assembly in Hi-C+UL mode, a Verkko assembly using HERRO-only data, and an experimental Verkko assembly using R10 uncorrected data. A hifiasm assembly in the experimental ONT-ligation-only mode was generated and was used to patch one telomeric end of maternal chr32 and one of paternal chr34. All Verkko assemblies were run on a 32-core machine with memory capped at 200G. Clock wall time was about 80h or ∼3.5d for an average assembly, except when all ONT data was used, which took over 10 days. Subsequent consensus steps were rerun on a SLURM grid to speed up runtime. Hifiasm took less than 24h on a 64-core machine.

### Graph manual curation

The assembly graphs were manually curated using BandageNG (https://github.com/asl/BandageNG) assisted by information on parental kmers, node coverage, edge coverage, and chromosome assignments from bTaeGut1.4 and bTaeGut2. Graphs were annotated for telomeres and rDNA using https://github.com/marbl/training/blob/main/part2-assemble/docker/marbl_utils/verkko_helpers/remove_nodes_add_telomere.py with a custom human rDNA sequence used as a bait. ONT UL-reads were aligned to the graphs using GraphAligner^83^. Unitigs were assigned to haplotypes using homopolymer-compressed parental hapmers (k=21) and a Gaussian Mixture Model trained using the number of parental hapmers corrected for the total number of kmers in the unitig (https://github.com/gf777/T2T-zebra-finch/blob/main/hapmers_assign_chr.py). Edge coverage was added to the graph using gfalign (https://github.com/vgl-hub/gfalign). Several iterations of manual graph resolution and consensus were performed. At each iteration, the linear consensus sequence was inspected to identify residual gaps or missing telomere ends. The HiFi-only assembly was considered the default candidate for resolution; however, due to sequencing coverage dropouts in HiFi sequencing, some chromosomes were disconnected despite the high sequencing coverage. This was particularly true for microchromosomes, and all microchromosomes were resolved in the Verkko HiFi+HERRO-corrected assembly graph. Path resolution used inspection of the graph topology coupled with parental hapmer, Hi-C information, and ONT UL-read alignments. Once resolved, the updated paths in the assembly file were used to generate a new consensus. Individual chromosomes from multiple assemblies were then added to a single combined reference. Asm1 and asm5 had 41/80 and 20/80 T2T chromosomes from the automated assembly, respectively. All other chromosomes needed manual curation to make them T2T complete. Asm5 has 2 additional T2T chrs for pat (6,19) and 2 for mat (8,19). All dot chromosomes needed manual curation. Chr2 had extensive runs of homozygosity and was collapsed in homozygous regions in the initial Verkko assembly, requiring manual resolution of the individual parental sequences. The q arm of maternal chr32 was manually resolved using the hifiasm assembly as template. The submitted assembly was screened by FCS-adaptor^84^ and 4 regions (25-95 bp) were masked for potential ONT adaptor contamination, one on chr32, three on chrZ.

### Haplotype assignment

Hapmers were computed with Meryl, both in homopolymer-compressed space and without homopolymer compression using the parental Illumina reads and the offspring Illumina reads using k=21. Since Verkko did not use parental information, unitigs were manually reassigned when the Hi-C assignment did not match the parental hapmers. Resolved chromosomes were manually assigned to the respective haplotypes using the output of meryl-lookup (https://github.com/marbl/meryl).

### Chromosome assignment

Chromosomes were initially assigned by alignment with mashmap to the prior zebra finch reference bTaeGut1.4 (GCF_003957565.2). The strategy worked well for large autosomes, but required manual intervention for microchromosomes. Due to sequence similarities between microchromosomes, and misassemblies in the microchromosomes of bTaeGut1.4, the Verkko assembly using both HiFi and HERRO-corrected ONT simplex reads was manually inspected and resolved, and then the resolved chromosomes were assigned the best matches in the bTaeGut1.4 reference according to mashmap alignments run with --pi 95/99 -s 10000/100000.

### Haplotype-specific rDNA models

A haplotype-specific rDNA model was constructed using ribotin in Verkko-mode^32^. The chicken (*Gallus gallus*) rDNA reference sequence (Genbank accession: KT445934.2) was provided along with the HiFi + HERRO-corrected ONT zebra finch assembly graph. Assembly graph annotation revealed three distinct rDNA tangles (**Supplementary Figure 6**) in the zebra finch genome that could be assigned to haplotypes based on parental *k*-mers. A fully connected tangle represented the paternal haplotype, and two tangles separated by a gap in the graph, one of which was significantly larger, represented the maternal haplotype. The nodes.txt file generated by ribotin provided a mapping of these tangles onto the assembly graph. For each tangle, ribotin produced a morphs.fa file containing the reconstructed rDNA morph sequences and their observed copy number. The expected number of copies of each morph in the genome was estimated by counting the number of kmers long as the morph size (∼18.5 Kbp) in all ONT raw reads, normalized by genome size (∼2.15 Gbp). Rdeval was used to extract read lengths of all sequence reads^81^. The resulting estimate (∼77x) represents the expected genomic coverage of a full-length rDNA repeat unit. The observed morph copy number was divided by this value to determine the expected morph count in the genome. The haplotype-specific rDNA sequence was reconstructed by randomly mixing individual morph sequences in proportion to their expected copy number using rDNA Morph Mixer, implemented as follows using a custom rDNA Morph Mixer tool. These assembled rDNA models were subsequently aligned to the edges of rDNA gaps in the assembly to determine their insertion orientation and validate their structural arrangement. Command line arguments and scripts to replicate the process are provided in Github (https://github.com/gf777/T2T-zebra-finch).

### Mitogenome assembly

To assemble the mitogenome, a first attempt was made running mitohifi v3+Galaxy0^85^. The run was not successful, presumably because too few mitochondrial reads were present compared to mitochondrial-like reads from the nuclear genome, which was ascertained by mapping with minimap2 v2.22 all HiFi and uncorrected ONT reads to an existing reference available for a different individual (bTaeGut2, CM020865.2). Alignments were then filtered using minimap2 DE tag <0.02 and length >10kbp for ONT reads, further removing ONT reads that appeared unassembled regions in the alignment. 6 nearly full-length hifi and 30 ONT reads (average read length 14,932 bp) were then used to run the assembly with mitohifi, generating a complete and error-free mitochondrial DNA sequence for bTaeGut7.

### Karyotyping

Chromosomes were harvested using two different protocols^86,87^. Embryonic fibroblast cells were cultured from a female zebra finch following previously established methods^86^. However, the standard protocols were not suitable for this species, as the chromosomes appeared shortened and the metaphases showed poor quality. To overcome these limitations, in both protocols the incubation time and colchicine concentration (2 h at 37 °C, 0.05%) were modified in order to obtain high quality mitotic metaphases. Metaphase spreads were then stained with 5% Giemsa in phosphate buffer, and karyotypes were assembled according to the International System for Standardized Avian Karyotypes^88^.

### Assembly QC

The standard VGP assembly QC pipeline was run on the diploid assembly to assess its quality using various metrics^15^. BUSCO v5.5.0+galaxy0^89^ and gfastats v1.3.6+galaxy0^90^ were run to assess assembly completeness and generate summary statistics, respectively. Meryl v1.3+galaxy7 was run on the offspring HiFi reads and parental Illumina reads, and the resulting kmer count outputs were run on Merqury v1.3+galaxy4^91^ to evaluate haplotype-specific accuracy, completeness, phase block continuity, and switch errors, and on GenomeScope 2.0.1+galaxy0^92^ to assess genome size, coverage, and heterozygosity.

The bTaeGut7 Hi-C reads were mapped onto the assembly using BWA-MEM2 v2.2.1+galaxy4^93^ and multimapped bellerophon v1.0+galaxy1^94^. Pretext maps for each haplotype were generated from the mapped reads using PretextMap v0.1.9+galaxy1 (https://github.com/sanger-tol/PretextMap) and visualized using PretextSnapshot v0.0.3+galaxy2 (https://github.com/sanger-tol/PretextSnapshot). In addition, a diploid pretext map was generated with reads mapped using minimap2 v2.28+galaxy1^95^, gap tracks calculated from gfastats, telomere tracks from seqtk v1.4+galaxy0 (https://github.com/lh3/seqtk), and bigwig files from deepTools bamCoverage v3.5.4+galaxy0 (https://github.com/gartician/deepTools-bamCoverage). QV was estimated using hybrid 31-mers from Illumina and HiFi data combined.

### DNA methylation probability (5mC)

5-methyl-cytosine (5mC) probabilities were estimated by analyzing whole genome HiFi reads with MM and ML tags. Command-line tools were installed in a Conda environment: SAMtools^96^ (for FASTA indexing and BAM handling), BEDTools^97^ (for interval operations), PacBio utilities (https://github.com/PacificBiosciences/pbbioconda) including pbtk (https://github.com/PacificBiosciences/pbtk, providing extracthifi), Jasmine (https://github.com/PacificBiosciences/jasmine, PacBio base-modification caller; package name pbjasmine), pbmm2 (https://github.com/PacificBiosciences/pbmm2, HiFi alignment frontend to minimap2), and pb-CpG-tools (https://github.com/PacificBiosciences/pb-CpG-tools, for CpG probability summarization). For each sample label, run-level BAM files were merged using ‘samtools merge’. HiFi CCS reads were extracted using extracthifi, which retains HiFi reads from full CCS output. The resulting HiFi-only BAM was used as input to base-modification calling. Per-read base modifications by running the PacBio Jasmine caller (jasmine) on the HiFi BAM to produce a modBAM containing MM and ML tags. The mod-tagged HiFi reads to the reference genome with pbmm2. Alignments were written to disk, then sorted and indexed with SAMtools. The reference-anchored CpG methylation probabilities were computed using ‘aligned_bam_to_cpg_scores’ from pb-CpG-tools, providing the sorted, indexed modBAM and the reference FASTA. The tool outputs per-site 5mC probabilities as BED and BigWig files. The methylation probabilities were accessible on the UCSC genome browser (https://genome.ucsc.edu/s/clee03/bTaeGut7_pat_ACTB).

### Terminal and interstitial telomere annotation

Telomere annotation was performed using seqtk v1.4 (https://github.com/lh3/seqtk) and a new tool developed by Jack Medico and Giulio Formenti in the Jarvis lab, called Teloscope v0.1.0 (https://github.com/vgl-hub/teloscope). Seqtk was run using -d 50000 to increase the maximum drop cutoff and rescue additional telomeres. Teloscope was run using the parameters -w 200 -s 200 -c TTAGGG -p NNNGGG -d 200 -l 500 and flags -r -g -e -m -i to scan for canonical and noncanonical repeats. Contigs were assessed and classified into eight categories based on telomere completeness (T2T, incomplete, missassembly, and none) and gap presence. Telomere completeness of previously published zebra finch reference genomes in NCBI and GenomeArk (www.genomeark.org) were evaluated to compare them to the current reference (**Supplementary Table 6**).

To identify interstitial telomeric sequences (ITS), Teloscope’s annotation and BEDtools’ merge -d 200 were used to cluster telomeric-like blocks^97^. These blocks were filtered out by their TTAGGG repeat density (>0.5). The resulting ITS coordinates were classified according to their length and chromosomal position based on the literature^98^. ITSs of above 5 Kbp and their identity with terminal telomeres were visualized with StainedGlass v0.6^58^, using a window size of 400 bp.

### Transcriptome data and genome annotation through EGAPx

The bTaeGut7 maternal and paternal haplotype assemblies were annotated with the public version of NCBI’s Eukaryotic Genome Annotation Pipeline (EGAPx) v0.4.1 (https://github.com/ncbi/egapx). EGAPx integrates alignments of RNA-seq and protein data supplemented with ab initio modeling to construct high-confidence models of protein-coding and lncRNA genes^99^. A comprehensive set of 33 Illumina RNA-seq datasets representing diverse tissues and developmental stages were used for annotation. A subset of raw read data sets were downloaded from the NCBI Sequence Read Archive (SRA) under BioProjects PRJNA1032078, PRJNA516733, and PRJNA977995 (**Supplementary Table 23**). Curated RefSeq (NP_) and RNA-seq supported model RefSeq (XP_) proteins from human, chicken, song sparrow, Barn swallow, and additional sauropsid species provided additional support for model construction. One-to-one orthologs of human protein-coding genes were computed by EGAPx based on protein homology and local synteny analysis^100^.

### Repeat annotation

Repeat annotations were generated using a combination of de novo and curated resources. EDTA2 and RepeatModeler were used for de novo identification, with outputs processed through RepeatMasker. EDTA was run with several curated libraries, including the zebra finch-specific high-frequency repeats^31^, a bTaeGut1.4-based library, RepBase (primarily derived from TaeGut1.0), and a set of CR1 elements^101^. WindowMasker was used to identify low-complexity and simple repeats. SRF (Zhang et al. 2023) and TAREAN (Novak et al. 2017) were used for de novo identification of satellite DNA sequences. The SRF library was created using the genome assembly as input, while TAREAN was performed on unassembled Nanopore reads from zebra finch testis (Ruiz-Ruano et al. 2025) after multiple rounds of FASTQ subsetting using seqtk (v1.2). Sequences 20 bp or shorter were filtered out and the remaining sequences were manually curated. Additional satellites were identified manually by analysis of gap-adjacent repeats in previous zebra finch assembly versions. RepeatMasker v4.1.5 was used to localize the identified satellites in the assembly. Additional avian libraries were reviewed and incorporated as appropriate.

To further refine the non-satellite repeat annotation, the output repeat library file generated by EDTA was curated using TEtrimmer (v1.5.4), and TE consensus sequences were subsequently verified through manual inspection. TE classifications inconsistent with TEtrimmer curation results, including coverage, self-alignment, and conserved domain identification, were manually corrected, and the initial repeat library was updated with revised TE identifiers where TEtrimmer provided improved classifications. The curated repeat library was provided to EDTA using the “--curatedlib” option to reannotate the genome, generating a more accurate repeat annotation.

### Non-B DNA motif annotation

A-phased repeats (APR), direct repeats (DR), inverted repeats (IR), mirror repeats (MR) short tandem repeats (STR) and Z-DNA motifs were annotated with gfa^102^. Triplex motifs (TRI) were marked by gfa and extracted from mirror repeats (‘subset=1’). G-quadruplexes (G4) were annotated using Quadron^103^. Predicted G4s with score “NA” were removed (typically 1-2 motifs per chromosome - Quadron does not predict scores for sequences closer than 50bp from the ends of the input sequences).

### Centromere identification

To map known centromeric and distal markers, 64 previously described primer pairs were used^41^. All primers were queried against our T2T maternal (mat+Z) and paternal bTaeGut7 haplotype assemblies using blastn -task blastn-short. Outputs from both haplotypes were concatenated and filtered to retain only chromosome-specific primer hits with forward/reverse pairs <1 kbp apart. Because of divergence from the original reference^17^ used in primer design, primers lacking paired hits were rescued by selecting the single highest bitscore match per chromosome. Final primer coordinates were exported as BED and manually inspected to confirm their positions in our T2T assembly. The similarity of Tgut716A to crowSat1 was detected using Censor (https://www.girinst.org/censor/). Scripts used in centromere analyses are available here https://github.com/gf777/T2T-zebra-finch.

### TIGD4 expression analysis

To confirm overexpression of TIGD4, a transcriptome analysis on RNA-seq data from NCBI BioProject PRJNA768106 was performed by aligning reads to the paternal haploid zebra finch genome with STAR and quantifying gene counts using featureCounts. Differential expression was then assessed with DESeq2 both between brain and testis and among three temperature conditions (27 °C, 35 °C and 43 °C) for each tissue. No significant differences in TIGD4 expression were observed across the 27 °C, 35 °C and 43 °C conditions in either brain or testis samples.

### Annotation of structural variants

To initially identify large structural variants, maternal and paternal haplotypes were aligned to each other using MashMap (https://github.com/marbl/MashMap) with the number of threads set to 8, and the resultant output file displayed in JBrowse2 as a dotplot (https://jbrowse.org/jb2/). All structural variants seen in the alignment data were recorded and confirmed in the HiC data displayed in PretextView (as a PretextMap, https://github.com/sanger-tol/PretextView). The same method was utilized to align the previous reference, bTaeGut1.4, to the maternal and paternal haplotypes individually. Genome-wide sequence and structural variation was quantified using SyRI v1.6.3 with default filtering^104^. Synteny and structural rearrangements were visualized using plotsr v1.1.1^105^. Genome-wide heterozygosity was calculated using 1Mbp window size and 100kbp step size and plotted using *ggplot2*in R ( https://github.com/YingChen94/zebra_finch_T2T_Fig5).

### Annotation of AB compartments

Adaptor sequences from Arima Hi-C paired-end reads were trimmed using cutadapt v5.1^106^. ∼414 million trimmed reads were aligned against the diploid, maternal+ Z, and the paternal+W assemblies using BWA-MEM v0.7.17-r1188^107^ with the -SP5M flags. Quality control of aligned BAMs was assessed with SAMtools stats^96^ and rdeval^81^. To annotate the three-dimensional chromatin organization, the Open Chromosome Collective (Open2C)’s pipeline comprising pairtools^108^, cooler^109^, and cooltools was used^110^. To extract Hi-C read pairs from name-sorted BAMs, pairtools parse with the default --walks-policy mask was used. After pairtools sort and pairtools dedup with --max-mismatch 0, 41.9, 88, and 87.8 million, unique pairs (UU) remained, which represents 81.8%, 67.5% and 67.4% of total valid pairs for the diploid, maternal+Z and paternal+W assemblies, respectively. Using cooler cload pairs, .cool files with 10 kbp bins were generated and then normalized with cooler balance. To explore the relationship of contacts vs. distance, cooltools expected-cis on the .cool file to calculate the contact probability or P(s) curve was used. To obtain a multi-resolution .mcool, cooler zoomify at 10, 20, 50, 100, and 200 kbp resolutions was used. Bedtools coverage and the EGAPx gene annotation were used to generate a gene density track for all resolutions. Cooltools eigs-cis on the .mcool file was used to obtain eigenvectors using a gene coverage bedgraph as the phasing track, to flip their signals. Positive correlations with the first eigenvector (E1) and the phasing track were used to determine A/B compartments. Saddle plots were generated to represent the enrichment profiles or preferences captured by the eigenvectors with cooltools saddle. Finally, topologically associated domain (TAD) boundaries were computed at 200kbp using cooltools insulation.

### Identification of unassembled regions

Previously unassembled regions were identified by aligning the new diploid assembly to the previous reference genome GCF_003957565.2 (bTaeGut 1.4) using FASTGA^111^. The initial unassembled regions BED file was filtered to retain only regions longer than 1Kb. Subsequently, each gene was assessed for its spatial relationship to the unassembled regions—classified as entirely included, partially overlapping, or located outside of any unassembled regions segment. For overlapping cases, the proportion of the gene’s length covered by unassembled regions was calculated. Genes completely included inside the unassembled regions boundaries, or genes exhibiting an overlap equal to or greater than 75% of their total length were classified as putative novel/improved genes. To verify the novelty, the gene sequences were blasted against the bTaeGut1.4 to identify the presence or not of hits. Hits matching on other chromosomes were excluded. To characterize the genomic content of unassembled regions, unassembled regions coordinates were intersected with gene annotations to quantify the amount of overlap per region and feature type. The sum of overlapping base pairs for each gene biotype (e.g., protein-coding, lncRNA, pseudogene) was calculated. When an unassembled region overlapped multiple biotypes, the region was labeled as “overlapping_features”. To identify genomic features enriched in unassembled regions, a feature-wise Fisher’s exact test stratified by chromosome was performed. For each feature type, a contingency table was built comparing the number of base pairs of that feature within unassembled regions of a specific chromosome versus all other feature types in that chromosome, and the same feature in all the other chromosomes versus all other features. The resulting 2×2 table was used to compute an odds ratio and a p-value via a two-sided Fisher’s exact test. This analysis was repeated for each feature across all chromosomes. Multiple testing correction was applied using the Benjamini-Hochberg false discovery rate (FDR) method. Adjusted p-values (FDR ≤ 0.05) and log2-transformed odds ratios were used to determine and visualize significant enrichment.

### Gene family evolution

Gene family evolution in the zebra finch genome was assessed using orthology-based comparative analysis across multiple bird species. Gene families were clustered using OrthoFinder.

### microRNA evaluation

The precursor microRNA (miRNA) repertoire of the Zebra finch genome and 131 other bird genomes available through the VGP were predicted using MirMachine^112^. There were 32 bird genomes from within Passeriformes, 99 from other groups (**Supplementary Table 20**). Bitscore-filtered predictions were converted to a miRNA gene family presence/absence matrix across all 132 species. The presence/absence matrix was subsequently used to calculate Jaccard similarity across all 132 bird species using the metaMDS function of the vegan package in R^113^. The value of k used for Non-Metric Multidimensional Scaling (NMDS) was selected by performing clustering with k values from 1-10 (maximum 1000 iterations per run), and extracting minimum stress values for each run. A k value of 5 was used (this was the smallest k value with a minimum stress value <0.1. A set of 15 miRNAs previously associated with vocal learning and song were assessed for levels of conservation across the dataset (**Supplementary Table 21**). Where required, the designations of these gene families were converted from miRBase to mirGeneDB nomenclature^114^. Using the map_dollo_changes function of the Claddis package^115–117^ miRNA family presence/absence was used to reconstruct gene gain and loss patterns across bird species. For each miRNA family, reconstruction by Dollo parsimony was performed 100 times, retaining the most commonly-supported point of gene gain and point(s) of gene loss.

### Retrocopy Identification and Expression Analysis

Retrocopies were identified using an improved version of the RCPedia pipeline^118^. mRNA sequences from annotated protein-coding genes were extracted with gffread^119^ and aligned to the genome using LAST^120^ (lastal -D1000). Candidate retrocopies were retained if alignments exceeded 120 bp and were located at least 200 kb from the parental locus to minimize inclusion of segmental duplications. To reflect the reverse transcription mechanism, candidates were required to preserve at least one of the last three exon–exon junctions of the parental transcript. Alignments containing ≥40% repetitive sequence (RepeatMasker, simpleRepeats, windowMasker) were excluded. Retrocopies derived from the same parental gene were required to be ≥500 kb apart; those overlapping ≥3 annotated exons or arising from gene families with >5 redundant copies were discarded. For each retrocopy, the best source mRNA was selected based on sequence identity and match proportion; ties were resolved by the highest score ratio. Continuous alignments separated by ≤6 kb were merged to account for gaps introduced by repetitive elements. To ensure accurate expression quantification, we applied a simulation-based filtering strategy. Paired-end RNA-seq data (30 million reads, HiSeq 76 profile) were simulated with SANDY v1.0^121^, excluding exonic retrocopy regions. Quantification was performed with kallisto v0.48.0 (-b 100)^122^, and retrocopies with detectable expression in simulations were excluded as potentially confounding. RNA-seq quantification was then carried out on seven datasets (SRR8551559, SRR8551561, SRR8551563, SRR8551565, SRR8551567, SRR8695298, SRR8695300) using the same kallisto parameters. To confirm expression, kallisto estimates were complemented with alignments generated by STAR v2.7.7a ^123^ (--outSAMtype BAM SortedByCoordinate --outSAMattributes NH HI AS nM MD XS). Only retrocopies supported by at least one uniquely mapped read were retained.

### Synteny of TIGD4

Chromosome level genome assemblies and gene annotations for six species (*Taeniopygia guttata* GCF_048771995.1, *Calypte anna* GCF_003957555.1, *Dromaius novaehollandiae* GCF_036370855.1, *Homo sapiens* GCF_009914755.1, *Eublepharis macularius* GCF_028583425.1, *Pelodiscus sinensis* GCF_049634645.1) were obtained from NCBI. Protein sequences were inferred from the GFF3 annotations using gffread. Gene annotations were then parsed into BED format, ensuring proper formatting and consistent headers between the BED and protein FASTA files. Synteny between the species was inferred using GENESPACE (ref https://elifesciences.org/articles/78526). The resulting orthology information was used to generate whole-chromosome synteny plots. Local synteny around the TIGD4 gene was visualised using scripts adapted from the TidyLocalSynteny repository (https://github.com/cxli233/TidyLocalSynteny).

### Phylogenetic distribution of Tgut191A-like repeats across VGP bird genomes

The Phylogenetic tree among VGP bird genomes was inferred by the OpenTree API (https://github.com/ropensci/rotl). The abundance of Tgut191A-like repeats in bird genomes was assessed using a database of satellite DNA sequences from all available VGP bird genomes, generated through Satellome (https://github.com/aglabx/satellome). The canonical Tgut191A sequence was mapped to this database via BLAST^124^ to identify repetitive sequences homologous to Tgut191A. The Tandem Repeats Finder (TRF) tool^125^ was then employed to extract monomer units from sequence hits that remained in repetitive arrays. Monomers were then aligned with MAFFT^126^ to standardize orientation within each species, and seventeen species-specific consensus sequences were generated with HMMER^127^ from these alignments. These consensus sequences were subsequently supplied to RepeatMasker (https://www.repeatmasker.org) to annotate a final, consistent set of Tgut191A-like elements across all bird species (https://github.com/duartetorreserick/tgut191a_analysis_VGP_birds).

## Data Availability

The reference genome is currently available and annotated in NCBI under accessions GCA_048771995.1 (maternal+Z) and GCA_048772025.1 (paternal autosomes). EGAP gene annotation is available under accession GCF_048771995.1-RS_2025_04. Raw sequencing data for bTaeGut7 (offspring), bTaeGut8 (maternal), and bTaeGut8 (paternal) are available in the NCBI SRA under BioProject accessions PRJNA1241433, PRJNA1241434, and PRJNA1241435, respectively.

## Author contributions

G.F. performed the genome assembly, satellite-repeat analyses, and wrote the original draft. S.B. performed cell culture, chromosome extraction and karyotyping. H.C., H.L., S.K., and D.A. contributed to genome assembly. A.R. performed genome polishing analyses. M.S. performed the gene, repeat and transposable elements annotations, the orthology analysis and evaluated the gene families enriched in segmental duplications. H.B.C., L.D.O., P.A.G., and R.M. analyzed processed pseudogenes and retrocopies. J.F., N.F., and M.J.O’C. performed microRNA annotation and analyses. M.B. and G.D. collected blood samples for data generation. N.J. generated data (DNA isolation; simplex and ultra-long ONT sequencing). B.K. performed data pre-processing, organization, and quality control. C.L. computed DNA-methylation probabilities from HiFi reads and built the UCSC Genome Browser assembly hub. K.M. and L.S. annotated non-B DNA. P.M. and T.M. contributed genome annotation. K.M. performed manual curation and structural-variant analyses. J.M. annotated and analyzed telomeres, centromeres, and three-dimensional genome features. Y.N., K.G., and S.O. curated transposable elements. S.S. contributed to initial manual curation, performed enrichment analyses for dot chromosomes and generated some of the figures. T.T. generated Hi-C data. Y.C. analyzed nucleotide diversity and structural variation. A.K. and M.P. analyzed satellite DNA and identified candidate proteins potentially linking centromeric satellite DNA to binding activity. E.D. performed an analysis of the presence of Tgut191A-like sequence across VGP bird genomes. G.F. and E.D.J. conceived the study and coordinated the project, and J.B. coordinated the sequencing. All authors read and approved the final manuscript.

## Acknowledgments

We thank Jennifer Graves and Hardip Patel for the useful discussion on chromosome rearrangements. E.D.J. was supported by HHMI. J.F., N.F., and M.J.O’C would like to thank the Biotechnology and Biological Sciences Research Council UK (award number BB/X007332/1) for funding and University of Nottingham HPC facilities. M.J.O’C would like to thank the Leverhulme Trust for her personal fellowship (RF-2024-492). The work of P.M. and T.M. was supported by the National Center for Biotechnology Information of the National Library of Medicine (NLM), National Institutes of Health (NIH). This work was supported, in part, by the Intramural Research Program of the US National Human Genome Research Institute, National Institutes of Health (A.M.P., A.R., D.A., J.Kim, S.K.). The contributions of the NIH authors are considered Works of the United States Government. The findings and conclusions presented in this paper are those of the authors and do not necessarily reflect the views of the NIH or the U.S. Department of Health and Human Services. This work utilized the computational resources of the NIH HPC Biowulf cluster (http://hpc.nih.gov).

## Declaration of generative AI and AI-assisted technologies in the manuscript preparation process

During the preparation of this work some of the authors used ChatGPT in order to review or improve text and code. After using this tool/service, the authors reviewed and edited the content as needed and take full responsibility for the content of the published article.

## References

1. Kapusta, A., and Suh, A. (2017). Evolution of bird genomes-a transposon’s-eye view. Ann. N. Y. Acad. Sci. 1389, 164–185.

2. Jarvis, E.D., Güntürkün, O., Bruce, L., Csillag, A., Karten, H., Kuenzel, W., Medina, L., Paxinos, G., Perkel, D.J., Shimizu, T., et al. (2005). Avian brains and a new understanding of vertebrate brain evolution. Nat. Rev. Neurosci. 6, 151–159.

3. Rhie, A., McCarthy, S.A., Fedrigo, O., Damas, J., Formenti, G., Koren, S., Uliano-Silva, M., Chow, W., Fungtammasan, A., Kim, J., et al. (2021). Towards complete and error-free genome assemblies of all vertebrate species. Nature 592, 737–746.

4. Kim, J., Lee, C., Ko, B.J., Yoo, D.A., Won, S., Phillippy, A.M., Fedrigo, O., Zhang, G., Howe, K., Wood, J., et al. (2022). False gene and chromosome losses in genome assemblies caused by GC content variation and repeats. Genome Biol. 23, 204.

5. Srikulnath, K., Ahmad, S.F., Singchat, W., and Panthum, T. (2021). Why do some vertebrates have microchromosomes? Cells 10, 2182.

6. Mathers, T.C., Paulini, M., Sotero-Caio, C.G., and Wood, J.M.D. (2025). MicroFinder: Conserved gene-set mapping and assembly ordering for manual curation of bird microchromosomes. Genomics.

7. Liu, J., Wang, Z., Li, J., Xu, L., Liu, J., Feng, S., Guo, C., Chen, S., Ren, Z., Rao, J., et al. (2021). A new emu genome illuminates the evolution of genome configuration and nuclear architecture of avian chromosomes. Genome Res. 31, 497–511.

8. van Brink, J.M. (1959). L’expression morphologique de la digamétie chez les sauropsidés et les monotrèmes. Chromosoma 10, 1–72.

9. Waters, P.D., Patel, H.R., Ruiz-Herrera, A., Álvarez-González, L., Lister, N.C., Simakov, O., Ezaz, T., Kaur, P., Frere, C., Grützner, F., et al. (2021). Microchromosomes are building blocks of bird, reptile and mammal chromosomes. bioRxiv. 10.1101/2021.07.06.451394.

10. Mello, C.V. (2014). The zebra finch, Taeniopygia guttata: an avian model for investigating the neurobiological basis of vocal learning. Cold Spring Harb. Protoc. 2014, 1237–1242.

11. O’Connor, R.E., Romanov, M.N., Kiazim, L.G., Barrett, P.M., Farré, M., Damas, J., Ferguson-Smith, M., Valenzuela, N., Larkin, D.M., and Griffin, D.K. (2018). Reconstruction of the diapsid ancestral genome permits chromosome evolution tracing in avian and non-avian dinosaurs. Nat. Commun. 9, 1883.

12. Pigozzi, M.I., and Solari, A.J. (1998). Germ cell restriction and regular transmission of an accessory chromosome that mimics a sex body in the zebra finch, Taeniopygia guttata. Chromosome Res. 6, 105–113.

13. Dos Santos, M. da S., Kretschmer, R., Frankl-Vilches, C., Bakker, A., Gahr, M., O Brien, P.C.M., Ferguson-Smith, M.A., and de Oliveira, E.H.C. (2017). Comparative cytogenetics between two important songbird, models: The zebra finch and the canary. PLoS One 12, e0170997.

14. Giani, A.M., Gallo, G.R., Gianfranceschi, L., and Formenti, G. (2020). Long walk to genomics: History and current approaches to genome sequencing and assembly. Comput. Struct. Biotechnol. J. 18, 9–19.

15. Larivière, D., Abueg, L., Brajuka, N., Gallardo-Alba, C., Grüning, B., Ko, B.J., Ostrovsky, A., Palmada-Flores, M., Pickett, B.D., Rabbani, K., et al. (2024). Scalable, accessible and reproducible reference genome assembly and evaluation in Galaxy. Nat. Biotechnol. 42, 367–370.

16. Stapley, J., Birkhead, T.R., Burke, T., and Slate, J. (2008). A linkage map of the zebra finch Taeniopygia guttata provides new insights into avian genome evolution. Genetics 179, 651–667.

17. Warren, W.C., Clayton, D.F., Ellegren, H., Arnold, A.P., Hillier, L.W., Künstner, A., Searle, S., White, S., Vilella, A.J., Fairley, S., et al. (2010). The genome of a songbird. Nature 464, 757–762.

18. Koren, S., Rhie, A., Walenz, B.P., Dilthey, A.T., Bickhart, D.M., Kingan, S.B., Hiendleder, S., Williams, J.L., Smith, T.P.L., and Phillippy, A.M. (2018). De novo assembly of haplotype-resolved genomes with trio binning. Nat. Biotechnol. 10.1038/nbt.4277.

19. Nurk, S., Koren, S., Rhie, A., Rautiainen, M., Bzikadze, A.V., Mikheenko, A., Vollger, M.R., Altemose, N., Uralsky, L., Gershman, A., et al. (2022). The complete sequence of a human genome. Science 376, 44–53.

20. Rautiainen, M., Nurk, S., Walenz, B.P., Logsdon, G.A., Porubsky, D., Rhie, A., Eichler, E.E., Phillippy, A.M., and Koren, S. (2023). Telomere-to-telomere assembly of diploid chromosomes with Verkko. Nat. Biotechnol. 41, 1474–1482.

21. Cheng, H., Asri, M., Lucas, J., Koren, S., and Li, H. (2024). Scalable telomere-to-telomere assembly for diploid and polyploid genomes with double graph. Nat. Methods 21, 967–970.

22. Antipov, D., Rautiainen, M., Nurk, S., Walenz, B.P., Solar, S.J., Phillippy, A.M., and Koren, S. (2025). Verkko2 integrates proximity-ligation data with long-read De Bruijn graphs for efficient telomere-to-telomere genome assembly, phasing, and scaffolding. Genome Res. 35, 1583–1594.

23. He, Y., Chu, Y., Guo, S., Hu, J., Li, R., Zheng, Y., Ma, X., Du, Z., Zhao, L., Yu, W., et al. (2023). T2T-YAO: A telomere-to-telomere assembled diploid reference genome for Han Chinese. Genomics Proteomics Bioinformatics 21, 1085–1100.

24. Wu, H., Luo, L.-Y., Zhang, Y.-H., Zhang, C.-Y., Huang, J.-H., Mo, D.-X., Zhao, L.-M., Wang, Z.-X., Wang, Y.-C., He-Hua, E., et al. (2024). Telomere-to-telomere genome assembly of a male goat reveals variants associated with cashmere traits. Nat. Commun. 15, 10041.

25. Luo, L.-Y., Wu, H., Zhao, L.-M., Zhang, Y.-H., Huang, J.-H., Liu, Q.-Y., Wang, H.-T., Mo, D.-X., EEr, H.-H., Zhang, L.-Q., et al. (2025). Telomere-to-telomere sheep genome assembly identifies variants associated with wool fineness. Nat. Genet. 57, 218–230.

26. Yoo, D., Rhie, A., Hebbar, P., Antonacci, F., Logsdon, G.A., Solar, S.J., Antipov, D., Pickett, B.D., Safonova, Y., Montinaro, F., et al. (2025). Complete sequencing of ape genomes. Nature 641, 401–418.

27. Liu, J., Li, Q., Hu, Y., Yu, Y., Zheng, K., Li, D., Qin, L., and Yu, X. (2024). The complete telomere-to-telomere sequence of a mouse genome. Science 386, 1141–1146.

28. Huang, Z., Xu, Z., Bai, H., Huang, Y., Kang, N., Ding, X., Liu, J., Luo, H., Yang, C., Chen, W., et al. (2023). Evolutionary analysis of a complete chicken genome. Proc. Natl. Acad. Sci. U. S. A. 120, e2216641120.

29. Jarvis, E.D., Mirarab, S., Aberer, A.J., Li, B., Houde, P., Li, C., Ho, S.Y.W., Faircloth, B.C., Nabholz, B., Howard, J.T., et al. (2014). Whole-genome analyses resolve early branches in the tree of life of modern birds. Science 346, 1320–1331.

30. Cheng, H., Jarvis, E.D., Fedrigo, O., Koepfli, K.-P., Urban, L., Gemmell, N.J., and Li, H. (2022). Haplotype-resolved assembly of diploid genomes without parental data. Nat. Biotechnol. 10.1038/s41587-022-01261-x.

31. Takki, O., Komissarov, A., Kulak, M., and Galkina, S. (2022). Identification of centromere-specific repeats in the zebra finch genome. Cytogenet. Genome Res. 162, 55–63.

32. Rautiainen, M. (2024). Ribotin: automated assembly and phasing of rDNA morphs. Bioinformatics 40, btae124.

33. Stults, D.M., Killen, M.W., Pierce, H.H., and Pierce, A.J. (2008). Genomic architecture and inheritance of human ribosomal RNA gene clusters. Genome Res. 18, 13–18.

34. Piégu, B., Arensburger, P., Guillou, F., and Bigot, Y. (2018). But where did the centromeres go in the chicken genome models? Chromosome Res 26, 297–306.

35. Shang, W.-H., Hori, T., Toyoda, A., Kato, J., Popendorf, K., Sakakibara, Y., Fujiyama, A., and Fukagawa, T. (2010). Chickens possess centromeres with both extended tandem repeats and short non-tandem-repetitive sequences. Genome Res. 20, 1219–1228.

36. Matzke, M.A., Varga, F., Berger, H., Schernthaner, J., Schweizer, D., Mayr, B., and Matzke, A.J. (1990). A 41-42 bp tandemly repeated sequence isolated from nuclear envelopes of chicken erythrocytes is located predominantly on microchromosomes. Chromosoma 99, 131–137.

37. Deryusheva, S., Krasikova, A., Kulikova, T., and Gaginskaya, E. (2007). Tandem 41-bp repeats in chicken and Japanese quail genomes: FISH mapping and transcription analysis on lampbrush chromosomes. Chromosoma 116, 519–530.

38. Saifitdinova, A.F., Derjusheva, S.E., Malykh, A.G., Zhurov, V.G., Andreeva, T.F., and Gaginskaya, E.R. (2001). Centromeric tandem repeat from the chaffinch genome: Isolation and molecular characterization. Genome 44, 96–103.

39. Mc Cartney, A.M., Shafin, K., Alonge, M., Bzikadze, A.V., Formenti, G., Fungtammasan, A., Howe, K., Jain, C., Koren, S., Logsdon, G.A., et al. (2022). Chasing perfection: validation and polishing strategies for telomere-to-telomere genome assemblies. Nat. Methods 19, 687–695.

40. Jarvis, E.D., Formenti, G., Rhie, A., Guarracino, A., Yang, C., Wood, J., Tracey, A., Thibaud-Nissen, F., Vollger, M.R., Porubsky, D., et al. (2022). Semi-automated assembly of high-quality diploid human reference genomes. Nature 611, 519–531.

41. Knief, U., and Forstmeier, W. (2016). Mapping centromeres of microchromosomes in the zebra finch (Taeniopygia guttata) using half-tetrad analysis. Chromosoma 125, 757–768.

42. Gershman, A., Sauria, M.E.G., Guitart, X., Vollger, M.R., Hook, P.W., Hoyt, S.J., Jain, M., Shumate, A., Razaghi, R., Koren, S., et al. (2022). Epigenetic patterns in a complete human genome. Science 376, eabj5089.

43. Gartenberg, M. (2009). Heterochromatin and the cohesion of sister chromatids. Chromosome Res. 17, 229–238.

44. Sullivan, K.F., and Glass, C.A. (1991). CENP-B is a highly conserved mammalian centromere protein with homology to the helix-loop-helix family of proteins. Chromosoma 100, 360–370.

45. Gamba, R., and Fachinetti, D. (2020). From evolution to function: Two sides of the same CENP-B coin? Exp. Cell Res. 390, 111959.

46. Sasao, T., Itoh, N., Takano, H., Watanabe, S., Wei, G., Tsukamoto, T., Kuzumaki, N., and Takimoto, M. (2004). The protein encoded by cancer/testis gene D40/AF15q14 is localized in spermatocytes, acrosomes of spermatids and ejaculated spermatozoa. J Reprod Fertil 128, 709–716.

47. Bao, W., Kojima, K.K., and Kohany, O. (2015). Repbase Update, a database of repetitive elements in eukaryotic genomes. Mob. DNA 6, 11.

48. Weissensteiner, M.H., Pang, A.W.C., Bunikis, I., Höijer, I., Vinnere-Petterson, O., Suh, A., and Wolf, J.B.W. (2017). Combination of short-read, long-read, and optical mapping assemblies reveals large-scale tandem repeat arrays with population genetic implications. Genome Res. 27, 697–708.

49. Melters, D.P., Bradnam, K.R., Young, H.A., Telis, N., May, M.R., Ruby, J.G., Sebra, R., Peluso, P., Eid, J., Rank, D., et al. (2013). Comparative analysis of tandem repeats from hundreds of species reveals unique insights into centromere evolution. Genome Biol. 14, R10.

50. Otto, S.P., Pannell, J.R., Peichel, C.L., Ashman, T.-L., Charlesworth, D., Chippindale, A.K., Delph, L.F., Guerrero, R.F., Scarpino, S.V., and McAllister, B.F. (2011). About PAR: the distinct evolutionary dynamics of the pseudoautosomal region. Trends Genet. 27, 358–367.

51. Haussmann, M.F., and Vleck, C.M. (2002). Telomere length provides a new technique for aging animals. Oecologia 130, 325–328.

52. Criscuolo, F., Bize, P., Nasir, L., Metcalfe, N.B., Foote, C.G., Griffiths, K., Gault, E.A., and Monaghan, P. (2009). Real-time quantitative PCR assay for measurement of avian telomeres. J. Avian Biol. 40, 342–347.

53. Foote, C.G., Vleck, D., and Vleck, C.M. (2013). Extent and variability of interstitial telomeric sequences and their effects on estimates of telomere length. Mol. Ecol. Resour. 13, 417–428.

54. Salmón, P., Millet, C., Selman, C., and Monaghan, P. (2021). Growth acceleration results in faster telomere shortening later in life. Proc. Biol. Sci. 288, 20211118.

55. Lai, T.-P., Wright, W.E., and Shay, J.W. (2018). Comparison of telomere length measurement methods. Philos. Trans. R. Soc. Lond. B Biol. Sci. 373. 10.1098/rstb.2016.0451.

56. Nanda, I., Schrama, D., Feichtinger, W., Haaf, T., Schartl, M., and Schmid, M. (2002). Distribution of telomeric (TTAGGG)(n) sequences in avian chromosomes. Chromosoma 111, 215–227.

57. Oliveira de Rosso, V., Tura, V., Severo Salau, H., de Oliveira Machado, L., Pimentel Torres, F., Gunski, R.J., and Del Valle Garnero, A. (2025). Atypical presence of interstitial telomeric sequences in Thamnophilus species (Passeriformes: Thamnophilidae). Cytogenet. Genome Res., 1–11.

58. Vollger, M.R., Kerpedjiev, P., Phillippy, A.M., and Eichler, E.E. (2022). StainedGlass: interactive visualization of massive tandem repeat structures with identity heatmaps. Bioinformatics 38, 2049–2051.

59. Tegelström, H., and Ryttman, H. (2009). Chromosomes in birds (Aves): evolutionary implications of macro-and microchromosome numbers and lengths. Hereditas 94, 225–233.

60. Degrandi, T.M., Barcellos, S.A., Costa, A.L., Garnero, A.D.V., Hass, I., and Gunski, R.J. (2020). Introducing the Bird Chromosome Database: An overview of cytogenetic studies in birds. Cytogenet. Genome Res. 160, 199–205.

61. Secomandi, S., Gallo, G.R., Sozzoni, M., Iannucci, A., Galati, E., Abueg, L., Balacco, J., Caprioli, M., Chow, W., Ciofi, C., et al. (2023). A chromosome-level reference genome and pangenome for barn swallow population genomics. Cell Rep. 42, 111992.

62. Habermann, F.A., Cremer, M., Walter, J., Kreth, G., von Hase, J., Bauer, K., Wienberg, J., Cremer, C., Cremer, T., and Solovei, I. (2001). Arrangements of macro- and microchromosomes in chicken cells. Chromosome Res. 9, 569–584.

63. Guiblet, W.M., Cremona, M.A., Cechova, M., Harris, R.S., Kejnovská, I., Kejnovsky, E., Eckert, K., Chiaromonte, F., and Makova, K.D. (2018). Long-read sequencing technology indicates genome-wide effects of non-B DNA on polymerization speed and error rate. Genome Res. 28, 1767–1778.

64. O’Connor, R.E., Farré, M., Joseph, S., Damas, J., Kiazim, L., Jennings, R., Bennett, S., Slack, E.A., Allanson, E., Larkin, D.M., et al. (2018). Chromosome-level assembly reveals extensive rearrangement in saker falcon and budgerigar, but not ostrich, genomes. Genome Biol. 19, 171.

65. Damas, J., Kim, J., Farré, M., Griffin, D.K., and Larkin, D.M. (2018). Reconstruction of avian ancestral karyotypes reveals differences in the evolutionary history of macro- and microchromosomes. Genome Biol. 19, 155.

66. Itoh, Y., and Arnold, A.P. (2005). Chromosomal polymorphism and comparative painting analysis in the zebra finch. Chromosome Res. 13, 47–56.

67. Skinner, B.M., and Griffin, D.K. (2012). Intrachromosomal rearrangements in avian genome evolution: evidence for regions prone to breakpoints. Heredity (Edinb.) 108, 37–41.

68. Knief, U., Hemmrich-Stanisak, G., Wittig, M., Franke, A., Griffith, S.C., Kempenaers, B., and Forstmeier, W. (2016). Fitness consequences of polymorphic inversions in the zebra finch genome. Genome Biol. 17, 199.

69. Viitaniemi, H.M., Leder, E.H., Kauzál, O., Stopková, R., Stopka, P., Lifjeld, J.T., and Albrecht, T. (2024). Impact of Z chromosome inversions on gene expression in testis and liver tissues in the zebra finch. Mol. Ecol. 33, e17236.

70. Knief, U., Forstmeier, W., Pei, Y., Ihle, M., Wang, D., Martin, K., Opatová, P., Albrechtová, J., Wittig, M., Franke, A., et al. (2017). A sex-chromosome inversion causes strong overdominance for sperm traits that affect siring success. Nat. Ecol. Evol. 1, 1177–1184.

71. Itoh, Y., Kampf, K., Balakrishnan, C.N., and Arnold, A.P. (2011). Karyotypic polymorphism of the zebra finch Z chromosome. Chromosoma 120, 255–264.

72. Kim, K.-W., Bennison, C., Hemmings, N., Brookes, L., Hurley, L.L., Griffith, S.C., Burke, T., Birkhead, T.R., and Slate, J. (2017). A sex-linked supergene controls sperm morphology and swimming speed in a songbird. Nat. Ecol. Evol. 1, 1168–1176.

73. Riddle, N.C., and Elgin, S.C.R. (2018). The Drosophila dot chromosome: Where genes flourish amidst repeats. Genetics 210, 757–772.

74. Navarro, F.C.P., and Galante, P.A.F. (2015). A genome-wide landscape of retrocopies in primate genomes. Genome Biol. Evol. 7, 2265–2275.

75. Vollger, M.R., Guitart, X., Dishuck, P.C., Mercuri, L., Harvey, W.T., Gershman, A., Diekhans, M., Sulovari, A., Munson, K.M., Lewis, A.P., et al. (2022). Segmental duplications and their variation in a complete human genome. Science 376, eabj6965.

76. Ko, B.J., Lee, C., Kim, J., Rhie, A., Yoo, D.A., Howe, K., Wood, J., Cho, S., Brown, S., Formenti, G., et al. (2022). Widespread false gene gains caused by duplication errors in genome assemblies. Genome Biol. 23, 205.

77. Henikoff, S., and Furuyama, T. (2010). Epigenetic inheritance of centromeres. Cold Spring Harb. Symp. Quant. Biol. 75, 51–60.

78. Olsson, U., and Alström, P. (2020). A comprehensive phylogeny and taxonomic evaluation of the waxbills (Aves: Estrildidae). Mol. Phylogenet. Evol. 146, 106757.

79. Kinsella, C.M., Ruiz-Ruano, F.J., Dion-Côté, A.-M., Charles, A.J., Gossmann, T.I., Cabrero, J., Kappei, D., Hemmings, N., Simons, M.J.P., Camacho, J.P.M., et al. (2019). Programmed DNA elimination of germline development genes in songbirds. Nat. Commun. 10, 5468.

80. Ruiz-Ruano, F.J., Schlebusch, S.A., Vontzou, N., Moreno, H., Biegler, M.T., Kutschera, V.E., Ekman, D., Borges, I., Pei, Y., Rossini, R., et al. (2025). Programmed DNA elimination drives rapid genomic innovation in two thirds of all bird species. bioRxivorg. 10.1101/2025.07.16.664580.

81. Formenti, G., Koo, B., Sollitto, M., Balacco, J., Brajuka, N., Burhans, R., Duarte, E., Giani, A.M., McCaffrey, K., Medico, J.A., et al. (2025). Evaluation of sequencing reads at scale using rdeval. bioRxivorg. 10.1101/2025.02.01.636073.

82. Stanojevic, D., Lin, D., Florez De Sessions, P., and Sikic, M. (2024). Telomere-to-telomere phased genome assembly using error-corrected Simplex nanopore reads. bioRxiv. 10.1101/2024.05.18.594796.

83. Rautiainen, M., and Marschall, T. (2020). GraphAligner: rapid and versatile sequence-to-graph alignment. Genome Biol. 21, 253.

84. Astashyn, A., Tvedte, E.S., Sweeney, D., Sapojnikov, V., Bouk, N., Joukov, V., Mozes, E., Strope, P.K., Sylla, P.M., Wagner, L., et al. (2024). Rapid and sensitive detection of genome contamination at scale with FCS-GX. Genome Biol. 25, 60.

85. Uliano-Silva, M., Ferreira, J.G.R.N., Krasheninnikova, K., Darwin Tree of Life Consortium, Formenti, G., Abueg, L., Torrance, J., Myers, E.W., Durbin, R., Blaxter, M., et al. (2023). MitoHiFi: a python pipeline for mitochondrial genome assembly from PacBio high fidelity reads. BMC Bioinformatics 24, 288.

86. Saski, M., Ikechi, T., and Makino, S. (1968). A feather pulp culture technique for avian chromosomes, with notes on the chromosomes of the peafowl and the ostrich. Experientia 24, 1292–1293.

87. Barcellos, S.A., de Souza, M.S., Tura, V., Pereira, L.R., Kretschmer, R., Gunski, R.J., and Garnero, A.D.V. (2022). Direct chromosome preparation method in avian embryos for cytogenetic studies: Quick, easy and cheap. DNA 2, 22–29.

88. Ladjali-Mohammedi, K., Bitgood, J.J., Tixier-Boichard, M., and Ponce De Leon, F.A. (1999). International system for standardized avian karyotypes (ISSAK): standardized banded karyotypes of the domestic fowl (Gallus domesticus). Cytogenet. Cell Genet. 86, 271–276.

89. Manni, M., Berkeley, M.R., Seppey, M., and Zdobnov, E.M. (2021). BUSCO: Assessing genomic data quality and beyond. Curr. Protoc. 1, e323.

90. Formenti, G., Abueg, L., Brajuka, A., Brajuka, N., Gallardo-Alba, C., Giani, A., Fedrigo, O., and Jarvis, E.D. (2022). Gfastats: conversion, evaluation and manipulation of genome sequences using assembly graphs. Bioinformatics 38, 4214–4216.

91. Rhie, A., Walenz, B.P., Koren, S., and Phillippy, A.M. (2020). Merqury: reference-free quality, completeness, and phasing assessment for genome assemblies. Genome Biol. 21, 245.

92. Ranallo-Benavidez, T.R., Jaron, K.S., and Schatz, M.C. (2020). GenomeScope 2.0 and Smudgeplot for reference-free profiling of polyploid genomes. Nat. Commun. 11, 1432.

93. Vasimuddin, M., Misra, S., Li, H., and Aluru, S. (2019). Efficient architecture-aware acceleration of BWA-MEM for multicore systems. In 2019 IEEE International Parallel and Distributed Processing Symposium (IPDPS) (IEEE). 10.1109/ipdps.2019.00041.

94. Cheng, Y., Grueber, C., Hogg, C.J., and Belov, K. (2022). Improved high-throughput MHC typing for non-model species using long-read sequencing. Mol. Ecol. Resour. 22, 862–876.

95. Li, H. (2018). Minimap2: pairwise alignment for nucleotide sequences. Bioinformatics 34, 3094–3100.

96. Li, H., Handsaker, B., Wysoker, A., Fennell, T., Ruan, J., Homer, N., Marth, G., Abecasis, G., Durbin, R., and 1000 Genome Project Data Processing Subgroup (2009). The Sequence Alignment/Map format and SAMtools. Bioinformatics 25, 2078–2079.

97. Quinlan, A.R., and Hall, I.M. (2010). BEDTools: a flexible suite of utilities for comparing genomic features. Bioinformatics 26, 841–842.

98. Aksenova, A.Y., and Mirkin, S.M. (2019). At the beginning of the end and in the middle of the beginning: Structure and maintenance of telomeric DNA repeats and interstitial telomeric sequences. Genes (Basel) 10, 118.

99. Goldfarb, T., Kodali, V.K., Pujar, S., Brover, V., Robbertse, B., Farrell, C.M., Oh, D.-H., Astashyn, A., Ermolaeva, O., Haddad, D., et al. (2025). NCBI RefSeq: reference sequence standards through 25 years of curation and annotation. Nucleic Acids Res. 53, D243–D257.

100. NCBI Orthologs: public resource and scalable method for computing high-precision orthologs across eukaryotic genomes, Dong-Ha Oh et al Journal of Molecular Evolution.

101. George E. Liu, Yali Hou*, Twain Brown (2013). Analysis of CR1 repeats in the zebra finch genome. International Institute of Informatics and Cybernetics 11.

102. Cer, R.Z., Bruce, K.H., Mudunuri, U.S., Yi, M., Volfovsky, N., Luke, B.T., Bacolla, A., Collins, J.R., and Stephens, R.M. (2011). Non-B DB: a database of predicted non-B DNA-forming motifs in mammalian genomes. Nucleic Acids Res. 39, D383–D391.

103. Sahakyan, A.B., Chambers, V.S., Marsico, G., Santner, T., Di Antonio, M., and Balasubramanian, S. (2017). Machine learning model for sequence-driven DNA G-quadruplex formation. Sci. Rep. 7, 14535.

104. Goel, M., Sun, H., Jiao, W.-B., and Schneeberger, K. (2019). SyRI: finding genomic rearrangements and local sequence differences from whole-genome assemblies. Genome Biol. 20, 277.

105. Goel, M., and Schneeberger, K. (2022). Plotsr: Visualizing structural similarities and rearrangements between multiple genomes. Bioinformatics 38, 2922–2926.

106. Martin, M. (2011). Cutadapt removes adapter sequences from high-throughput sequencing reads. EMBnet J. 17, 10.

107. Li, H., and Durbin, R. (2009). Fast and accurate short read alignment with Burrows-Wheeler transform. Bioinformatics 25, 1754–1760.

108. Open2C, Abdennur, N., Fudenberg, G., Flyamer, I.M., Galitsyna, A.A., Goloborodko, A., Imakaev, M., and Venev, S.V. (2024). Pairtools: From sequencing data to chromosome contacts. PLoS Comput. Biol. 20, e1012164.

109. Abdennur, N., and Mirny, L.A. (2020). Cooler: scalable storage for Hi-C data and other genomically labeled arrays. Bioinformatics 36, 311–316.

110. Open2C, Abdennur, N., Abraham, S., Fudenberg, G., Flyamer, I.M., Galitsyna, A.A., Goloborodko, A., Imakaev, M., Oksuz, B.A., Venev, S.V., et al. (2024). Cooltools: Enabling high-resolution Hi-C analysis in Python. PLoS Comput. Biol. 20, e1012067.

111. Myers, G., Durbin, R., and Zhou, C. (2025). FastGA: Fast genome alignment. Bioinform. Adv. 10.1093/bioadv/vbaf238.

112. Umu, S.U., Paynter, V.M., Trondsen, H., Buschmann, T., Rounge, T.B., Peterson, K.J., and Fromm, B. (2023). Accurate microRNA annotation of animal genomes using trained covariance models of curated microRNA complements in MirMachine. Cell Genom. 3, 100348.

113. Oksanen, J., Simpson, G.L., Blanchet, F.G., Kindt, R., Legendre, P., Minchin, P.R., O’Hara, R.B., Solymos, P., Stevens, M.H.H., Szoecs, E., et al. (2001). Vegan: Community ecology package. (The R Foundation). 10.32614/cran.package.vegan 10.32614/cran.package.vegan.

114. Clarke, A.W., Høye, E., Hembrom, A.A., Paynter, V.M., Vinther, J., Wyrożemski, Ł., Biryukova, I., Formaggioni, A., Ovchinnikov, V., Herlyn, H., et al. (2025). MirGeneDB 3.0: improved taxonomic sampling, uniform nomenclature of novel conserved microRNA families and updated covariance models. Nucleic Acids Res. 53, D116–D128.

115. Lloyd, G.T. (2016). Estimating morphological diversity and tempo with discrete character-taxon matrices: implementation, challenges, progress, and future directions. Biol. J. Linn. Soc. Lond. 118, 131–151.

116. Lloyd, G.T. (2018). Journeys through discrete-character morphospace: synthesizing phylogeny, tempo, and disparity. Palaeontology 61, 637–645.

117. Lehmann, O.E.R., Ezcurra, M.D., Butler, R.J., and Lloyd, G.T. (2019). Biases with the Generalized Euclidean Distance measure in disparity analyses with high levels of missing data. Palaeontology 62, 837–849.

118. Conceição, H.B., Mercuri, R.L.V., de Castro, M.P.M., Ohara, D.T., Guardia, G.D.A., and Galante, P.A.F. (2024). RCPedia: a global resource for studying and exploring retrocopies in diverse species. Bioinformatics 40. 10.1093/bioinformatics/btae530.

119. Pertea, G., and Pertea, M. (2020). GFF utilities: GffRead and GffCompare. F1000Res. 9, 304.

120. Kiełbasa, S.M., Wan, R., Sato, K., Horton, P., and Frith, M.C. (2011). Adaptive seeds tame genomic sequence comparison. Genome Res. 21, 487–493.

121. Miller, T.L.A., Conceição, H.B., Mercuri, R.L., Santos, F.R.C., Barreiro, R., Buzzo, J.L., Rego, F.O., Guardia, G., and Galante, P.A.F. (2023). Sandy: A user-friendly and versatile NGS simulator to facilitate sequencing assay design and optimization. bioRxiv. 10.1101/2023.08.25.554791.

122. Bray, N.L., Pimentel, H., Melsted, P., and Pachter, L. (2016). Near-optimal probabilistic RNA-seq quantification. Nat. Biotechnol. 34, 525–527.

123. Dobin, A., Davis, C.A., Schlesinger, F., Drenkow, J., Zaleski, C., Jha, S., Batut, P., Chaisson, M., and Gingeras, T.R. (2013). STAR: ultrafast universal RNA-seq aligner. Bioinformatics 29, 15–21.

124. Altschul, S.F., Gish, W., Miller, W., Myers, E.W., and Lipman, D.J. (1990). Basic local alignment search tool. J. Mol. Biol. 215, 403–410.

125. Benson, G. (1999). Tandem repeats finder: a program to analyze DNA sequences. Nucleic Acids Res. 27, 573–580.

126. Katoh, K., and Standley, D.M. (2013). MAFFT multiple sequence alignment software version 7: improvements in performance and usability. Mol. Biol. Evol. 30, 772–780.

127. Eddy, S.R. (2011). Accelerated profile HMM searches. PLoS Comput. Biol. 7, e1002195.

