## Supplementary Information for "The complete genome of a songbird"

**Index**

1. Gfalign, a tool to assist T2T manual graph curation
2. Completion of the Z chromosome
3. Nature of telomeres
4. Chromatin conformation annotation
5. Gene enrichment
6. Retrocopies
7. MicroRNAs

### 1. Gfalign, a tool to assist T2T manual graph curation

Gfalign (https://github.com/vgl-hub/gfalign), part of the gfastar tool suite[1,2], is a graph-aware tool designed to facilitate the manual curation of telomere-to-telomere (T2T) genome assemblies by analyzing assembly graph topology and read-to-graph alignments generated by tools that can map long-read sequencing data onto assembly graphs (e.g. GraphAligner[3]). The tool is intended to work with the assembly graphs produced by Verkko[4] or other T2T pipelines. Gfalign computes alignment summary statistics and decorates the graph with information from the alignment, such as annotating graph nodes with read depth information. Alignments are processed to assess read support for different paths within the graph, helping to distinguish between true genomic structures and potential assembly artifacts. gfalign enables visualization of read alignments in a graph context, allowing curators to interactively examine ambiguous regions, such as repetitive elements or segmental duplications, breakpoints, collapsed repeats, or misoriented contigs. Gfalign can also filter alignments based on quality and coverage.

For the T2T zebra finch genome project, gfalign was used to decorate the graph with edge coverage support from ultralong reads. This enables manual curators to decide the most likely paths in ambiguous bubbles. Additionally, gfalign also provides a way to automatically solve complex tangles by performing a constrained search for the optimal path between two nodes in the assembly graph. This was implemented as a A* algorithm with the two nodes as start and end goals. The search uses a heuristic based on the support provided by reads aligned to edges. This was tested to generate model sequences for the two complex tangles identified on the Z chromosome, which ultimately led to the development of an even more optimized strategy, as described in the next section.

### 2. Completion of the Z chromosome

The TTT tool (https://github.com/marbl/TTT) was used for resolving two tangles on the Z chromosome (Antipov et al., in preparation). Briefly, this tool finds a traversal through genomic tangles in the assembly graph using ONT read alignments and coverage information. First, TTT estimates median coverage *med_cov* of unique heterozygous nodes in the graph. Then it finds *multiplicities* of each node N in a tangle, such that coverage(N)/multiplicity(N) is as close to *med_cov* as possible and there exists a path through the tangle that contains each node N exactly multiplicity(N) times. This is done with mixed integer linear programming optimization. Then among all paths with given node multiplicities TTT searches for the one with best match to the read alignment through gradient descent-like optimization.

#### 3. Nature of telomeres

Comparative analysis of telomeres revealed slight, non-significant differences by haplotype and by chromosome arms. Paternal telomeres averaged 10.62 ± 4.42 Kbp (median 9.34 Kbp, range 0.95-23.24 Kbp), while maternal telomeres averaged 12.16 ± 5.54 Kbp (median 10.57 Kbp, range 3.23-30.34 Kbp). The variation observed likely arises from specific chromosomal extremes, such as the notably large maternal chrW q-arm telomere and the exceptionally short paternal chr34 p-arm telomere. Terminal telomere lengths in Zebra finches show very high heritability ($h^2 \approx 0.99$, 95% CI 0.87-1) with minimal influence from the environment[5]. The comparable telomere lengths between maternal and paternal haplotypes and across chromosome arms suggest balanced inheritance mechanisms.

#### 4. Chromatin conformation annotation

Across all multiple resolutions tested, a positive correlation was found between the first eigenvector (E1) and gene density for all chromosomes at 200 Kbp and 100 Kbp. However, at 10 kbp, both paternal chr33 and maternal chr34 showed inverse relationships, likely due to sparse Hi-C interaction data. In contrast, there was a negative correlation between E1 and GC content in chr16, chr21, chr36, and chrW at 200 Kbp resolution. Therefore, we found the gene density track to be a much better classifier than GC content in both high and low resolutions.

#### 5. Gene enrichment

Gene enrichment analysis of the duplicated regions reveals a marked expansion of olfactory receptor (OR) gene families in the zebra finch genome. Several distinct OR gene families are represented, indicating a broad diversification of the OR repertoire (e.g., OG0000000). In particular, 122 copies of the 14J1-like family were identified, alongside 58 of the 14A16-like, 29 of the 14C36-like, and 9 of the 14I1-like (with additional pseudogenes: 56, 15, 7, and 2, respectively). Moreover, the 52H1-like OR family is represented solely by a single pseudogene. OR genes are even annotated with the GO term "*G protein-coupled receptor activity*", a functional category also shared by taste receptor genes. However, taste receptor family proteins are included in different orthogroups (OG0000040, OG0000323, OG0001647, OG0012326), and they do not appear to be expanded in the zebra finch, despite the presence of nine functional copies and one pseudogene detected. The GO term "*histone H3K4 methyltransferase activity*" is significantly enriched within duplicated regions of the genome, largely due to duplications of the SETD1A gene. These duplications are mainly found on micro-dot chromosomes, with copy numbers distributed as follows: four on chromosome 16, two on chromosome 32, one on chromosomes 33 and 34, and a peak of five copies on chromosome 35. This gene family (included in two different orthogroups OG0000396, OG0014197) also shows a broader expansion in the zebra finch genome, pointing to a lineage-specific amplification of chromatin regulatory functions. Another enriched term is the cellular component "*MHC class II protein complex*", with corresponding genes distributed across multiple chromosomes, but showing a strong concentration on chromosome 35 (14 copies). This pattern suggests a localized expansion and potential functional specialization (OG0000022). The term "*protein phosphorylation*" is also significantly enriched. This class includes multiple kinase families, as the p21-activated kinases (PAK3-like, or OG0000002), with 30 copies overall, which are strongly present in chromosome Z and appear to be more expanded in Oscines than Suboscines (56 vs. 6

copies). Similarly, PIM1, PIM2, and PIM3-like genes, a family of serine/threonine protein kinases, are strongly expanded in Oscines (OG0000004, OG0000023 with a copy number of around 42 and 7 respectively). In particular, PIM3-like copies are predominantly located on microchromosomes 25, 29, 30, 31, 33, and 34. The GO term "*viral process*" is also enriched within duplicated regions, largely driven by endogenous retroviral elements. This includes OG0000033 (ERV-K group) and OG0000179 (C15orf39-like protein). Notably, the latter exhibits a tandem expansion of 12 copies within a ~2 Mbp region on chromosome 10. The molecular function "*scavenger receptor activity*" is associated with OG0000089, which is significantly enriched in duplicated regions. While this orthogroup shows an average of 9.0 copies in Suboscines versus 2.81 in Oscines, the zebra finch exhibits a remarkable expansion, with 11 functional copies and one pseudogene identified in the TaeGut7 assembly. Lastly, the GO term "*sequence-specific DNA binding*", which includes heat shock transcription factors, is also enriched among duplicated genes. One representative orthogroup, OG0000051, shows an average of 6.0 copies in Oscines and 4.0 in Suboscines, but it is notably expanded in the zebra finch, with 24 copies in the TaeGut7 genome, reflecting possible adaptation in transcriptional regulation.

### 6. Retrocopies

#### Retrocopies identified in maternal and paternal zebra finch assemblies

In addition to DNA-mediated gene duplication (unprocessed pseudogenes, or simply pseudogenes; Fig. 6B), we investigated mRNA-mediated duplication, which gives rise to processed pseudogenes, or retrocopies[6], in the maternal and paternal genomes. Using an RCPedia-based pipeline[7], retrocopies were cataloged independently in the phased assemblies. On autosomes, we identified 78 retrocopies, with extensive overlap between assemblies (96.2%): 77 in the maternal and 76 in the paternal genome (**Supplementary Figure 20A**). Beyond autosomes, we observed a significant enrichment on the sex chromosomes ($\chi^2(1, N = 107) = 8.72$, $p = 0.0031$; Supplementary Fig. 20B), with 16 copies on Z and 13 on W (**Supplementary Tables 14–15**).

Several parental genes exhibited recurrent retroposition. In the maternal assembly, LOC140680666 (14 copies) and LOC100231660 (8) were most prolific, followed by DNAJC15 (5) and LOC100228280 (4). In the paternal assembly, LOC140680666 (5) and DNAJC15 (4) were major contributors, alongside LOC100228280 (4), LOC100231660 (3), TRERF1 (3), and SP1 (3) (**Supplementary Table 16**). Full rankings are provided in **Supplementary Table 14**. Notably, 50 retrocopies inserted within annotated protein-coding genes (host genes), including ACTA2P1 in ACTB, ACTBP1 in ACTA1, ACTBP2 in ACTC1, and ACTC1P1 in ACTG1 (**Supplementary Table 17**). These insertions provide a framework for downstream analyses of potential regulatory disruption or exonization.

Chromosomal distribution analyses revealed a strong correlation between retrocopy number and chromosome size, with mild enrichment on dot chromosomes and depletion on microchromosomes. The sex chromosomes were clear outliers, showing marked enrichment relative to this trend ($\chi^2(1, N = 107) = 8.72$, $p = 0.0031$; **Supplementary Figure 21**). Overall, retrocopies were broadly distributed, with multiple inter-chromosomal events, recurrent donor loci, and pronounced involvement of the sex chromosomes (**Supplementary Figure 22**).

#### Sex-chromosome distribution of retrocopies in zebra finch

Among the 29 retrocopies on sex chromosomes, 16 reside on Z (15% of retrocopies in the maternal assembly) and 13 on W (12%; **Supplementary Figure 23A**; **Supplementary Tables 18–19**). On Z, 13 retrocopies originated from autosomal genes and three from Z-linked genes. Additionally, one Z-linked parental gene produced a retrocopy on an autosome (**Supplementary Table 14**). Given the size, gene density, and hemizygous state of Z in females (ZW), dosage-compensation pressures likely shape its retrocopy landscape. The predominance of autosomally derived Z-linked retrocopies (81.2%; 13/16) suggests that Z may serve as a receptive site for insertions that buffer or modulate dosage-sensitive genes, especially in females. The three Z-derived retrocopies may instead reinforce expression, alter regulation, or provide sexually antagonistic variants. The observation of a Z-derived retrocopy inserted on an autosome (FSD1LP1 from FSD1L, chrZ to chr3) further illustrates the dual role of Z as both source and sink. FSD1LP1 is expressed in muscle and testis (**Supplementary Figure 23**), providing a candidate example of an "out-of-Z" dosage-balancing mechanism[8], in which an autosomal copy mitigates haploinsufficiency in females.

Expression analyses detected activity in 10 of 16 Z-linked retrocopies (62.5%), predominantly testis-biased (9), with additional cases in ovary (4) and brain (4). This pattern supports the well-known testis permissive transcriptional environment and suggests retention of retrocopies contributing to sex-biased or germline functions.

By contrast, W-linked retrocopies showed a distinct pattern. All 13 derived from autosomal parental genes, consistent with the advanced degeneration and low transcriptional output of W. Most were silent (10/13; 76.9%) across surveyed tissues (brain, muscle, ovary, testis), supporting a "graveyard" model in which W acts largely as a passive sink. However, three W-linked retrocopies were expressed, including RAB7AP1 (derived from RAB7A), which was active in ovary and brain. These cases suggest a limited "colonization" model, in which a subset of autosome-derived insertions on W acquires female-biased regulation. Together, the coexistence of largely silent W-linked retrocopies with a few expressed examples points to W as both a repository and occasional incubator of novelty. Our analyses are limited to four tissues, and additional profiling across developmental stages and deeper sequencing will likely uncover further expression. Direct functional assays (e.g., ribosome profiling, CRISPR perturbations) will be critical to assess coding potential and regulatory impact, particularly for candidates such as FSD1LP1 and RAB7AP1.

Because gene retroduplication is a major source of genomic innovation, the continued rollout of telomere-to-telomere (T2T) assemblies across avian lineages will: i) enable more accurate estimates of retrocopy burden and turnover; ii) allow rigorous tests of Z/W "source–sink," "graveyard," and "colonization" dynamics; and iii) provide high-resolution insights into the interplay between retroposition, dosage compensation, and retroduplication-associated phenomena.

### 7. MicroRNAs

Across 132 bird species, MirMachine identified 116 unique miRNA families[9] of which 67 were present in every species (**Supplementary Table 20**). NMDS clustering based on the presence/absence matrix of miRNA families shows a distinct clustering of passerine vs non-passerine birds (**Supplementary Figure 25A**), with the zebra finch clustering well within passerine birds. The NMDS distance of zebra finch from the centroid of all passerine species (*i.e.* the hypothetical 'typical' passerine miRNA repertoire) was close to the centre of values across all passerine species, indicating that the zebra finch is no more diverged from this typical ‘passerine miRNA repertoire’ than passerine species generally (**Supplementary Figure 25B**). Of the 15 miRNA families identified from literature, 11 are present in every bird genome (*i.e.* Passerine and non-passerine birds). Of the remaining 4 miRNAs (MIR-132, -140, -192 & -2954) only MIR-2954, a microRNA with specific expression in parts of the zebra finch brain associated with song perception and production[10], shows particular association with Passeriformes, having been lost 4 times across birds, but never in passerine species (**Supplementary Figure 26A**). Independent of previous literature, two other microRNA families (MIR-459 and MIR-737) were lost in the passerine ancestor, but were also widely lost in other bird species (**Supplementary Figure 26B,C**). The pattern suggests that the loss of MIR-459 and MIR-737 may have had an important role in the evolution of passerine life history which is not currently characterised from experimental studies.

### 8. TIGD4 as CENP-B functional candidate in Zebra finch: Evidence for Convergent Evolution

He follows evidence that TIGD4, a domesticated pogo/Tigger family transposase, may serve as a functional CENP-B homolog in birds. Structural analysis revealed remarkable similarity between zebra finch TIGD4 and human CENP-B, with conservation of critical DNA-binding domains (45% identity in CENP-B domain, 86-90% conservation between human and avian TIGD4). Interestingly, TIGD4 completely lacks the C-terminal dimerization domain of CENP-B, putatively having evolved an alternative oligomerization mechanism through its central region.

Functional evidence includes: (1) TIGD4 expression pattern characteristic of centromeric proteins— approximately 2.5-fold higher in proliferating tissues (testis) versus post-mitotic tissues (brain); (2) Identification of CENP-B box-like motifs in centromeric repeats across all avian chromosomes and AlphaFold3 modeling confirmed the formation of functional TIGD4-dimer/CENP-C/DNA complexes analogous to mammalian centromeric architecture.

Evolutionary implications suggest convergent transposon domestication for centromeric function: mammals utilize CENP-B with C-terminal dimerization, while birds may employ TIGD4 with alternative dimerization, yet both systems retain capacity for CENP-B/TIGD4 heterodimer formation. Conserved synteny of TIGD4 across birds, amphibians, and mammals indicates ancient origin followed by functional divergence. These findings suggest that the absence of CENP-B in birds, rather than making centromere formation epigenetic, may be compensated by a functionally equivalent but structurally divergent protein, resolving the paradox of centromere stability without canonical CENP-B and revealing unexpected plasticity in vertebrate centromere evolution.

*Discovery by human CENP-B similarity*

The CENP-B protein evolved specifically within the mammalian lineage from a domesticated pogo-like transposon and is absent in non-mammalian genomes, including birds[11]. However, organisms like the yeast Saccharomyces cerevisiae have functional analogs, such as the Centromere Binding Factor 1 (Cbf1) protein. Although Cbf1 also binds to specific DNA sequences within the centromere, it shares no evolutionary origin with the mammalian CENP-B, highlighting the general possibility of convergent evolution for centromere-binding functions[12]. This raised a fundamental question: although birds possess key centromeric proteins such as CENP-A and CENP-C, they appear to lack CENP-B entirely. Given the well-established role of CENP-B in directly binding centromeric DNA and facilitating the deposition of CENP-A into centromeric nucleosomes in mammals, we asked — what, if anything, performs this function in birds? To begin addressing this question, we conducted an initial experiment in which we used the human CENP-B protein (NP_001801) as a query in a BLAST search against the transcriptome of the zebra finch (Taeniopygia guttata), a model avian species.

A nucleotide tblastn search (human CENP-B protein queried against the translated zebra finch transcriptome) revealed three significant hits in the zebra finch (Taeniopygia guttata) genome, all corresponding to tigger transposable element-derived (TIGD) genes:

TIGD4 (XM_072928462.1): a mRNA (length 1,781 nt) as the top hit (Max/Total Score = 142; $E = 5\times10^{-35}$). The alignment covers 57% of the query and shows 28.8% amino-acid identity. Although the percent identity is modest, the extremely low E-value together with the alignment length indicates genuine homology driven by extended, collinear similarity across the coding region rather than short matches by chance.

TIGD3 (XM_030266919.4): Annotated as LOC115494068, this sequence demonstrated comparable alignment metrics (Max Score: 137, Total Score: 137) with 64% query coverage and 28.61% identity. The E-value of 4e-33 confirms significant homology. This transcript spans 2,706 nucleotides.

TIGD5 (XM_030266836.4): While showing lower overall alignment scores (Max Score: 83.2, Total Score: 83.2), this sequence maintained similar percent identity (28.92%) despite reduced query coverage (41%). The E-value of 1e-15 still indicates significant similarity. The transcript length is 2,929 nucleotides.

All three sequences represent predicted mRNA transcripts encoding proteins derived from tigger transposable elements, suggesting potential evolutionary conservation or functional significance of these mobile genetic element-derived sequences in the zebra finch genome. The consistent ~28-29% sequence identity across all three hits indicates divergent but related sequences, typical of transposable element-derived genes that have undergone domestication and subsequent evolutionary divergence.

*TIGD4 and CENP-B from one Group 1 pogoR-derived*

The TIGD protein family can be categorized based on their transposase ancestry into three distinct evolutionary groups[11]. Group I, derived from pogoR transposases, includes TIGD3, TIGD4, TIGD6, and CENP-B. Group II, derived from Tigger elements, comprises CENPBD1, JRK, JRKL, TIGD2, TIGD5, and TIGD7. Group III, originating from Passer transposons, includes POGK and POGZ[11]. This classification reflects independent domestication events and suggests functional diversification following transposon co-option. Most TIGD-derived proteins retain core features of their transposase ancestors, such as DNA-binding domains (DBDs) and DDE catalytic domains. However, several have acquired additional domains that facilitate specialized functions. For example, CENP-B has evolved a dimerization domain essential for centromere binding and artificial chromosome formation. POGK has acquired a KRAB repression domain, and POGZ contains multiple zinc finger motifs. Functionally, these proteins have been implicated in diverse biological processes: POGZ plays a role in chromatin remodeling and is associated with autism spectrum disorders; JRK has been linked to epilepsy and cancer; and CENP-B remains a critical component in centromere identity and artificial chromosome assembly. Together, these examples highlight the functional innovation that can emerge from transposase domestication.

*TIGD4 is located in synteny region conserved between zebra finch and other vertebrates*

Comparative genomic analysis revealed remarkable syntenic conservation of the TIGD4-containing region between human and zebra finch genomes (**Supplementary Figure 12**). In humans, TIGD4 is located on chromosome 12 (152,545,700-153,028,083) within a gene-dense region flanked by the ribosome biogenesis factor NSA2 and other highly conserved genes. The orthologous region in zebra finch (chromosome NC_133028.1: 142,820-142,880 kb) maintains identical gene order and orientation, with TIGD4 (XM_072928462.1) positioned between the same flanking genes ARFIP1 and TMEM154.

This syntenic conservation spanning ~300 million years of avian-mammalian divergence indicates strong purifying selection acting on this genomic locus. RNA-sequencing data from both species demonstrate robust TIGD4 expression, with particularly high levels observed in specialized tissues (**Supplementary Figure 13**). In zebra finch, intron-spanning reads reach up to 5,080 counts, confirming active transcription. In humans, TIGD4 shows notable overexpression in the retina—a tissue characterized by distinctive chromatin architecture compared to other somatic cell types.

The preservation of both genomic context and tissue-specific expression patterns strongly suggests that TIGD4 performs an essential, evolutionarily conserved function. While its precise role has not yet been described, the combination of deep conservation, syntenic maintenance, and enriched expression in tissues with specialized nuclear organization requirements points to a fundamental biological importance that has been maintained across vertebrate evolution. This raises the intriguing possibility that TIGD4 may serve critical, yet currently unrecognized, functions in both avian and mammalian genomes, potentially related to chromatin organization or specialized gene regulatory mechanisms.

*Structural similarity between CENP-B and TIGD4*

Following sequence-based identification, we proceeded to investigate the structural characteristics of the identified TIGD proteins. Given that proteins with low sequence identity can exhibit conserved three-dimensional architectures and maintain functional similarity, we performed comprehensive structure-based alignments of the predicted protein products from the three zebra finch transcripts.

*Structural Homology Results*

The structural alignment analysis revealed distinct patterns of similarity for each TIGD protein.

TIGD4 Structural Analysis: The predicted TIGD4 protein structure demonstrated significant alignment with the crystal structure of human CENP-B (PDB: 1HLV), specifically the N-terminal DNA-binding domain (residues 1-129) complexed with CENP-B box DNA. This structural similarity suggests that TIGD4 may retain DNA-binding capabilities analogous to CENP-B, despite the modest sequence identity observed in our initial BLAST analysis.

TIGD3 Structural Analysis: Structural alignment of TIGD3 revealed homology with two zinc finger-containing proteins: Human PRDM9 allele-A zinc finger domain in complex with its associated recombination hotspot DNA sequence (PDB: 5EGB). The engineered six-finger zinc finger protein Aart, designed for recognition of ANN triplet sequences (PDB: 2I13). These alignments indicate that TIGD3 likely possesses zinc finger domains with DNA-binding potential, though the specific DNA recognition sequences may differ from canonical CENP-B targets.

TIGD5 Structural Analysis: TIGD5 showed structural similarity to the solution structure of a CENP-B N-terminal DNA-binding domain from Drosophila (PDB: 2ELH), derived from the distal antenna protein CG11849-PA. This alignment suggests evolutionary conservation of CENP-B-like structural features across diverse species, extending from arthropods to avians.

The structural alignments demonstrate that despite relatively low sequence conservation (~28-29% identity), all three TIGD proteins maintain structural features characteristic of DNA-binding proteins, with TIGD4 and TIGD5 showing direct structural homology to CENP-B domains. This finding supports the hypothesis that these transposable element-derived proteins may have retained or evolved DNA-binding functions potentially relevant to centromeric or heterochromatic regions in the zebra finch genome.

*Domain level similarity between CENP-B and TIGD4*
To further investigate domain-level conservation, we conducted a detailed alignment between human CENP-B, human TIGD4, and the zebra finch TIGD4 homolog (Supplementary Figure 9). We annotated five key functional domains in CENP-B: the N-terminal DNA-binding domain (CENP-B_N, residues 2–56), a broader DNA-binding region (1–125), a putative centromere-specific DNA-binding domain (CENP-B, 74–135), a DDE superfamily catalytic domain (DDE_1, 222–384), and the CENP-B dimerization domain (539–598) (Gao et al., 2020).

Strikingly, these domains were at least partially identifiable in TIGD4. Despite relatively modest sequence identity between CENP-B and TIGD4 (30.4 % for full-length human proteins), domain-level similarity was notable. For example, the DNA-binding domains showed 36% identity and 46.4% similarity, and even the DDE_1 domain retained approximately 43% similarity. The dimerization domain showed lower identity (21.7%) and was markedly reduced in the zebra finch homolog (15% identity, 28.3% similarity), though still detectable.

These results challenge the assumption that CENP-B contains unique functional domains absent in other TIGD proteins. Instead, TIGD4 appears to harbor the full complement of core CENP-B domains, suggesting functional potential that may have been underestimated. Notably, conservation between the human and zebra finch TIGD4 orthologs was high across all domains, further supporting functional relevance in avian species.

Our comparative domain-level analysis of CENP-B and TIGD4 across species revealed several striking evolutionary patterns, providing insights into functional divergence and conservation. It was previously shown that the CENP-B dimerization domain (residues 539–598) is almost entirely absent in TIGD4[11]. Sequence identity between human CENP-B and TIGD4 in this region is 21.7%, and 63% in this region was observed between the human and zebra finch TIGD4 orthologs. These findings strongly suggest that TIGD4 has partially lost the canonical dimerization mechanism used by CENP-B. Despite overall sequence divergence, the DNA-binding regions are remarkably conserved. The general DNA-binding domain (residues 1–125) shows approximately 36% identity between CENP-B and TIGD4, which is high for transposon-derived proteins. The more specific CENP-B domain (74–135) exhibits even higher identity (43.6%). Between human and zebra finch TIGD4, identity in these regions is 90%, highlighting strong purifying selection and suggesting retained DNA-recognition capability.

The DDE_1 domain (residues 222–384), derived from the ancestral transposase catalytic core, displays 29.2% identity between CENP-B and TIGD4, and 66.7% identity between TIGD4 orthologs. While likely catalytically inactive for transposition in TIGD4, the domain may retain structural or scaffolding roles. This pattern supports a scenario in which DNA-binding function was conserved, while the dimerization interface was lost, possibly replaced by alternative interaction strategies.

TIGD4 evolves more slowly across species (e.g., 67.3% identity between human and zebra finch TIGD4) than the divergence observed between CENP-B and TIGD4 (28–30%), suggesting strong purifying selection on TIGD4 after its divergence from CENP-B. These data quantitatively support the model in which TIGD4 represents a functional adaptation of the CENP-B architecture, conserving its DNA-binding role while diverging significantly in oligomerization strategy. This is consistent with the broader hypothesis of convergent evolution in centromere assembly mechanisms.

*Similarly to CENP-B, TIGD4 can form dimer*

Among the candidate transcripts, the TIGD4 homolog emerged as the closest sequence match to human CENP-B. We selected TIGD4 for further analysis and modeled its structure using AlphaFold 3, alongside a model of human CENP-B for direct comparison. Structural predictions were performed for both monomeric and dimeric forms. While it has been suggested that TIGD4-family proteins may not form stable dimers[11], our AlphaFold-based modeling clearly indicated that the zebra finch TIGD4 homolog is capable of forming a dimeric structure, similar to CENP-B. This observation challenges prior assumptions and supports the hypothesis that TIGD4 may functionally compensate for the absence of CENP-B in birds.

Our structural modeling of TIGD4 and CENP-B dimers yielded striking insights into their respective dimerization mechanisms and evolutionary trajectories. Similar to human CENP-B (**Supplementary Figure 10A**), he predicted TIGD4 homodimer revealed well-defined interdomain contacts, as visualized in the Predicted Aligned Error (PAE) matrix, with strong signal around residues ~540-576 (**Supplementary Figure 10B**). This indicates that TIGD4 forms a stable homodimer, through its C-terminal region similarly to the canonical C-terminal dimerization domain used by CENP-B. The CENP-B homodimer exhibited the expected architecture, with clear dimerization through the C-terminal domain (residues ~540-599). Most unexpectedly, we observed that TIGD4 and CENP-B are also capable of forming a heterodimer. While the confidence scores for the heterodimeric model (ipTM = 0.25, pTM = 0.31) suggest moderate reliability, the predicted structure is plausible, and the PAE map shows distinct inter-protein contact regions. These results point to a previously unrecognized dimerization mechanism in TIGD4[11], and reveal that TIGD4 utilizes a dimerization mechanism analogous to CENP-B rather than a distinct strategy. The shared reliance on C-terminal domains for oligomerization, coupled with the unexpected capacity for heterodimer formation, suggests evolutionary conservation of this functional architecture despite sequence divergence in protein regions. The ability of TIGD4 to form both homodimers and heterodimers with CENP-B (**Supplementary Figure 10C**) suggests the possibility of combinatorial dimer configurations in species where both proteins are present, potentially reflecting transitional evolutionary states. In birds, where CENP-B is absent, TIGD4 may function autonomously, with dimerization enabling recognition of palindromic or tandem repeat DNA motifs—features characteristic of centromeric sequences. These findings collectively corroborate the hypothesis that TIGD4 may represent a structurally adapted, functionally competent replacement for CENP-B in avian species, and further modeling of its interaction with centromeric DNA will be instrumental in elucidating its precise role in centromere organization.

*Structural alignment of CENP-B dimer and TIGD4 dimer*

Upon closer inspection, we observed that while TIGD4 and CENP-B share a highly conserved central core, their terminal regions differ substantially. Specifically, TIGD4 possesses an extended N-terminal region that is absent in CENP-B, whereas CENP-B features a long C-terminal extension not present in TIGD4. These terminal regions do not align structurally, suggesting they are divergent. The presence of a conserved central domain flanked by distinct termini in each protein suggests a possible domain rearrangement or internal translocation event during evolution. This structural mosaicism raises intriguing questions about the modular evolution and functional divergence of these centromere-associated proteins.

*TIGD4 can bind CENP-C in silico as CENP-B*

As a next step, we incorporated CENP-C into the analysis and introduced centromeric DNA sequences predicted by our centromere-identification pipeline (Tgut716A). Our aim was to assess whether these components could assemble into a coherent complex. Remarkably, the modeling confirmed that complex formation is indeed possible. We successfully constructed in silico centromeric complexes for both Homo sapiens using human CENP-B, CENP-C, and alphoid centromeric DNA (**Supplementary Figure 10D**) and Taeniopygia guttata (zebra finch) using its endogenous proteins (**Supplementary Figure 10E**). These models provide further structural support for the hypothesis that the avian centromere machinery, although lacking canonical CENP-B, may functionally compensate through alternative protein assemblies involving TIGD4.

*Predicted TIGD4 transcription in meiosis*

We performed transcriptome analysis on RNA-seq data from NCBI BioProject PRJNA768106[13] by aligning reads to the paternal haploid zebra finch genome with STAR and quantifying gene counts using featureCounts; differential expression was then assessed with DESeq2 both between brain and testis and among three temperature conditions (27 °C, 35 °C and 43 °C) for each tissue. We found that TIGD4 is significantly more highly expressed in testis than in brain, with a $\log_2$ fold change (MLE; brain vs testis) of –2.49526, a Wald test p-value of $9.72\times10^{-69}$ (padj = $3.02\times10^{-68}$), baseMean = 130.444, lfcSE = 0.1424 and Wald statistic = –17.5221, a classical hallmark of centromeric proteins. In contrast, no significant differences in TIGD4 expression were observed across the 27 °C, 35 °C and 43 °C conditions in either brain or testis samples.

*Zebra Finch has TIGD4 box enriched in satellite DNA*

Letter height in Supplementary Figure 11 reflects positional conservation (information content, 0–2 bits). The CENP-B box shows a sharply conserved core (~15–23) with an A-rich stretch followed by GGG, consistent with strong purifying selection on binding-critical positions. The zebra finch TIGD4 box also displays a well-defined, G-rich core (~13–23) and conserved T's at ~7–8, but lower conservation in the flanks. Overall, CENP-B boxes are highly constrained, whereas TIGD4 boxes in birds exhibit focused constraint at the core with more flexible flanking positions.

*Evolutionary competition between pogo/Tigger-derived proteins reveals functional replacement of TIGD4 by CENP-B in primates*

Both CENP-B and TIGD4 belong to the pogo/Tigger transposase-derived protein family, but show markedly different phylogenetic distributions. While TIGD4 is broadly conserved across vertebrates, CENP-B emerged as a mammalian innovation, presenting a unique opportunity to study functional evolution of domesticated transposases[11]. Our analysis reveals a striking pattern consistent with functional replacement. In zebra finch, where CENP-B is absent, the TIGD4 box displays high information content and clear sequence conservation (Supplementary Figure 11, bottom), indicating strong purifying selection maintaining this motif. Conversely, in humans where both proteins coexist the TIGD4 box shows dramatically reduced information content despite our relaxed search criteria (edit distance ≤5), while the CENP-B box maintains sharp conservation with high information content.

This pattern supports a model of competitive exclusion where the newly evolved CENP-B has progressively displaced ancestral TIGD4 functions in primates. The quantitative distribution data further supports this hypothesis: CENP-B boxes vastly outnumber TIGD4 boxes across most human chromosomes, with some showing 6-fold enrichment (e.g., chromosome 18). The relaxed selection on human TIGD4 boxes, evidenced by their degenerate sequence logo, suggests ongoing functional decay following replacement by CENP-B. Notably, chromosome 2—the only human autosome where TIGD4 boxes outnumber CENP-B boxes—may represent a genomic refuge where ancestral TIGD4 function persists. Similarly, the retention of TIGD4 boxes on the Y chromosome (942 occurrences) with near-complete absence of CENP-B boxes (10 occurrences) might reflect chromosome-specific constraints preventing CENP-B invasion. This evolutionary scenario exemplifies how gene duplication and neofunctionalization events can lead to functional turnover, with newer paralogs potentially outcompeting and replacing ancestral functions—a process particularly evident in the rapid evolution of centromeric and heterochromatic DNA-binding proteins derived from transposable elements.

*TIGD4 boxes show centromere-specific localization in zebra finch genome*

To test whether TIGD4 functionally replaces CENP-B in non-primate vertebrates, we examined the chromosomal distribution of TIGD4 boxes in the zebra finch genome. Strikingly, TIGD4 boxes showed highly non-random distribution with strong enrichment at specific chromosomes known to harbor centromeric satellite DNA The W chromosome displayed the highest TIGD4 box density (1,585 occurrences), consistent with its heterochromatic nature and enriched satellite content. Several autosomes showed similarly high enrichment, including chromosomes 8 (>1,000 boxes per haplotype), 13_pat (1,008 boxes), and 25 (973 maternal, 593 paternal). In contrast, most microchromosomes (chr14-28) contained fewer than 100 TIGD4 boxes per haplotype, despite their substantial combined genomic length. Crucially, our analysis revealed that TIGD4 boxes in zebra finch colocalize almost exclusively with centromeric Tgut716A satellite DNA arrays, with virtually no occurrences outside these repetitive regions. This centromere-specific enrichment mirrors the known distribution pattern of CENP-B boxes in primate genomes, where they are predominantly found within alpha-satellite DNA at centromeres (Suntronpong et al. 2016). This spatial restriction, combined with the high sequence conservation of zebra finch TIGD4 boxes (Figure X), strongly supports the hypothesis that TIGD4 serves the centromeric function in Zebra finch that CENP-B later acquired in primates. The emergence of CENP-B in the primate lineage thus appears to represent a molecular replacement event, where a newly evolved pogo/Tigger family member assumed the centromeric targeting role from its ancient paralog, subsequently leading to relaxed selection and functional decay of TIGD4 in primates while maintaining its essential centromeric function in other vertebrate lineages.

*References*


1. Formenti, G., Abueg, L., Brajuka, A., Brajuka, N., Gallardo-Alba, C., Giani, A., Fedrigo, O., and Jarvis, E.D. (2022). Gfastats: conversion, evaluation and manipulation of genome sequences using assembly graphs. Bioinformatics *38*, 4214–4216.

2. Formenti, G., Koo, B., Sollitto, M., Balacco, J., Brajuka, N., Burhans, R., Duarte, E., Giani, A.M., McCaffrey, K., Medico, J.A., et al. (2025). Evaluation of sequencing reads at scale using rdeval. bioRxivorg. https://doi.org/10.1101/2025.02.01.636073.

3. Rautiainen, M., and Marschall, T. (2020). GraphAligner: rapid and versatile sequence-to-graph alignment. Genome Biol. *21*, 253.

4. Rautiainen, M., Nurk, S., Walenz, B.P., Logsdon, G.A., Porubsky, D., Rhie, A., Eichler, E.E., Phillippy, A.M., and Koren, S. (2023). Telomere-to-telomere assembly of diploid chromosomes with Verkko. Nat. Biotechnol. *41*, 1474–1482.

5. Atema, E., Mulder, E., Dugdale, H.L., Briga, M., van Noordwijk, A.J., and Verhulst, S. (2015). Heritability of telomere length in the Zebra Finch. J. Ornithol. *156*, 1113–1123.

6. Navarro, F.C.P., and Galante, P.A.F. (2015). A genome-wide landscape of retrocopies in primate genomes. Genome Biol. Evol. *7*, 2265–2275.

7. Conceição, H.B., Mercuri, R.L.V., de Castro, M.P.M., Ohara, D.T., Guardia, G.D.A., and Galante, P.A.F. (2024). RCPedia: a global resource for studying and exploring retrocopies in diverse species. Bioinformatics *40*. https://doi.org/10.1093/bioinformatics/btae530.

8. Carelli, F.N., Hayakawa, T., Go, Y., Imai, H., Warnefors, M., and Kaessmann, H. (2016). The life history of retrocopies illuminates the evolution of new mammalian genes. Genome Res. *26*, 301–314.

9. Umu, S.U., Paynter, V.M., Trondsen, H., Buschmann, T., Rounge, T.B., Peterson, K.J.,

and Fromm, B. (2023). Accurate microRNA annotation of animal genomes using trained covariance models of curated microRNA complements in MirMachine. Cell Genom. *3*, 100348.

10. Lin, Y.-C., Balakrishnan, C.N., and Clayton, D.F. (2014). Functional genomic analysis and neuroanatomical localization of miR-2954, a song-responsive sex-linked microRNA in the zebra finch. Front. Neurosci. *8*, 409.

11. Gao, B., Wang, Y., Diaby, M., Zong, W., Shen, D., Wang, S., Chen, C., Wang, X., and Song, C. (2020). Evolution of pogo, a separate superfamily of IS630-Tc1-mariner transposons, revealing recurrent domestication events in vertebrates. Mob. DNA *11*, 25.

12. Casola, C., Hucks, D., and Feschotte, C. (2008). Convergent domestication of pogo-like transposases into centromere-binding proteins in fission yeast and mammals. Mol. Biol. Evol. *25*, 29–41.

13. Lipshutz, S.E., Howell, C.R., Buechlein, A.M., Rusch, D.B., Rosvall, K.A., and Derryberry, E.P. (2022). How thermal challenges change gene regulation in the songbird brain and gonad: Implications for sexual selection in our changing world. Mol. Ecol. *31*, 3613–3626.
