## Supplementary Figures for "The complete genome of a songbird"

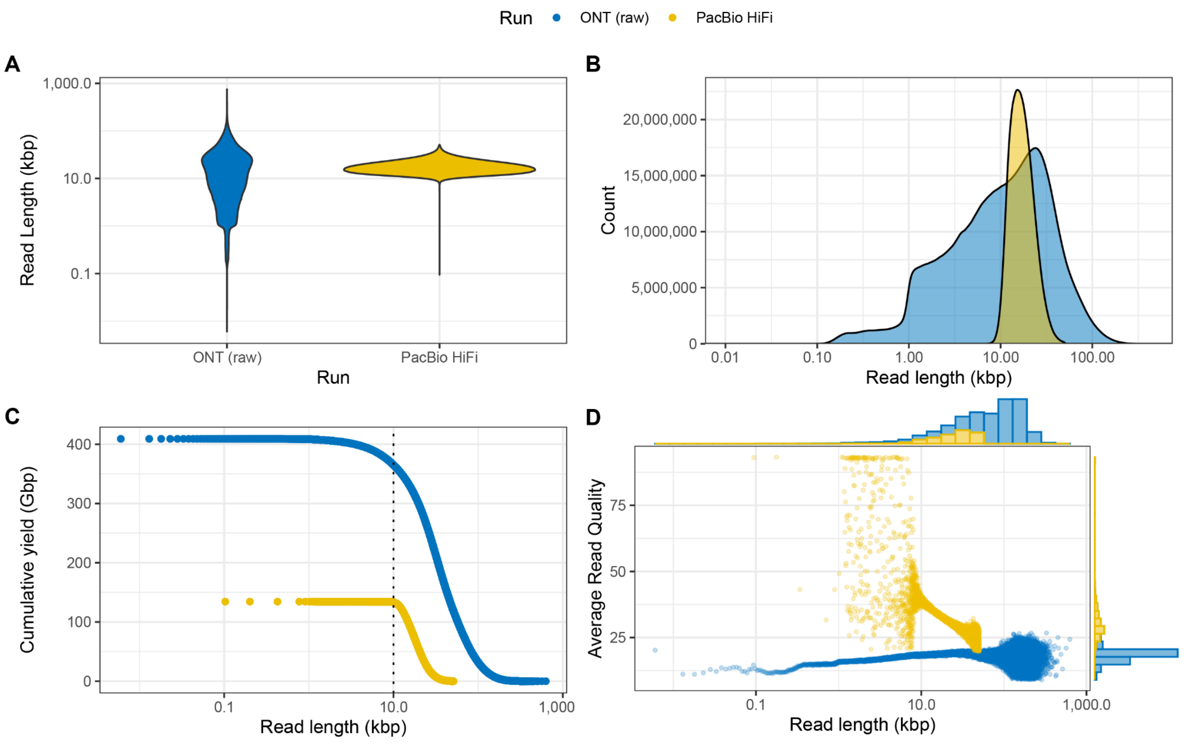


**Supplementary Figure 1. Representative QC report plots from rdeval comparing raw ONT and PacBio HiFi reads for bTaeGut7.** (A) Read length violin plot (log-scaled). ONT reads have a wider range of read lengths (6-751,592 bp) compared to PacBio HiFi (94-51,361 bp). (B) Read length density plots (log-scaled). Similar to the violin plots, the density plots show that ONT reads have a wider range of lengths. (C) Read length inverse cumulative distributions. The distribution can be used to assess read coverage at certain read length cutoffs. For ONT, there is about 350 Gbp of coverage for reads 10 kbp or longer (see vertical line), and for HiFi it is about 140 Gbp of coverage for reads 10 kbp or longer. (D) Read length vs. Average read quality, plotted with marginal histograms. As expected, HiFi read quality is overall higher than ONT. In addition, for HiFi reads above 10kbp, read length correlates inversely with average read quality at varying magnitudes, while for ONT reads, read lengths between 0.1 and 10 kbp positively correlate with average read quality.

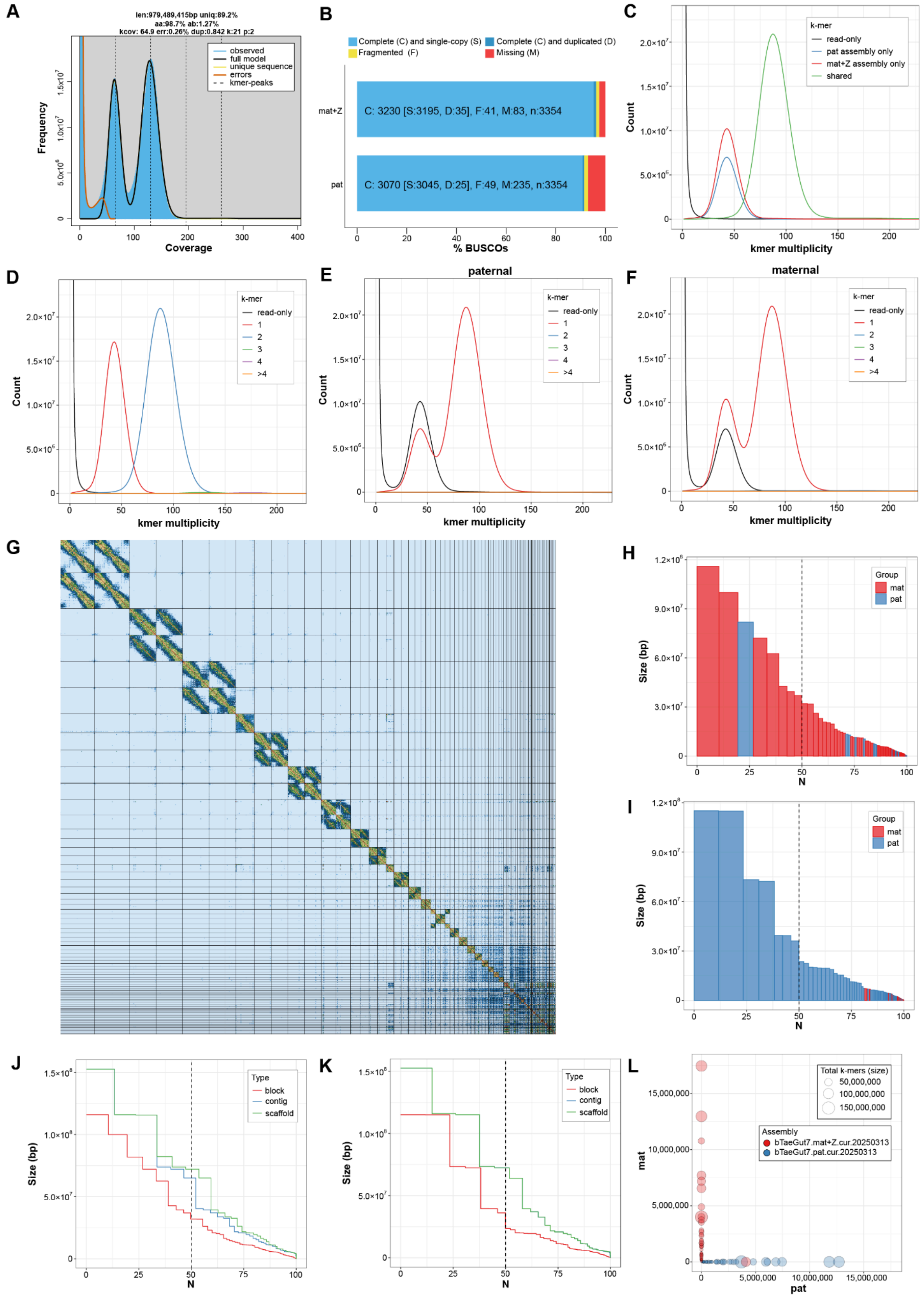


**Supplementary Figure 2. Assembly QC.** (A) GenomeScope2.0 analyses output for 21-mers computed using Meryl from bTaeGut7 HiFi reads. GenomeScope2[1] estimates genome haploid length estimated at 979,489,415bp, with estimated k-coverage of 129.8x and maximum read error rate 0.26%. (B) BUSCO analysis to determine representation of near-universal single-copy orthologs in the vertebrate lineage. Results show high completeness of gene content, 96.3% in the maternal haplotype + Z and 91.5% in the paternal haplotype, which is likely an underestimate due to the removed Z chromosome. (C) Spectra assembly plot, showing that the majority of 21-mers are shared between the maternal and paternal assemblies. The maternal + Z assembly has more unique 21-mers than the paternal assembly, due to the addition of the Z chromosome. (D) Spectra copy number plot, showing final k-mer multiplicity of 21-mers found in parental Illumina reads. (E-F) Spectra copy number plots for paternal and maternal assemblies, respectively. (G) Pretext map of dual curated assembly representing combined maternal and paternal haplotypes with gap and telomere tracks. Each square represents a pair of chromosomes, ordered from largest to smallest in size. (H,I) Phase block NG* plots. A phase block is defined by at least two hap-mers from the same haplotype. The x-axis reflects the percentage of the genome size covered by phase blocks of the size represented on the y-axis. Red and blue bars represent maternal and paternal hapmers found in the assembly. The paternal hapmer represented by the blue bar at 25N in the maternal assembly can be attributed to the Z chromosome. Otherwise, mismatched blocks are much smaller. J,K) Contiguity plots of curated scaffolds for maternal and paternal assemblies respectively. Roughly similar levels of contiguity between the two haplotypes are observed. L) Blob plot showing phasing of maternal and paternal hapmers among total k-mers counted per curated scaffolds. Each blob represents an individual scaffold, and the size of each blob represents the total number of k-mers counted for each scaffold. Blobs are positioned by the number of contained maternal and paternal hapmers.

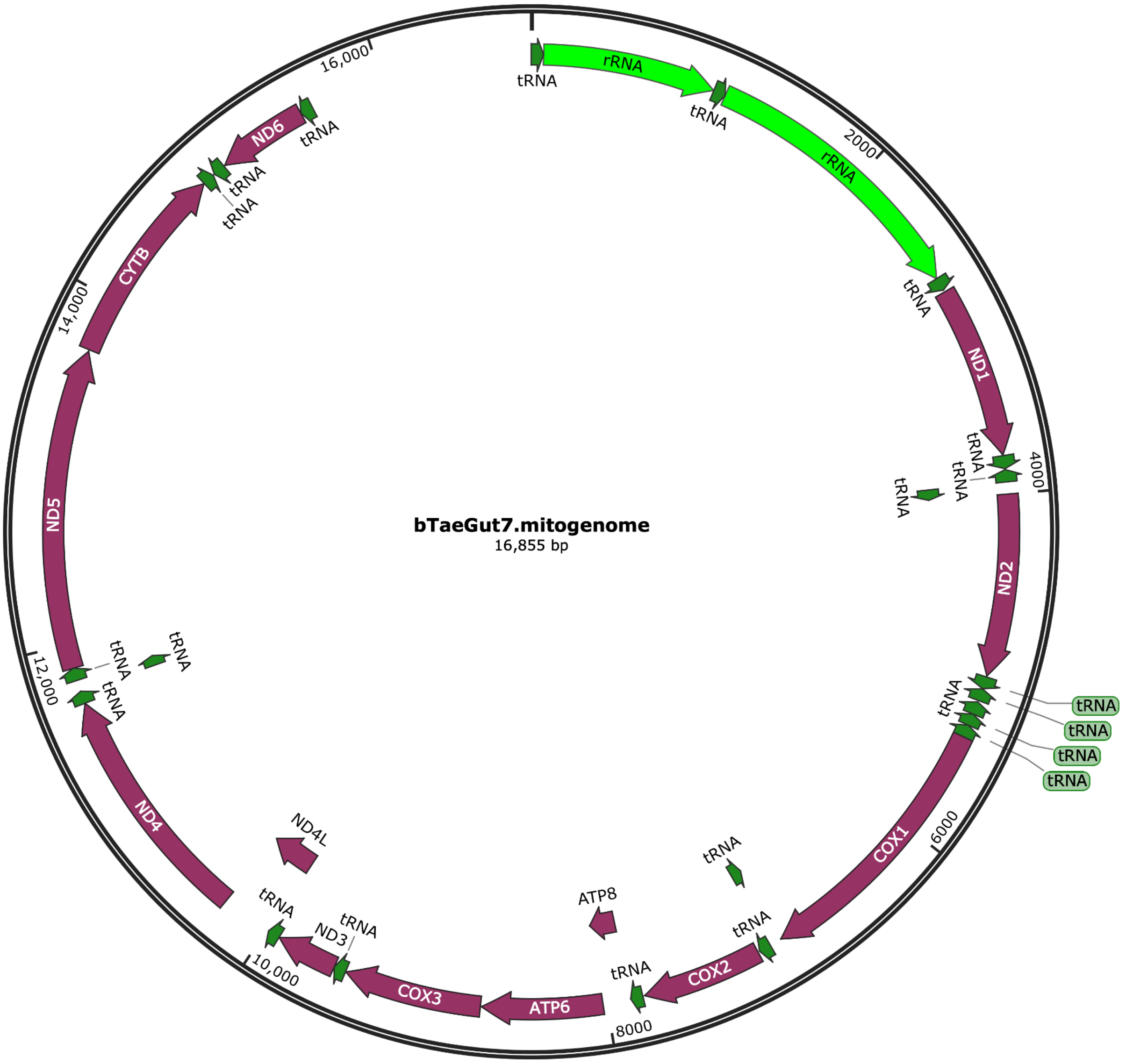


**Supplementary Figure 3. Complete mitogenome assembly for bTaeGut7 (16,855 bp)**. It shows the canonical 37 vertebrate genes, 13 encoding for proteins, 22 for tRNAs, and 2 for rRNAs, plus a control region of 1,275 bp. Mitos2[2] annotation shows no errors or warnings.

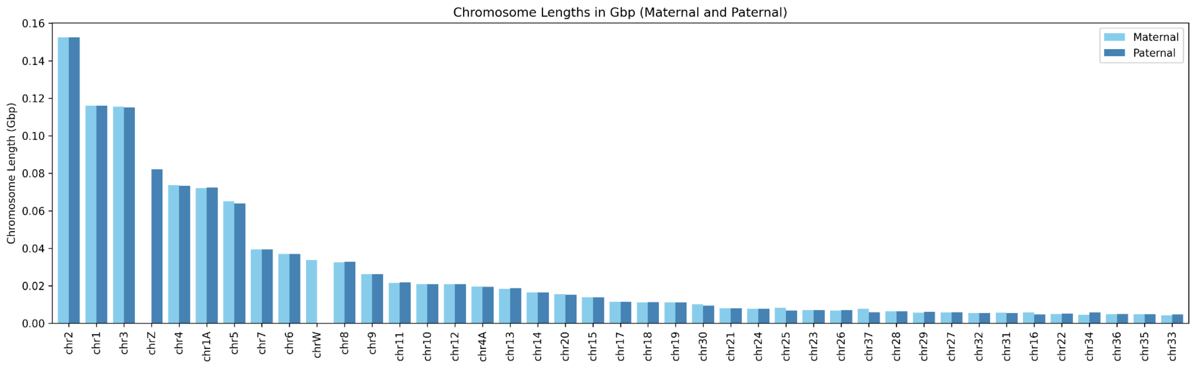


**Supplementary Figure 4. Maternal and paternal chromosomes sorted by length.** Macrochromosomes were originally named after chicken, in which derived fusions between chr1 and chr1A, chr4 and chr4A are observed. The other chromosomes were assigned by size but do not reflect their real length due to gaps in previous assemblies.

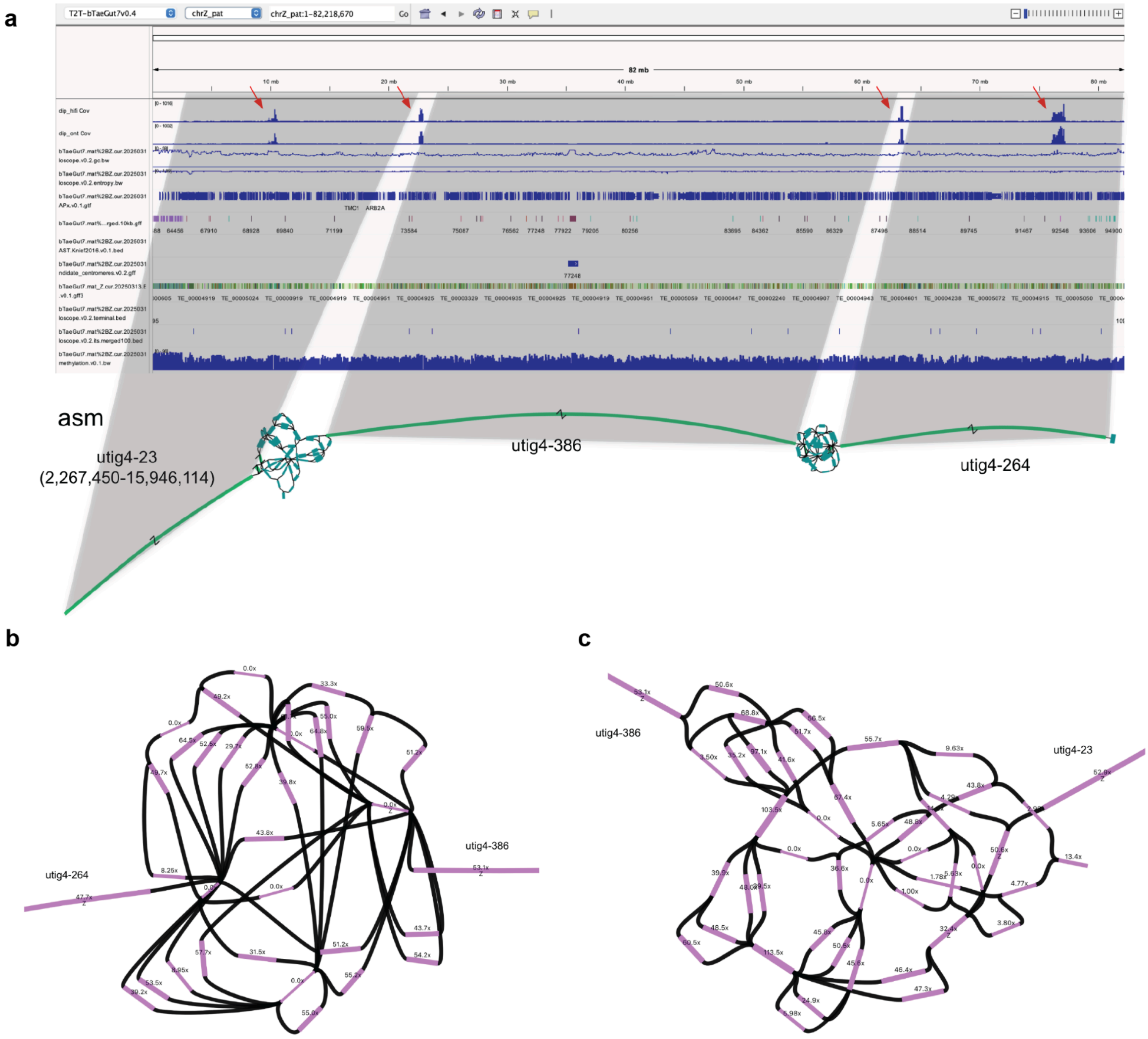


**Supplementary Figure 5. Assembly tangles and gaps in chrZ pre-curation**. (A) IGV screenshot with the location of the 4 gaps in chrZ highlighted with red arrows. In the HiFi-only assembly (asm), only two tangles are present, but the left arm of the chromosome is missing. (B,C) highlight of the two tangles, with HiFi coverage values reported for each node.

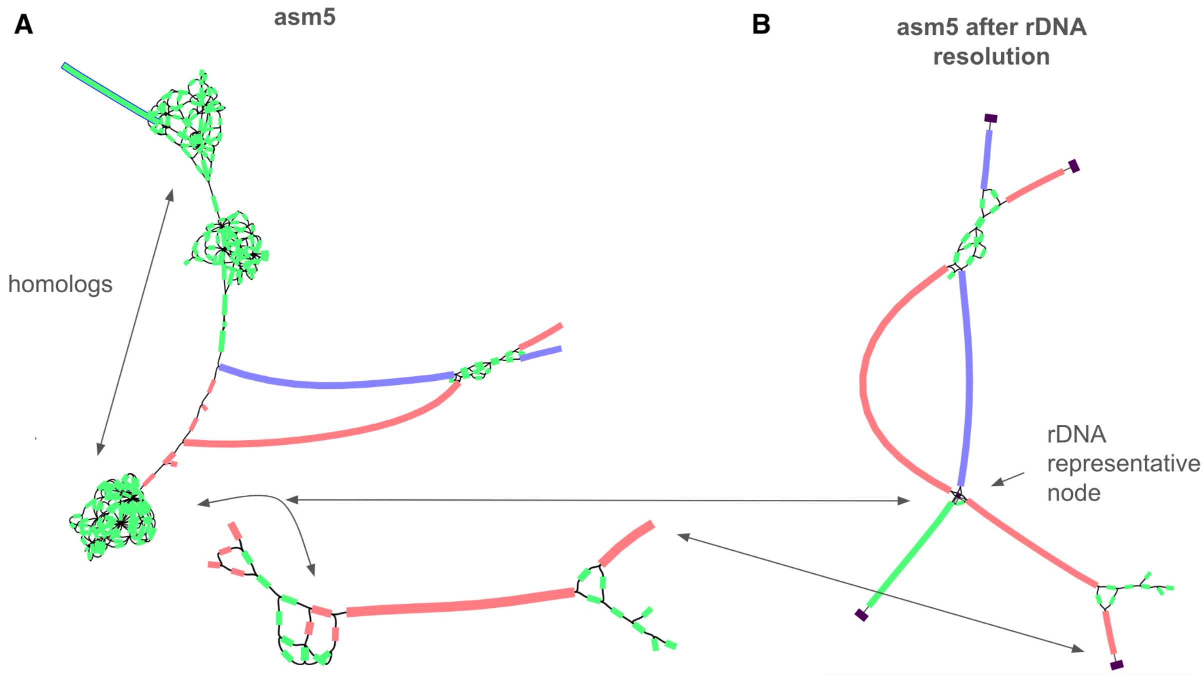


**Supplementary Figure 6. rDNA tangles in the assembly graph pre-curation**. 18S rDNA repetitive array generally is too similar and repetitive to be resolved with current technology (GC content >70%). In zebra finch it is found in one chromosome, chr37. Chr37 was disconnected in the HiFi-only assembly (asm1), and partially disconnected (the maternal haplotype) in the assembly that added corrected ONT reads (asm5), with 3 large tangles detected (A). The two haplotypes show full separation, suggesting sequence divergence in the rDNA sequence. Asm5 gets reconnected after Verkko's[3] rDNA resolution (B). A novel strategy was implemented to provide a sequence model for the rDNA using the output of Ribotin[4].

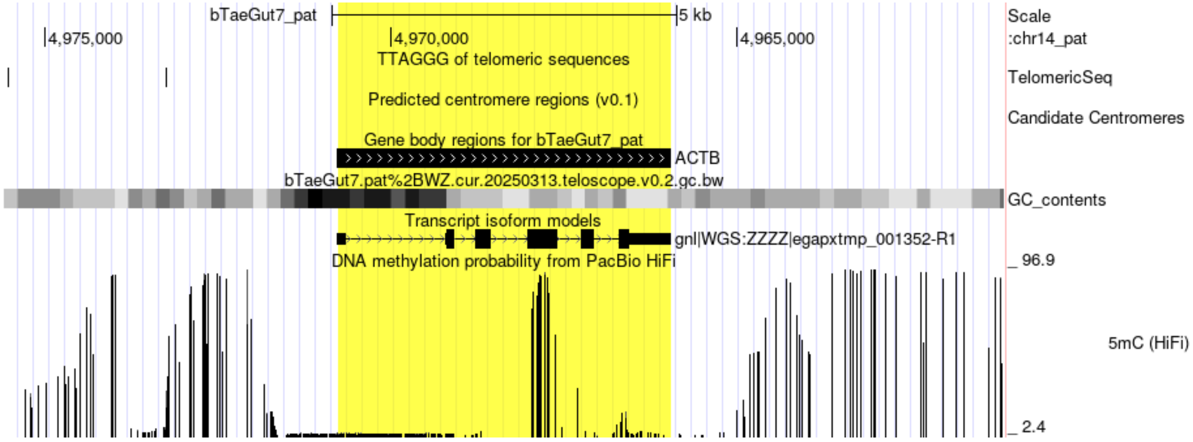


**Supplementary Figure 7. PacBio HiFi read-based DNA methylation probability near the housekeeping gene *ACTB* on the bTaeGut7 paternal assembly.** As visualized in the UCSC Genome Browser, each row displays, in order, the canonical telomeric repeat sequences ((TTAGGG)n), predicted centromere regions, GC content, gene annotations, transcript isoforms, and DNA methylation probabilities calculated from PacBio HiFi reads derived from blood samples of the bTaeGut7 individual. The figure was generated from https://genome.ucsc.edu/s/clee03/bTaeGut7_pat_ACTB.

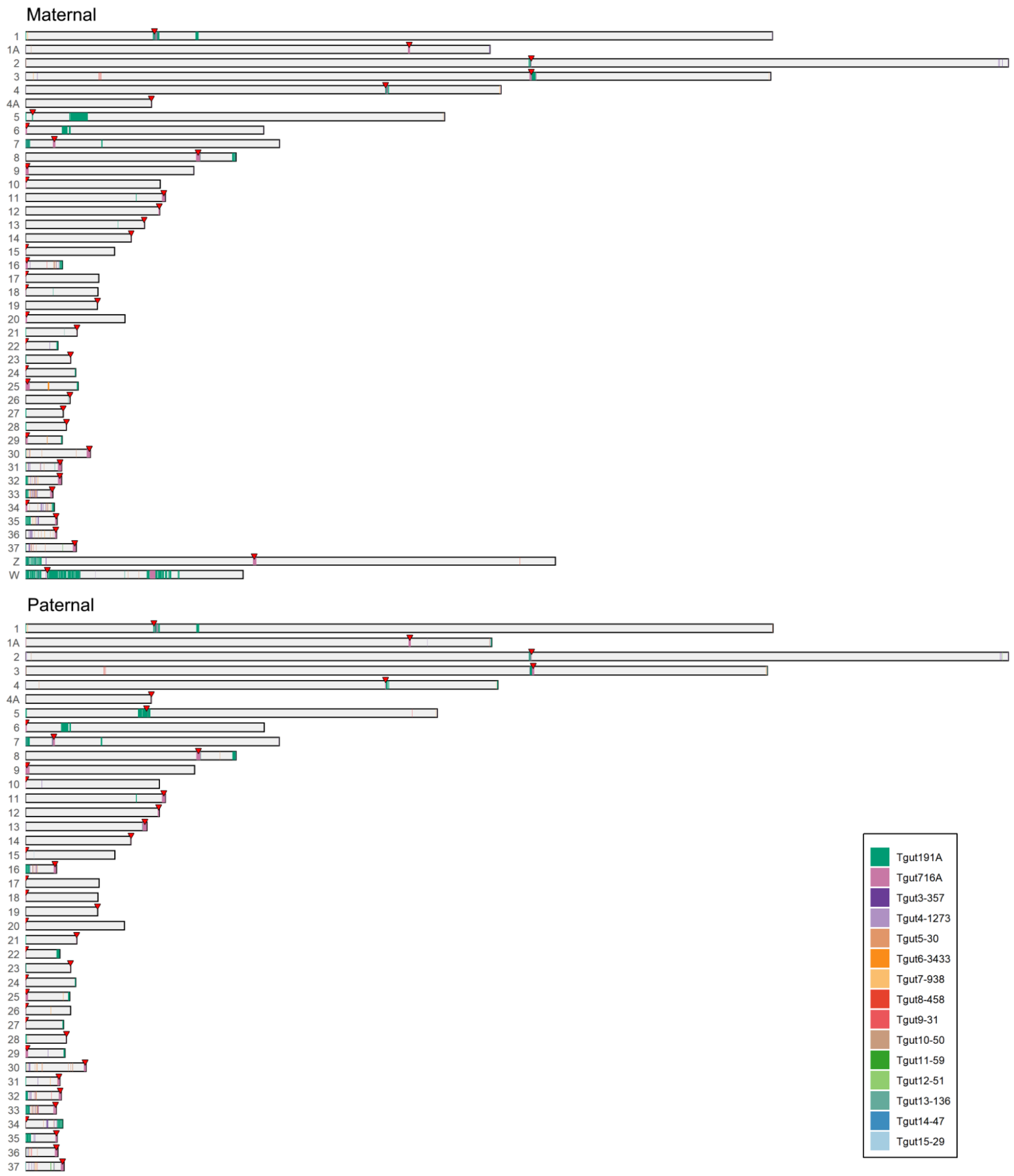


**Supplementary Figure 8. Satellite DNA distribution across bTaeGut7 chromosomes**. Chromosomes are grouped by haplotype, with both Z and W chromosomes in the maternal haplotype as represented in the primary assembly. Centromere positions are represented with red tick marks. Tandem repeats are colored.

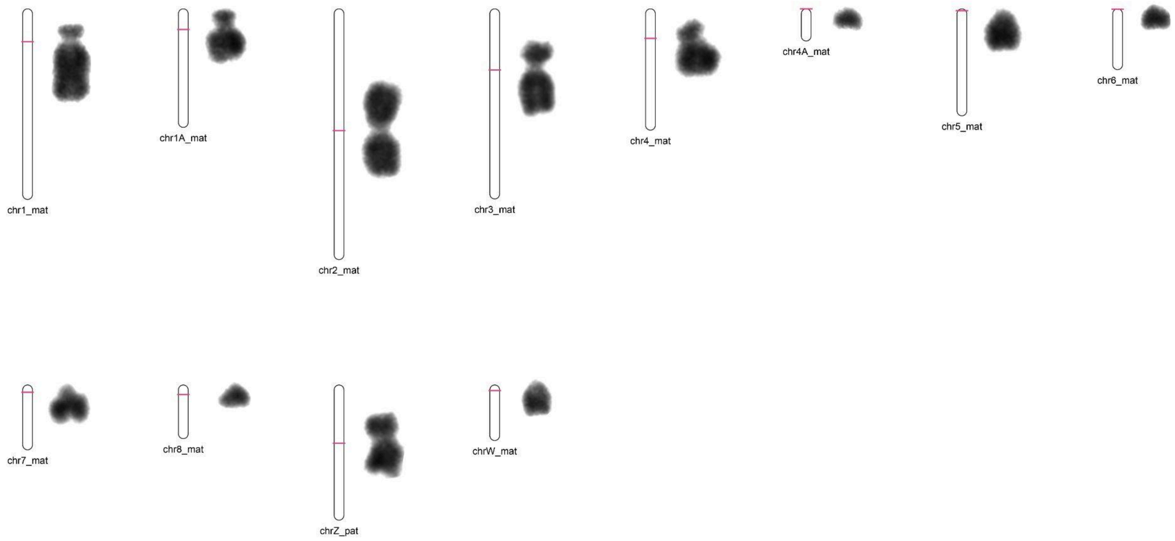


**Supplementary Figure 9. Centromere location inferred from the assembly versus chromosome primary constriction**. The drawings to the left of the cytogenetic images are schematic representations of chromosomes from the primary assembly shown to scale, with centromere annotations aligned to the corresponding karyotype images.

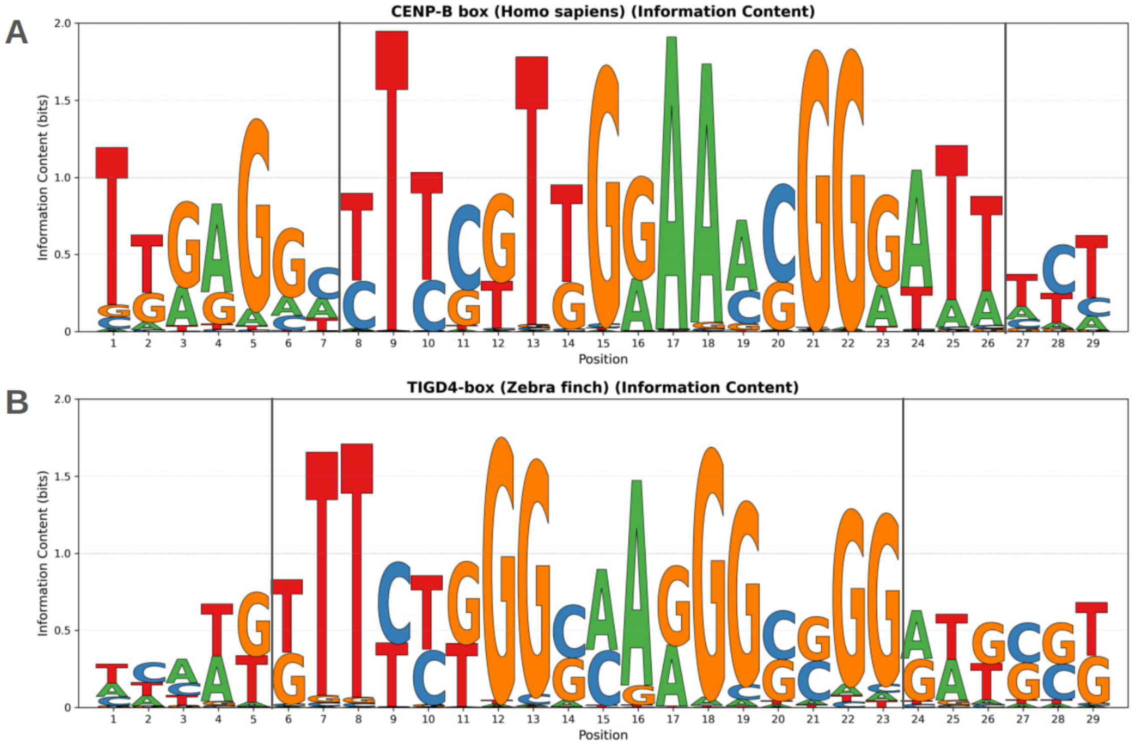


**Supplementary Figure 10. Sequence logos of CENP-B and TIGD4 boxes**. The canonical human CENP-B box was used as a reference motif. To assess its conservation, we searched for all genomic occurrences within edit distance ≤5 for cross-species and ≤2 for the same species in the human HG002 diploid (A) and zebra finch T2T assemblies (B). To establish motif boundaries and provide negative controls, we added ±5 flanking positions. This revealed a sharp transition from the conserved core box to the surrounding random sequence, indicating clear enrichment at the box itself. Sequence borders are shown with vertical lines.

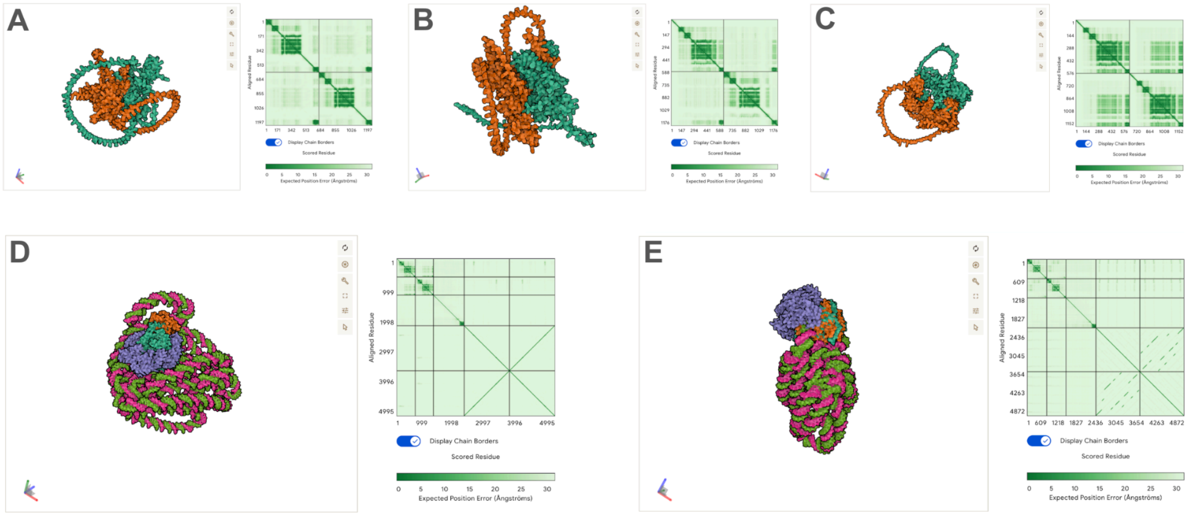


**Supplementary Figure 11. AlphaFold-predicted dimeric structures and interaction interfaces of CENP-B and TIGD4 proteins**. (A-C) Structural models with corresponding PAE matrices for (A) human CENP-B homodimer showing C-terminal dimerization (residues 539-598), (B) zebra finch TIGD4 homodimer with similar C-terminal dimerization (residues ~540-576), and (C) CENP-B/TIGD4 heterodimer (ipTM = 0.25, pTM = 0.31) demonstrating potential inter-protein interactions. PAE matrices indicate prediction confidence, with darker green regions representing higher confidence contacts. (D-E) Structural models of centromeric protein-DNA complexes. (D) Zebra finch TIGD4 homodimer assembled with CENP-C and Tgut716A centromeric DNA. (E) Human CENP-B homodimer with CENP-C and alphoid satellite DNA.

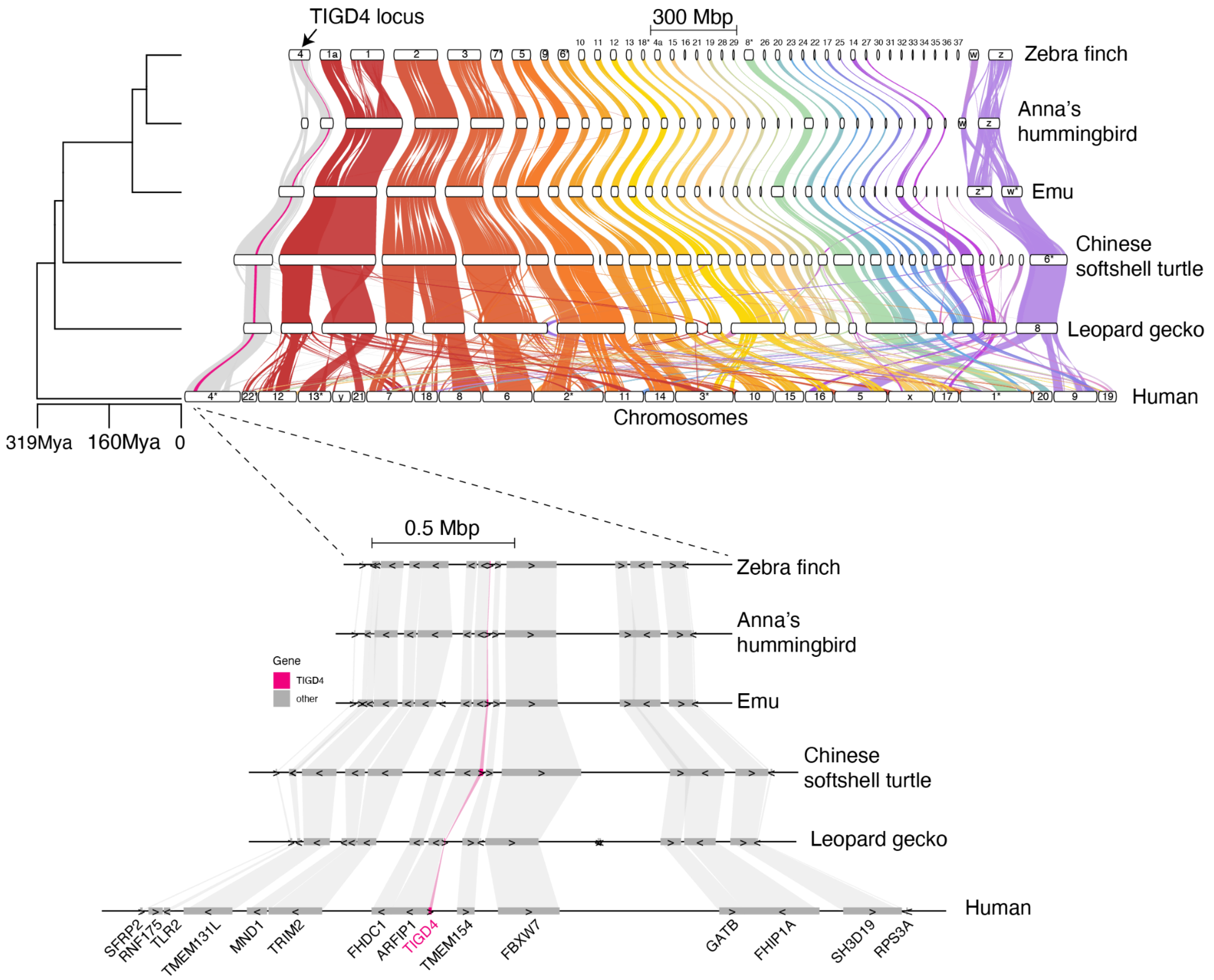


**Supplementary Figure 12.** Synteny conservation for TIGD4. Global genome synteny from zebra finch through to human. Phylogenetic relationships and estimated divergence dates are shown on the left. The TIGD4 locus (arrowed) is on zebra finch chromosome 4 and tracked through in pink. The expanded view of the TIGD4 locus shows that gene order surrounding TIGD4 (highlighted in pink) has remained conserved in all species depicted here, so it likely represents the ancestral amniote state.

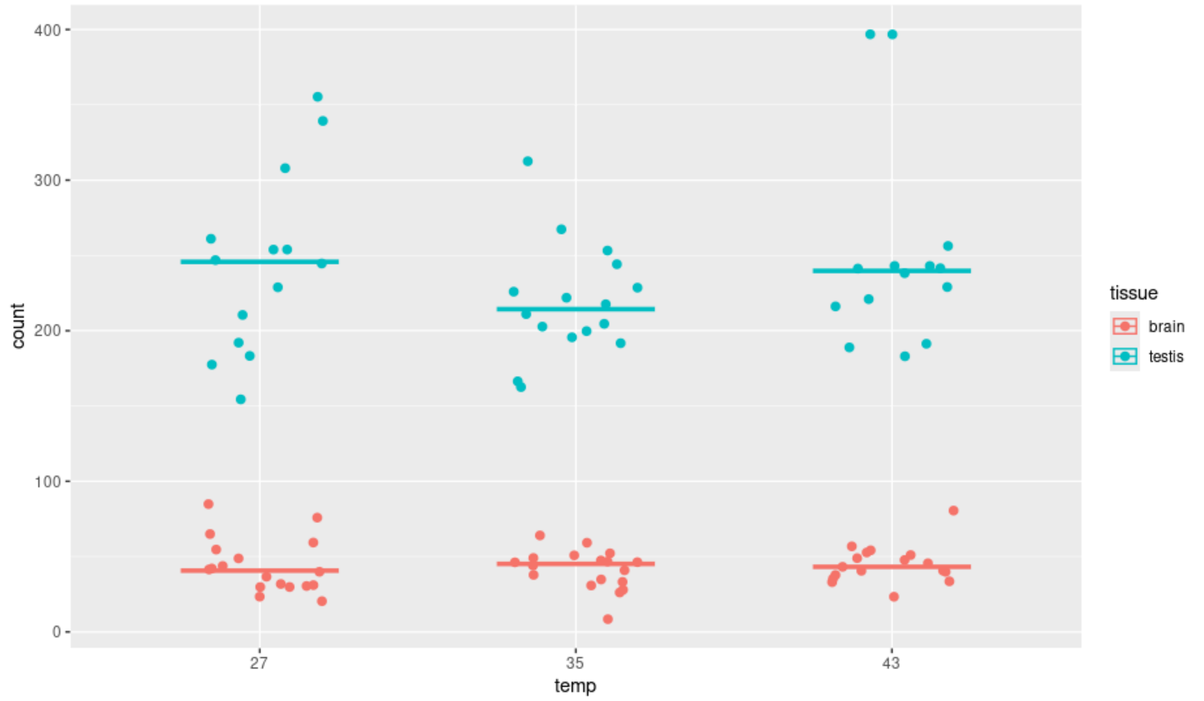


**Supplementary Figure 13. Overexpression of TIGD4 in testis versus brain**. Testis has higher expression relative to the brain, at all temperatures. Temperature does not affect expression either in the testis or brain.

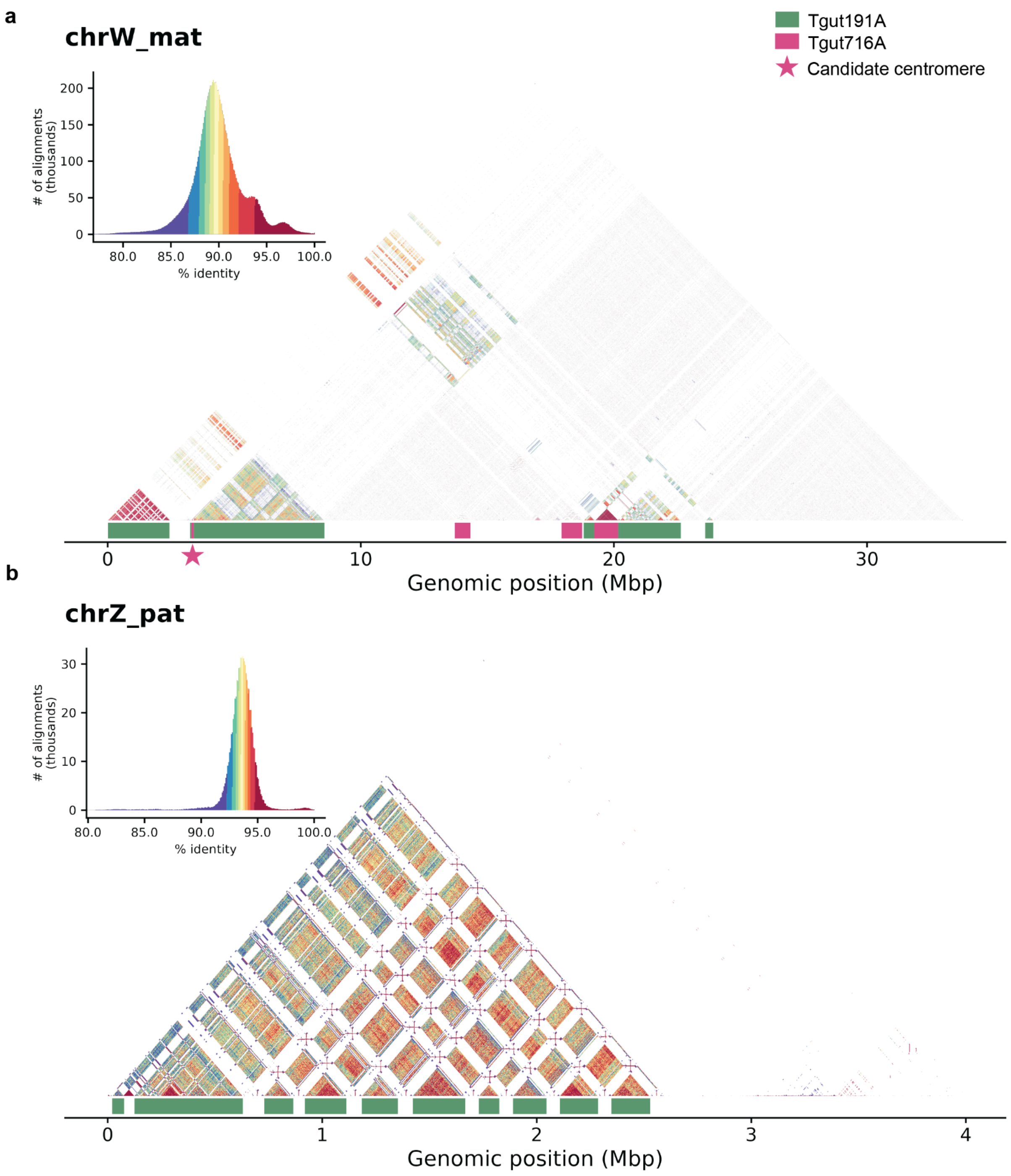


**Supplementary Figure 14.** Annotated StainedGlass[5] plots for chrZ and chrW. (A) In chrW, the PAR region is ~3.3 Mbp, and it is directly adjacent to the putative Tgut716A centromere array (star). The position of the array matches previous cytogenetic observations[6]. Other Tgut716A arrays exist in chrW, but the one next to the PAR is the only one hypomethylated and is in agreement with the position of the kinetochore binding site from previous cytogenetic observations, making it the most likely candidate centromere. There exist 2 other large Tgut191A arrays, but they show significantly lower repeat unity intrasequence similarity, suggesting that they are not functional anymore. (B) The PAR region of chrZ, showing high sequence similarity with chrW is in the first ~3.4 Mbp, predominantly constituted by the Tgut191A satellite (green). No other large Tgut191A regions are present in chrZ.

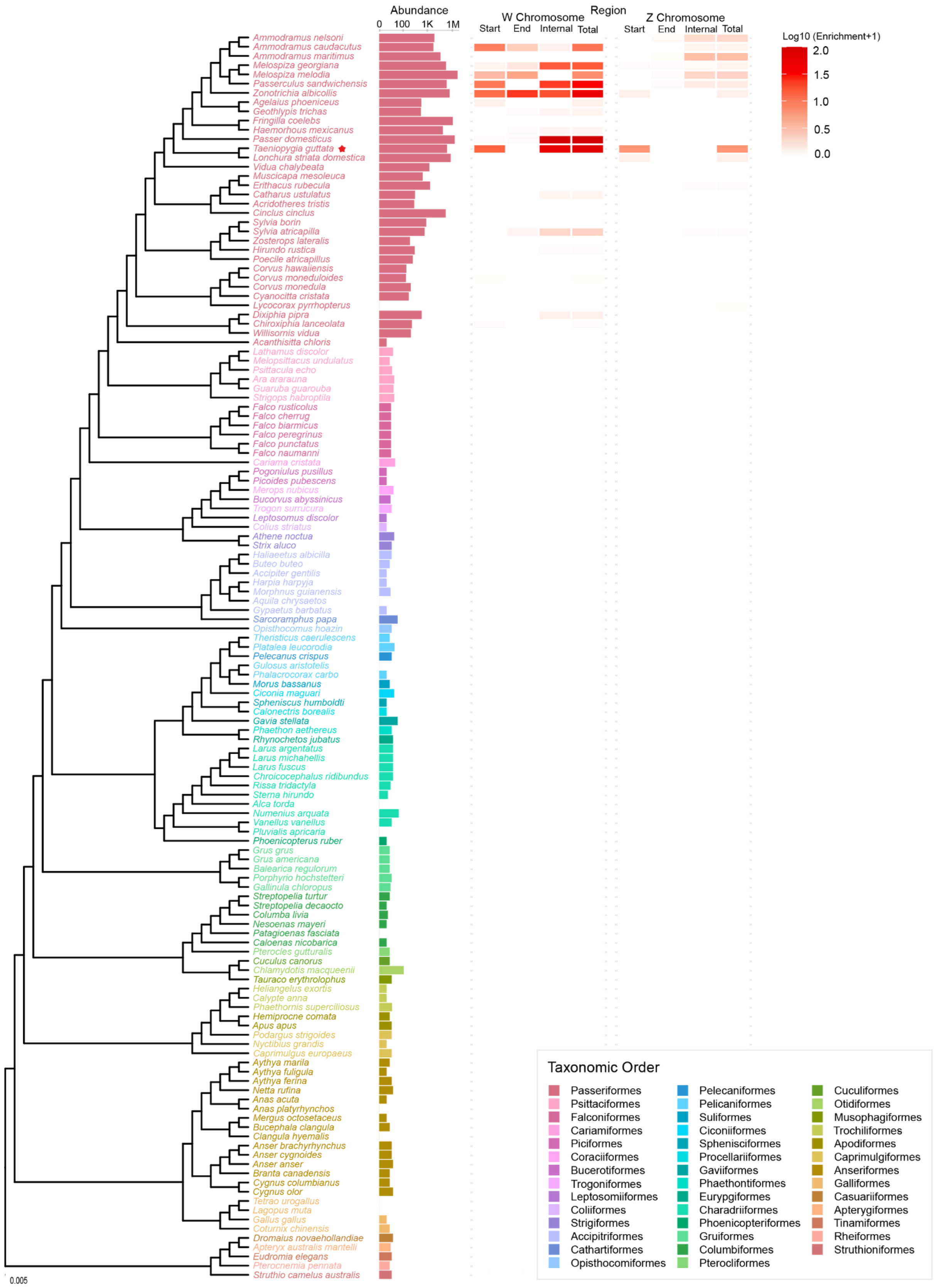


**Supplementary Figure 15.** Phylogenetic distribution of repeat abundance across the genome and sex chromosomes, and assembly contiguity, in VGP bird assemblies. The phylogeny is derived from the Open Tree of Life[7], with tip colors indicating avian orders. For each species, the horizontal bar plot (left) shows the genome-wide abundance of

Tgut191A-like repeat hits (log10 scale). To the right, heatmaps display the log10(enrichment+1) of Tgut191A-like repeats along the W and Z chromosomes within three regions: the start (first 3 Mbp window, approximately the size of the PAR region in zebra finch), the end (last 3 Mbp window), and the internal region (sequence between the start and end window). The legend includes only the avian orders present in the tree.

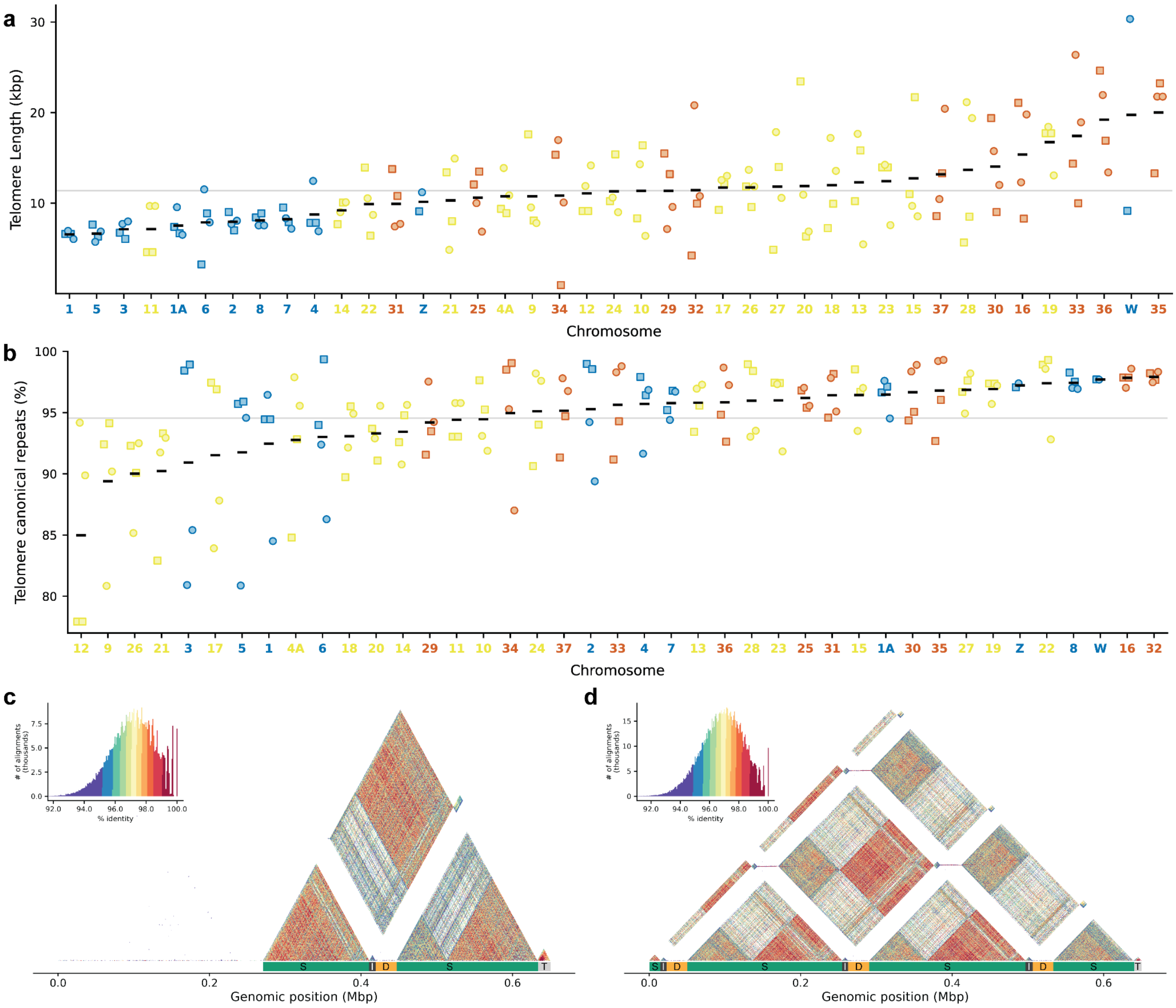


**Supplementary Figure 16.** Exploration of terminal and interstitial bTaeGut7 telomeres. (A) Telomere length distribution across bTaeGut7 chromosomes, ordered by ascending mean. (B) Percentage of telomeres covered by canonical CCCTAA/TTAGGG repeats. For (A) and (B), chromosomes are labeled by TCHEST class (acro, micro, and dot chromosomes, respectively). P- and q-arm telomeres are plotted as squares and circles, and the gray line indicates the assembly mean. c) Interstitial telomeres of maternal chr34 with one ITS unit. d). Interstitial telomeres of paternal chr34 with three ITS units.

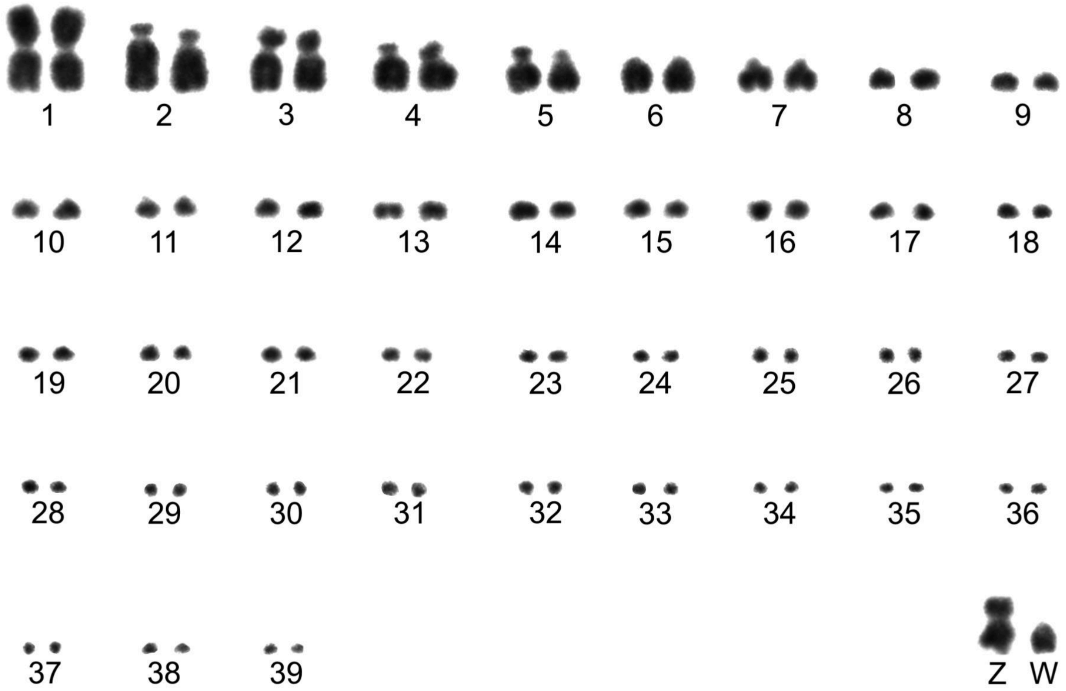


**Supplementary Figure 17.** Conventionally stained representative karyotype of a female *Taeniopygia guttata*. The karyotype was assembled and numbered according to the chromosome size and centromere position. Chr1 (chr2 in the assembly) is metacentric, 2 (chr1) and 7 (chr7) are acrocentric, 3 (chr3), 4 and 5 (similar in size and centromere position, either 4 or 1A), as well as Z are submetacentric and all the remaining, including 6 (chr5) and W, appear telocentric. Putative assignments to the assembled chromosomes are based on the relative size and centromere position.

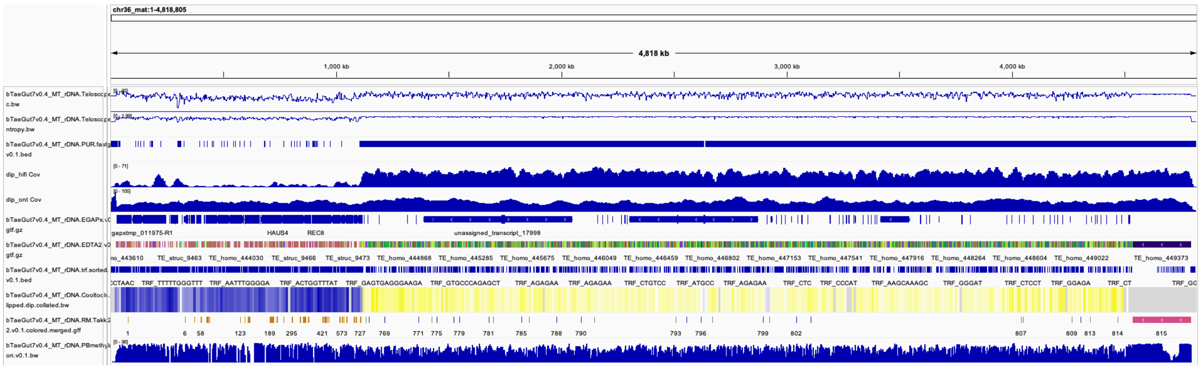

**Supplementary Figure 18.** Chr36 annotation, a representative dot chromosome highlighting the TCHEST structure. (A) GC content. (B) Entropy. (C) unassembled regions. (D) HiFi coverage. (E) ONT coverage. (F) EGAPx gene annotation. (G) Repeat annotation using EDTA2 (features colored according to feature type name). (H) Satellite annotation using TRF[8]. (I) AB compartments. (J) RepeatMasker annotation using library from Takki et al. 2022[9]. (K) PacBio methylation. From right to left, the terminal telomere is followed by the centromere, heterochromatin (B compartment, low gene density), euchromatin (A compartment, minisatellite-dense, gene-rich), and the other telomere. The euchromatic region is GC/GA rich, and PacBio HiFi struggles to sequence it, as shown by the lower coverage.

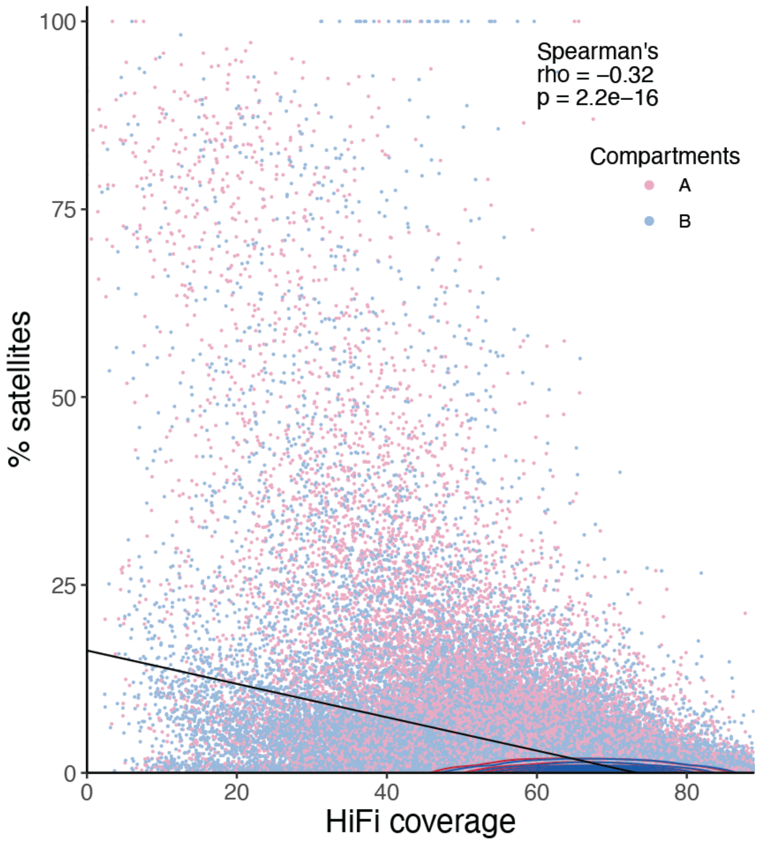


**Supplementary Figure 19. HiFi coverage vs minisatellite content in non-dot chromosomes.**

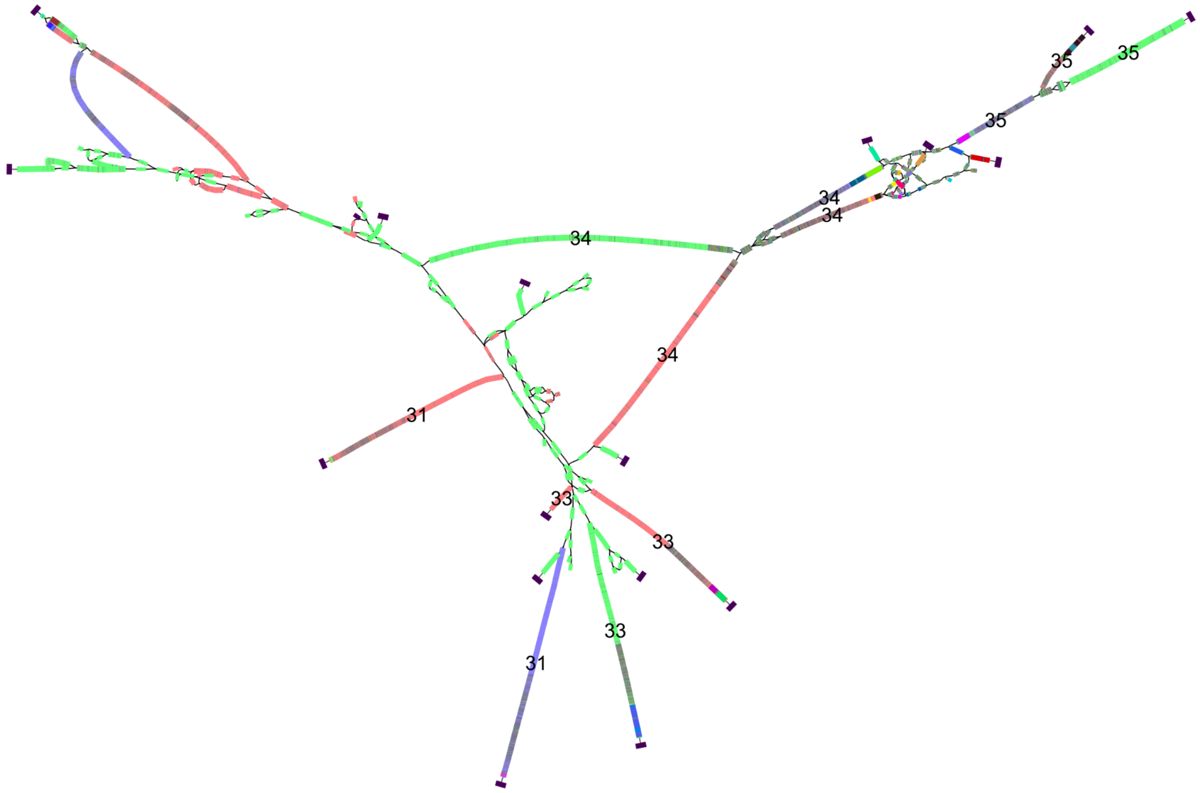


**Supplementary Figure 20. Asm5 assembly graph for dot chromosomes 31, 33-35**. Asm5 used both HiFi and ONT reads for graph construction. Microchromosome 29 is also visible in the upper left quadrant. Blue and red correspond to the paternal and maternal haplotypes, respectively. Green are unitigs that could not be confidently assigned using parental kmers. Satellite annotation generated by TRF[8] is also reported using multiple colors. Euchromatic regions are satellite-rich, and did not appear to have posed an assembly challenge, whereas heterochromatic sequences appear to be more frequently shared, complicating the graph.

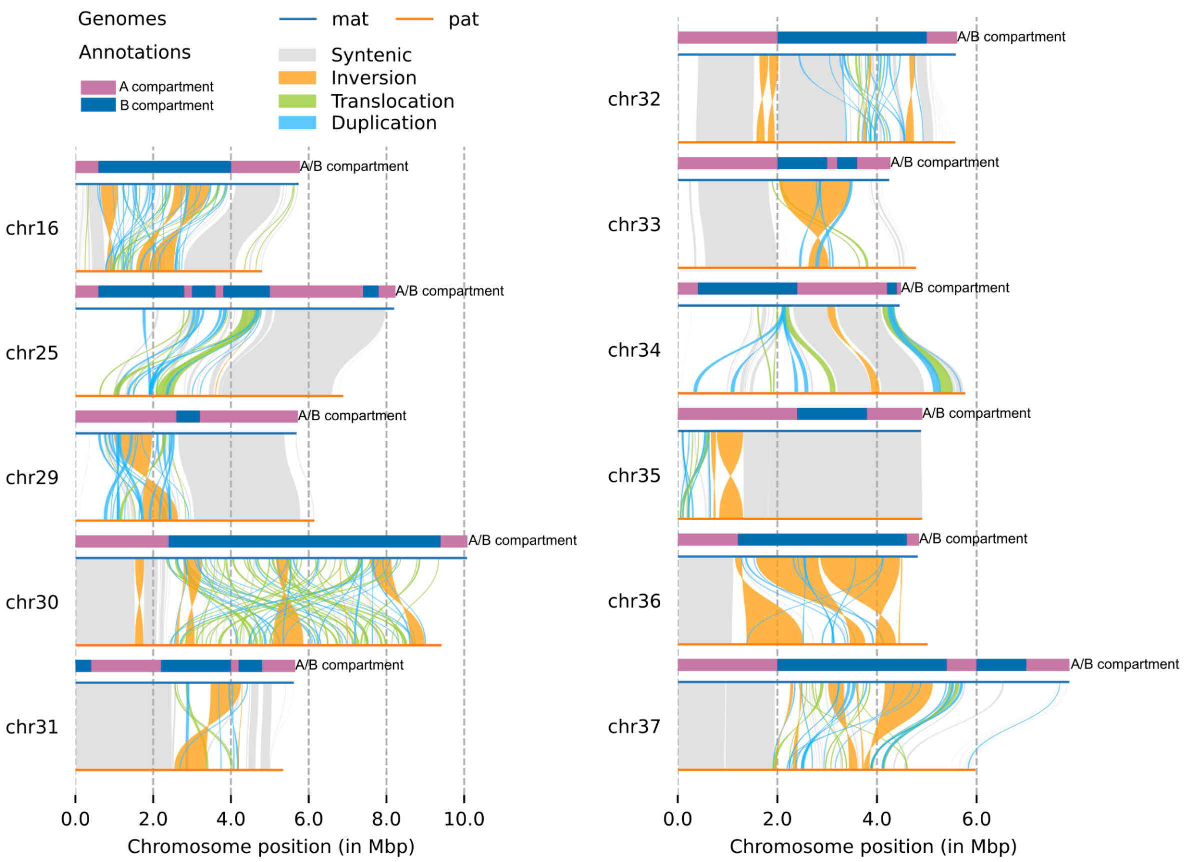


**Supplementary Figure 21. Synteny plot for 11 dot chromosomes with A/B compartment track shown on top of each chromosome.** Most of the genomic rearrangements on dot chromosomes are in heterochromatic regions (B compartment), except for chromosome 29 and 35.

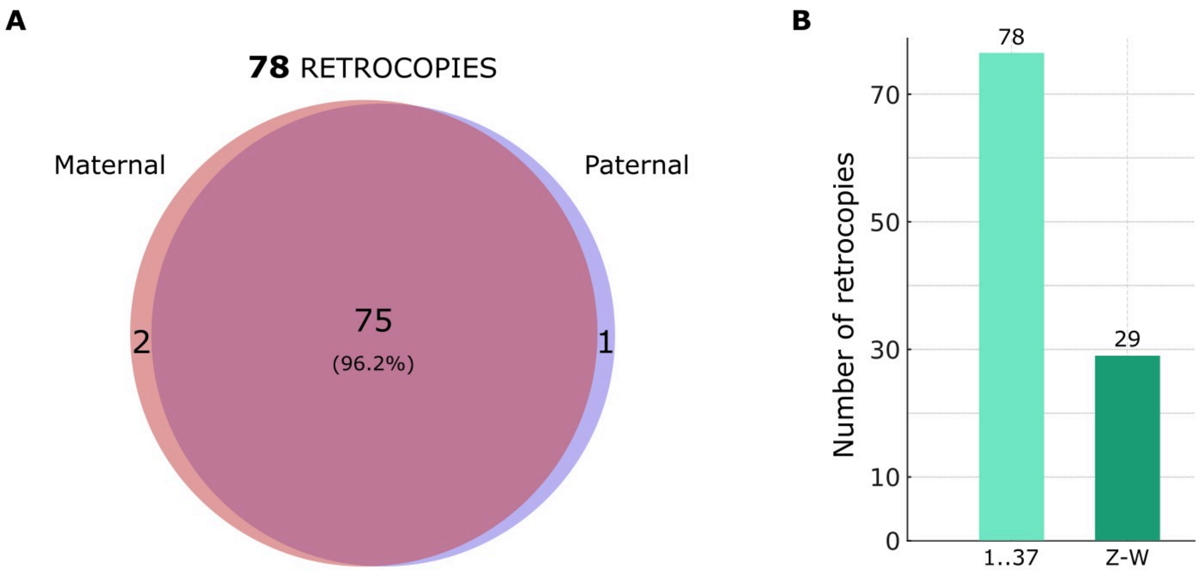


**Supplementary Figure 22. Retrocopies identified in the zebra finch genome assemblies.** (A) Overlap of retrocopies detected independently in the maternal and paternal assemblies. (B) Distribution across autosomes (chromosomes 1–37) and sex chromosomes (Z and W).

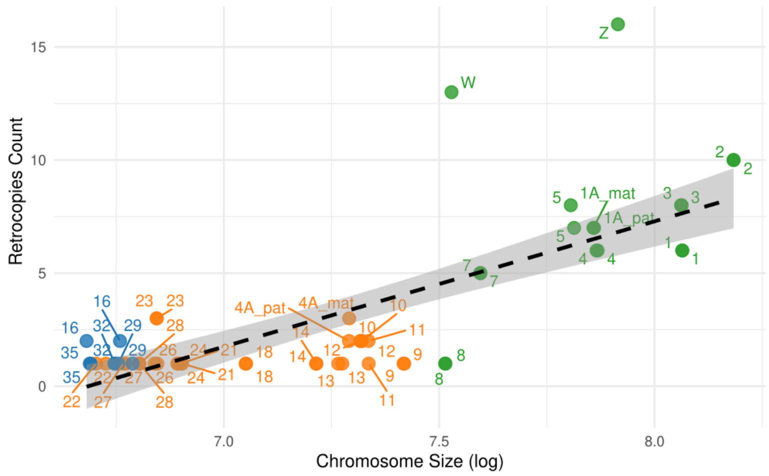


**Supplementary Figure 23. Number of retrocopies per chromosome length.** Counts aggregate maternal and paternal calls. Chromosomes are colored, considering their classification as macrochromosomes (green), microchromosomes (orange), or micro-dot chromosomes (blue). Each point represents a chromosome, with labels indicating chromosome IDs.

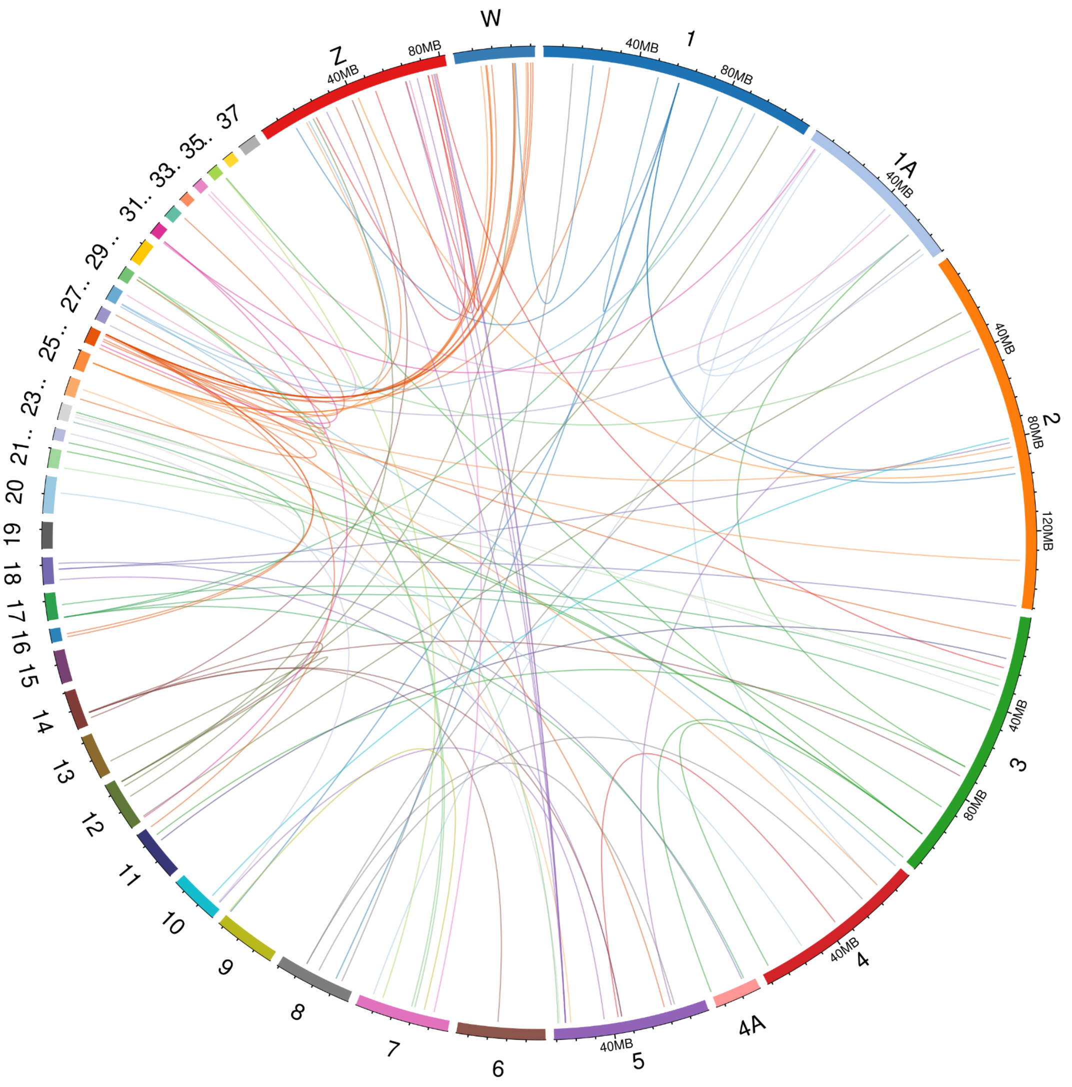


**Supplementary Figure 24. Circos representation of parental gene–retrocopy relationships (maternal assembly).** Chromosomes are displayed as colored segments around the circle. Connecting lines represent parental gene-retrocopy pairs, with line colors matching the chromosome color of the parental gene. Each line connects the parental gene's chromosome to the chromosome containing its retrocopy. For example, a parental gene located on chromosome 12 (orange segment) that has a retrocopy on chromosome Z (red segment) is represented by an orange line connecting these two chromosomes.

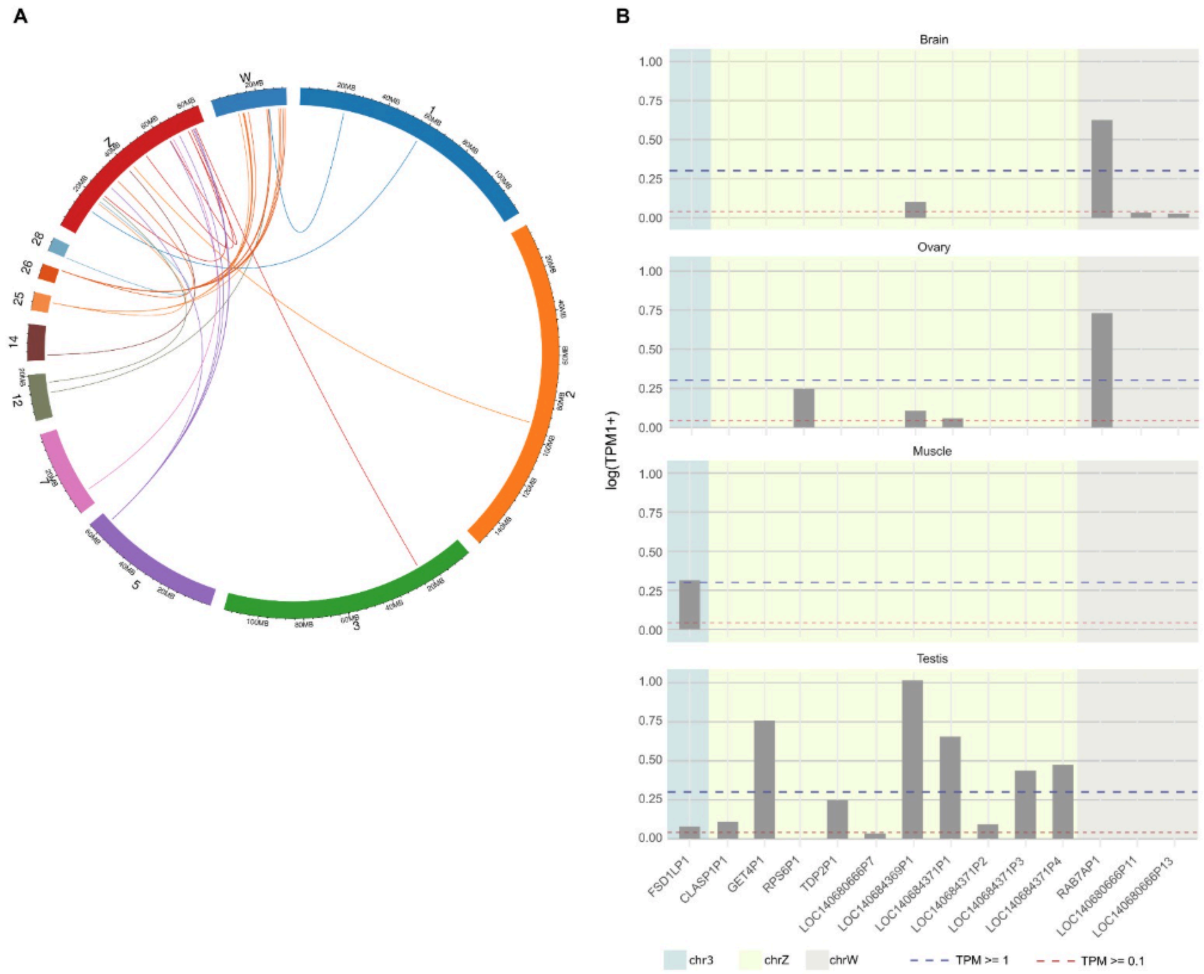


**Supplementary Figure 25. Retrocopy origin and expression patterns on sex chromosomes**. (A) Circos plot showing retrocopies and their parental genes mapped to the sex chromosomes. Colored links connect parental genes to their retrocopies. (B) Expression of sex-chromosome–associated retrocopies. Expression for individual retrocopies across four tissues (Brain, Muscle, Ovary, Testis). Background shading denotes the chromosomal location of each retrocopy (light blue, autosome; light yellow, chrZ; light gray, chrW). Horizontal dashed lines mark expression thresholds (red: TPM ≥ 0.1; blue: TPM ≥ 1).

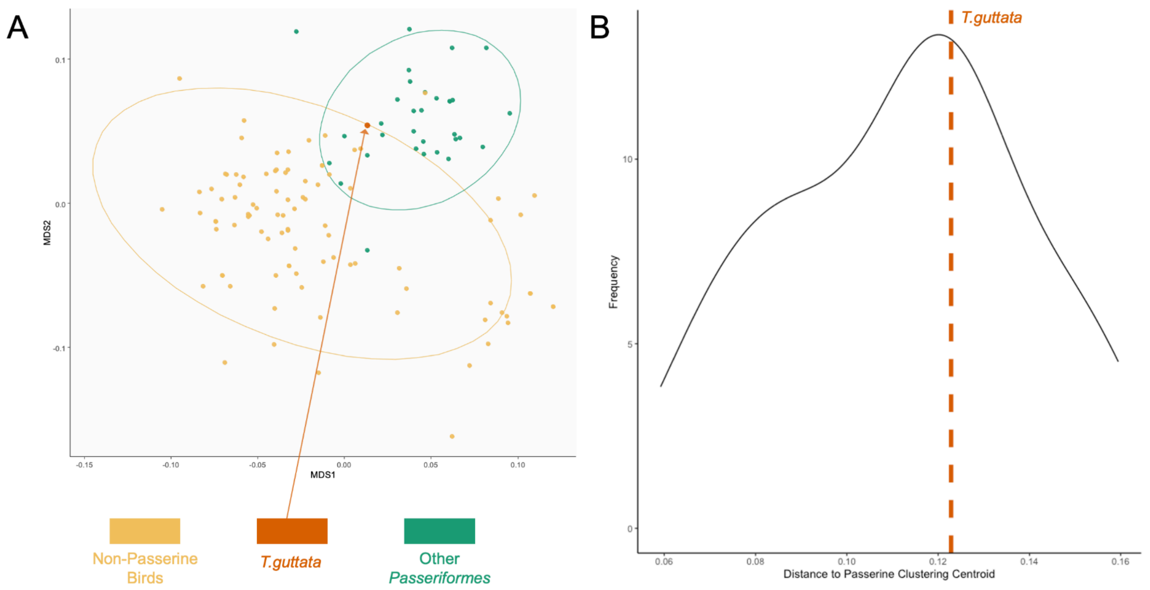


**Supplementary Figure 26. NMDS clustering of Zebra finch T2T in comparison to 131 other bird species.** (A) NMDS clustering of 132 bird species taken from the VGP (only MDS1-2 of 5 shown), with ellipses coloured by group (blue = Passeriformes, yellow = other bird species). The data point for zebra finch is highlighted in orange within the ellipse encompassing all passerine species. (B) Density plot showing NMDS distance of all passerine species from the centroid of the passerine cluster *(i.e.* the distance from a hypothetical 'typical' passerine miRNA repertoire), with an orange dashed line showing the position of the zebra finch close to the centre of the distribution.

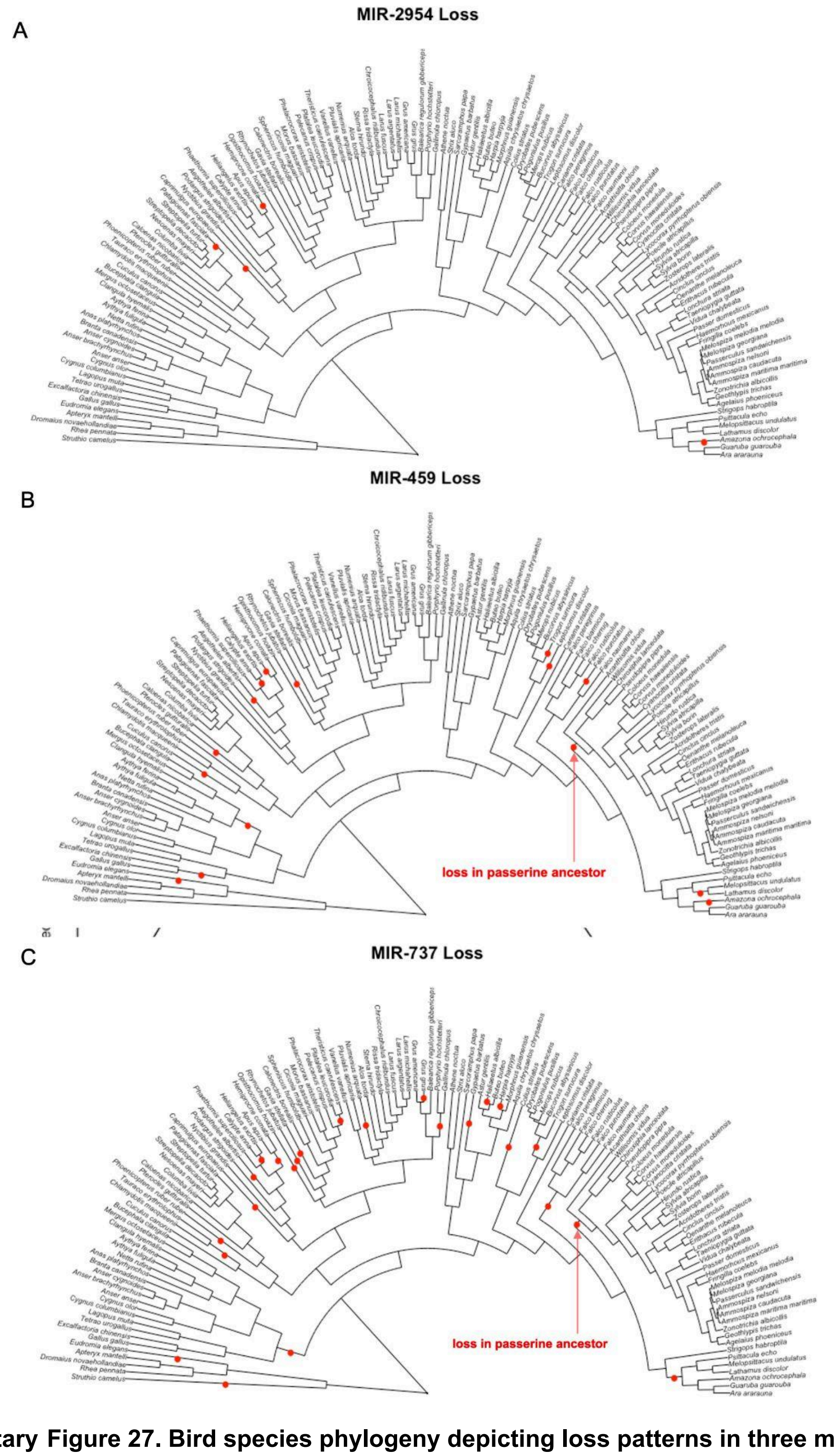


**Supplementary Figure 27. Bird species phylogeny depicting loss patterns in three miRNA gene families**. Losses of miRNA gene families inferred through reconstruction using Dollo parsimony are shown as red circles. (A) MIR-2954, associated with song-response in zebra finch (Lin et al, 2014) is lost four times in birds, but retained in all passerine species. (B) MIR-459 and (C) MIR-737 are both lost in the passerine ancestor and are absent throughout Passeriformes, and also show widespread losses across bird species generally.


1. Ranallo-Benavidez, T.R., Jaron, K.S., and Schatz, M.C. (2020). GenomeScope 2.0 and Smudgeplot for reference-free profiling of polyploid genomes. Nat. Commun. *11*, 1432.

2. Bernt, M., Donath, A., Jühling, F., Externbrink, F., Florentz, C., Fritzsch, G., Pütz, J., Middendorf, M., and Stadler, P.F. (2013). MITOS: improved de novo metazoan mitochondrial genome annotation. Mol. Phylogenet. Evol. *69*, 313–319.

3. Rautiainen, M., Nurk, S., Walenz, B.P., Logsdon, G.A., Porubsky, D., Rhie, A., Eichler, E.E., Phillippy, A.M., and Koren, S. (2023). Telomere-to-telomere assembly of diploid chromosomes with Verkko. Nat. Biotechnol. *41*, 1474–1482.

4. Rautiainen, M. (2024). Ribotin: automated assembly and phasing of rDNA morphs. Bioinformatics *40*, btae124.

5. Vollger, M.R., Kerpedjiev, P., Phillippy, A.M., and Eichler, E.E. (2022). StainedGlass: interactive visualization of massive tandem repeat structures with identity heatmaps. Bioinformatics *38*, 2049–2051.

6. Pigozzi, M.I., and Solari, A.J. (1998). Germ cell restriction and regular transmission of an accessory chromosome that mimics a sex body in the zebra finch, Taeniopygia guttata. Chromosome Res. *6*, 105–113.

7. McTavish, E.J., Hinchliff, C.E., Allman, J.F., Brown, J.W., Cranston, K.A., Holder, M.T., Rees, J.A., and Smith, S.A. (2015). Phylesystem: a git-based data store for community-curated phylogenetic estimates. Bioinformatics *31*, 2794–2800.

8. Benson, G. (1999). Tandem repeats finder: a program to analyze DNA sequences. Nucleic Acids Res. *27*, 573–580.

9. Takki, O., Komissarov, A., Kulak, M., and Galkina, S. (2022). Identification of centromere-specific repeats in the zebra finch genome. Cytogenet. Genome Res. *162*, 55–63.
